# Anoctamin-2-specific T Cells Link Epstein-Barr Virus to Multiple Sclerosis

**DOI:** 10.64898/2025.12.04.692416

**Authors:** Olivia G. Thomas, Urszula Rykaczewska, Marina Galešić, Rianne T. M. van der Burgt, Nils Hallén, Filippo Ferro, Mattias Bronge, Zoe Marti, Yue Li, Alexandra Hill Riqué, Jianing Lin, Aleksa Krstic, Alicja Gromadzka, András Levente Szonder, Chiara Sorini, María Reina-Campos, Ting Sun, Leslie A. Rubio Rodríguez-Kirby, Özge Dumral, Rasmus Berglund, Majid Pahlevan Kakhki, Milena Z. Adzemovic, Manuel Zeitelhofer, Birce Akpinar, Katarina Tengvall, Ola B. Nilsson, Erik Holmgren, Chiara Starvaggi Cucuzza, Klara Asplund Högelin, Guro Gafvelin, Katharina Fink, Gonçalo Castelo-Branco, Maria Needhamsen, Mohsen Khademi, Fredrik Piehl, Torbjörn Gräslund, Lars Alfredsson, Harald Lund, Per Uhlén, Ingrid Kockum, Roland Martin, Maja Jagodic, Hans Grönlund, André Ortlieb Guerreiro-Cacais, Tomas Olsson

## Abstract

Multiple sclerosis (MS) occurs when the central nervous system (CNS) is damaged by misguided adaptive immune responses, likely caused by a combination of environmental factors in genetically susceptible individuals. A known prerequisite for disease is Epstein-Barr virus (EBV) infection, and previous studies have demonstrated elevated Epstein-Barr virus nuclear antigen 1 (EBNA1) antibodies which cross-react with the calcium-activated chloride channel anoctamin-2 (ANO2) in persons with MS (pwMS). ANO2-reactive antibodies have been associated with greater neuroaxonal damage in MS, although their exact effector function is still uncertain. Here, we demonstrate that ANO2 is also the target of IFNγ-producing CD4^+^ T cells, which are more frequent in untreated and natalizumab-treated pwMS compared to control individuals. Immunisation of SJL/J mice with either ANO2 or EBNA1 elicited cross-reactive CD4^+^ T cell and antibody responses *in vivo.* Pre-immunisation of young mice with ANO2 worsened proteolipid protein (PLP)-induced experimental autoimmune encephalomyelitis (EAE), which in older mice included atypical clinical phenotype, immune infiltration into the brain and reduced survival. EAE exacerbation was recapitulated with the adoptive co-transfer of ANO2 and myelin antigen-specific CD4^+^ T cells, and ANO2-specific T cells alone could induce the cell death of ANO2-expressing glial cells *in vitro*. T cell clones with cross-reactivity to both EBNA1 and ANO2 antigens could be isolated from natalizumab-treated pwMS. Single cell sequencing of EBNA1 and ANO2-specific T cell receptors (TCR) from four pwMS revealed a significant overlap between their antigen-specific expanded TCR repertoires within donors and transcriptomic analysis showed cross-reactive T cells to have predominantly activated and cytotoxic phenotypes. In summary, we report the first mechanistic evidence that EBNA1 CD4^+^ T cells can target the MS-associated autoantigen ANO2, thereby establishing a link between EBV infection and development of autoimmune neuroinflammatory disease.

## Introduction

Multiple sclerosis (MS) is a common cause of neurological disability in young adults in developed countries, with an estimated 2.8 million people diagnosed worldwide and rising^1^. MS features widespread inflammatory lesions in the central nervous system (CNS) and usually presents clinically with a relapsing-remitting phenotype, often preceded by a preclinical phase with vague symptoms that may last for several years^2^. Both environmental and stochastic events are thought to work in combination to trigger disease in genetically susceptible individuals^3^, with the strongest genetic risk factor being *HLA-DRB1*15:01*^4^.

Epstein-Barr virus (EBV)^5^ is the common cause of infectious mononucleosis (IM)^6^ and is likely a prerequisite for MS development^5,7,8^, but there is still no accepted mechanism for how the virus may contribute to disease. However, mounting evidence suggests that molecular mimicry between immune responses targeting EBV and MS-associated autoantigens may be responsible^9^, where the established association of elevated Epstein-Barr nuclear antigen 1 (EBNA1)-reactive antibodies with MS^10–12^ has led to this antigen being the focus of investigation. Detailed analyses of this association showed that some regions of EBNA1 are more strongly linked to MS than others, with antibodies against the amino acid (aa) fragments covering the 385-450 region of EBNA1 being most strongly linked to MS and also positively associating with *HLA-DRB1*15:01* carriage^11,13,14^. Whilst the functional role of EBNA1-reactive antibodies in MS remains to be resolved, recent evidence of sequence homology and antibody cross-reactivity between EBNA1 and the autoantigens anoctamin-2 (ANO2)^15^, glial cell adhesion molecule (GlialCAM)^16,17^ and ɑ-crystallin B (CRYAB)^18^ suggests that anti-EBNA1 IgG is associated with damage of self-tissues and may contribute to MS pathology. Furthermore, observations from a pre-MS cohort showed that anti-ANO2 IgG are associated with 26% higher levels of serum neurofilament light (sNFL), a marker of ongoing neuroaxonal injury, supporting their functional relevance in disease^19^.

ANO2 is a calcium-activated chloride channel which may be involved in sensory transduction and neuronal excitation^20–22^. Staining of ANO2 in neuronal cell bodies is increased in and around MS lesions^23^ and carriage of this antibody specificity associates with drastically higher odds ratios (ORs) for MS in conjunction with different HLA risk alleles^15^. Together, these findings support a role for ANO2 autoreactivity in MS pathogenesis, at least in a subgroup of individuals.

However, cross-reactive autoantibodies are unable to fully account for all clinical and immunological features of MS, and evidence from both humans and experimental autoimmune encephalomyelitis (EAE) – the rodent model of MS – supports a key role for CD4^+^ T cells in its pathogenesis. For example, EAE can be induced by adoptive transfer of myelin-reactive CD4^+^ T cells but not by autoantibodies alone, and mice carrying human HLA class II alleles are particularly susceptible to demyelinating disease^24–28^. The presence of CD4^+^ T cells in early inflammatory brain lesions, correlation of T_H_1 cells in peripheral blood with relapses and the genetic association with *HLA-DRB1*15:01* all further support involvement of CD4^+^ T cells in MS pathology^4,29–31^.

EBV persists in the host predominantly in latently infected B cells, and lifelong infection primes T cell responses which have a role in suppressing viral reactivation. T cell responses to EBNA1 are elevated and respond to a broader range of epitopes in MS suggesting that CD4^+^ T cells specific for EBNA1, as well as antibodies, are similarly prominent in disease^18,32,33^. An important outstanding question, however, is whether EBNA1 responses are actively involved in autoimmune neuroinflammation or merely represent epiphenomena. The former notion is supported by a study showing that EBNA1-reactive CD4^+^ T cells more frequently recognise myelin antigens than other self-proteins and display a T_H_1 phenotype with co-expression of interferon-γ (IFNγ) and interleukin-2 (IL-2)^34^. More recently EBNA1-specific T cell clones (TCC) from pwMS were shown to recognise myelin oligodendrocyte glycoprotein (MOG)^35^. While providing only suggestive evidence, these findings nevertheless provide an indication that EBV-specific T cells can target CNS tissue. Importantly, studies of autoantibodies in other autoimmune diseases such as Addison’s disease and type 1 diabetes mellitus have demonstrated these autoantibodies to be markers of pathogenic T cell responses, which are the effector cells actually responsible for tissue damage^36,37^. Therefore, we hypothesised that a similar underlying mechanism may exist in MS.

Given the link between ANO2-specific antibodies and MS and also cross-reactive antibodies between EBNA1 and ANO2, we hypothesised that ANO2-specific T cells may also be present in pwMS and may cross-react with EBNA1 via molecular mimicry. We therefore set out to explore this in the present study using both human material from pwMS and animal models. We characterised T cell responses to ANO2 in controls and pwMS, and we investigated ANO2 autoimmunity and cross-reactivity of these responses in EAE by extensively characterizing the disease course and CNS immune infiltration. Finally, using single cell methods to profile antigen-specific CD4^+^ T cells from pwMS, we shed light on the transcriptional phenotype of T cells responding to EBNA1, ANO2 and CRYAB, and investigate the overlap between TCR repertoires responding to these antigens. Through these experiments, we aimed to understand what proportion of these T cells could have been originally primed *in vivo* by exposure to EBNA1 and EBV persistence in the host.

## Materials and Methods

### Study Participants

The study was approved by Regionala Etikprövningsnämnden in Stockholm (no. 04-252/1-4 and no. 2009/2107-3112, with last revision 2024-07377-02) and the Cantonal Ethics Committee of Zürich (EC-No. of the research project 2013-0001, approved on 5^th^ June 2013 and EC-No. of the ERC 2014-0699, approved on 27^th^ February 2015) and was conducted in accordance with the Declaration of Helsinki.

Study participant groups and demographics are presented in Table 1. All participants provided written, informed consent.

**Table 1.** Cohort Demographics.

|  | Antibody Cohort |  | T cell Cohorts |  |  |  |  |  |  |  |  |
| --- | --- | --- | --- | --- | --- | --- | --- | --- | --- | --- | --- |
|  |  |  | Cohort 1 |  | Cohort 2 |  |  | Validation cohort |  |  |  |
|  | MS | Controls | HC | MS-Nat | HC | OND | MS-Un | MS-Nat | MS-Un | HC | OND |
| **Size (n)** | 713 | 722 | 28 | 61 | 21 | 19 | 31 | 59 | 6 | 19 | 20 |
| **Age (years)** | 39.9 ± 10.2<br>(17-67) | 42.6 ± 10.4<br>(17-68) | 33.9 ± 8.2<br>(20-49) | 36.3 ± 9.7<br>(17-61) | 33.3 ± 6.8<br>(25-49) | 29.3 ± 10.5<br>(18-57) | 34.2 ± 9.7<br>(19-58) | 36.2 ± 10.0<br>(17-61) | 33.4 ± 4.3<br>(29-38) | 33.6 ± 6.7<br>(24-48) | 28.4 ± 9.7<br>(18-57) |
| **Sex (female)** | 70.3 % | 74.9 % | 44<br>(72.1%) | 21<br>(75.0%) | 15<br>(75.0%) | 9<br>(47.4%) | 23<br>(74.2%) | 71.2 % | 50.0 % | 63.2 % | 45.0 % |
| **HLA-**<br>**DRB1\*15:01**<br>**(n)** | 360/649<br>(55.5%) | 196/673<br>(29.1%) | 28 / 56<br>(50.0%) | 7 / 22<br>(31.8%) | 3 / 12<br>(25%) | 10 / 13*<br>(76.9 %) | 11 / 23<br>(47.8%) | 27 / 54<br>(50 %) | - | 2 / 8<br>(25 %) | 9 / 11<br>(81.8 %) |
| **EDSS (score)** | 1.75±1.5 | - | - | 2.1 ± 1.5<br>(0.0–7.5) | - | - | 1.5 ± 1.3<br>(0.0–6.5) | 2.3 ± 1.7<br>(0.0–7.5) | 1 ± 0.6<br>(0.0–1.5) | - | - |
| **Disease**<br>**Duration**<br>**(years)** | 5.7 ± 6.1<br>(0-33) | - | - | 8.9 ± 5.4<br>(1-20) | - | - | 2.1 ± 3.1<br>(0.0–12.7) | 8.5 ± 5.4<br>(1-20) | 0.3 ± 0.5<br>(0-1.2) | - | - |
| **History of IM**<br>**(n, +/-/NA)** | 81/407/225 | 53/518/151 | - | - | - | - | - | - | - | - | - |

± denotes SD. Range or percentage of whole in brackets. n / N denotes the number of individuals for which data was available if it was not the whole cohort.
\*DR15-status in two cases based on the high linkage disequilibrium with DQB1\*06:02. RRMS: Relapsing-remitting MS. HCs: Healthy Controls. Nat: Natalizumab-treated. Un: Untreated. EDSS: Expanded Disability Status Scale.

For the T cell studies in MS, a cohort consisting of pwMS without ongoing disease modifying treatment (MS-Un), or with natalizumab treatment (MS-Nat), age- and sex matched healthy controls (HC) and persons with other neurological disease (OND, n=20, cases of narcolepsy type 1 or 2, and idiopathic hypersomnia) was collected (Table 1). Peripheral blood mononuclear cells (PBMC) were collected from venous blood samples using density gradient (Ficoll) separation and cryopreserved at –150°C.

For antibody analysis, plasma samples from the Swedish nation-wide epidemiological incident case-control cohort (EIMS)^38^ totaling 713 pwMS and 722 Controls (Table 1). The controls were population-based controls matched to cases on sex, age, and geographic region. HLA-data were available for the cohort from previous studies at our institution and IM history was self-reported in a questionnaire answered at the time of consent and sampling ^15^.

### Sample preparation

Blood was collected from participants in EDTA blood tubes and processed within 2 hours of sampling by Ficoll density gradient centrifugation using SepMate tubes (StemCell Technologies) as per the manufacturer’s protocol. The PBMC fraction was then incubated for 5 minutes with ACK lysis buffer (Sigma-Aldrich) to lyse the remaining red blood cells before washing with RPMI-1640 medium (Sigma) and re-suspending in freezing medium (10% DMSO, 45% RPMI-1640, 45% heat-inactivated foetal calf serum (FCS)). PBMC were then frozen at -1°C/minute and transferred to -150°C within 48 hours for long-term storage.

Plasma samples were obtained from the EIMS cohort and were processed as previously described^38^.

### Production of recombinant antigens and antigen-coupled beads

Antigen beads were developed and tested in collaboration with NEOGAP Therapeutics AB (Stockholm, Sweden). Briefly, recombinant ANO2_79-168_, ANO2_1-275_, ANO2_409-525_, ANO2_750-1003_, EBNA1_1-120_, EBNA1_380-641_ and albumin binding domain (ABD)^39,40^ proteins were produced as previously described^18,40,41^. ABD served as an irrelevant protein control^18,42^. Custom genes covering regions of ANO2 (Uniprot identifier: Q9NQ90) and EBNA1 (B95.8 strain, Uniprot identifier: P03211) including Bsa I sites were ordered (Eurofins Genomics) and subcloned into a modified pET28 vector backbone containing an 8xHIS tag. Vectors containing gene constructs were then transformed into competent BL21-AI *E.coli* (Thermofisher Scientific) and grown as previously described^18^. Proteins were purified, analysed by SDS-PAGE using the SeeBlue plus2 protein ladder (Thermofisher Scientific) (Figure S1A) and concentration determined by Nanodrop (Thermofisher Scientific) as previously described^18^.

Recombinant proteins were coupled to paramagnetic Dynabeads MyOne beads (Thermofisher Scientific) via amine coupling to carboxylic acid groups on the bead surface as previously described^40^. NaOH concentration was determined for each antigen as previously described^41^ and coupling efficiency was determined^40^ (Figure S1B).

### FluoroSpot

Cryopreserved PBMC were thawed and counted before adding to IFNγ/IL-17A/IL-22 FluoroSpots (Mabtech) as previously described^18^. PBMC samples with <80% viability after thawing were excluded from further experiments and viability did not differ significantly between groups (Figure S1C). 2.5×10^5^ PBMC were added per well alongside 3×10^6^ antigen-coupled beads (1.25×10^5^ PBMC added for anti-CD3 (aCD3) control wells). For epitope mapping FluoroSpots, PBMC were stimulated with 2μg/ml peptide (Peptides & Elephants) in human IFNγ/TNFɑ/Granzyme B FluoroSpot (Mabtech). All test conditions were performed in duplicate and samples from ≥2 cohort groups were included on each plate. FluoroSpot plates for Cohort 2 (Figure 1B) were read using a iSPOT Spectrum device (Advanced Imaging Devices GmbH) which used AID ELISpot Reader v.7 software (Advanced Imaging Devices GmbH). Cohort 1 plates (Figure 1B) and all other plates in the current study were read using an IRIS plate reader (Mabtech, Sweden) which uses the software SpotReader v.1.1.9.

**Figure 1.**
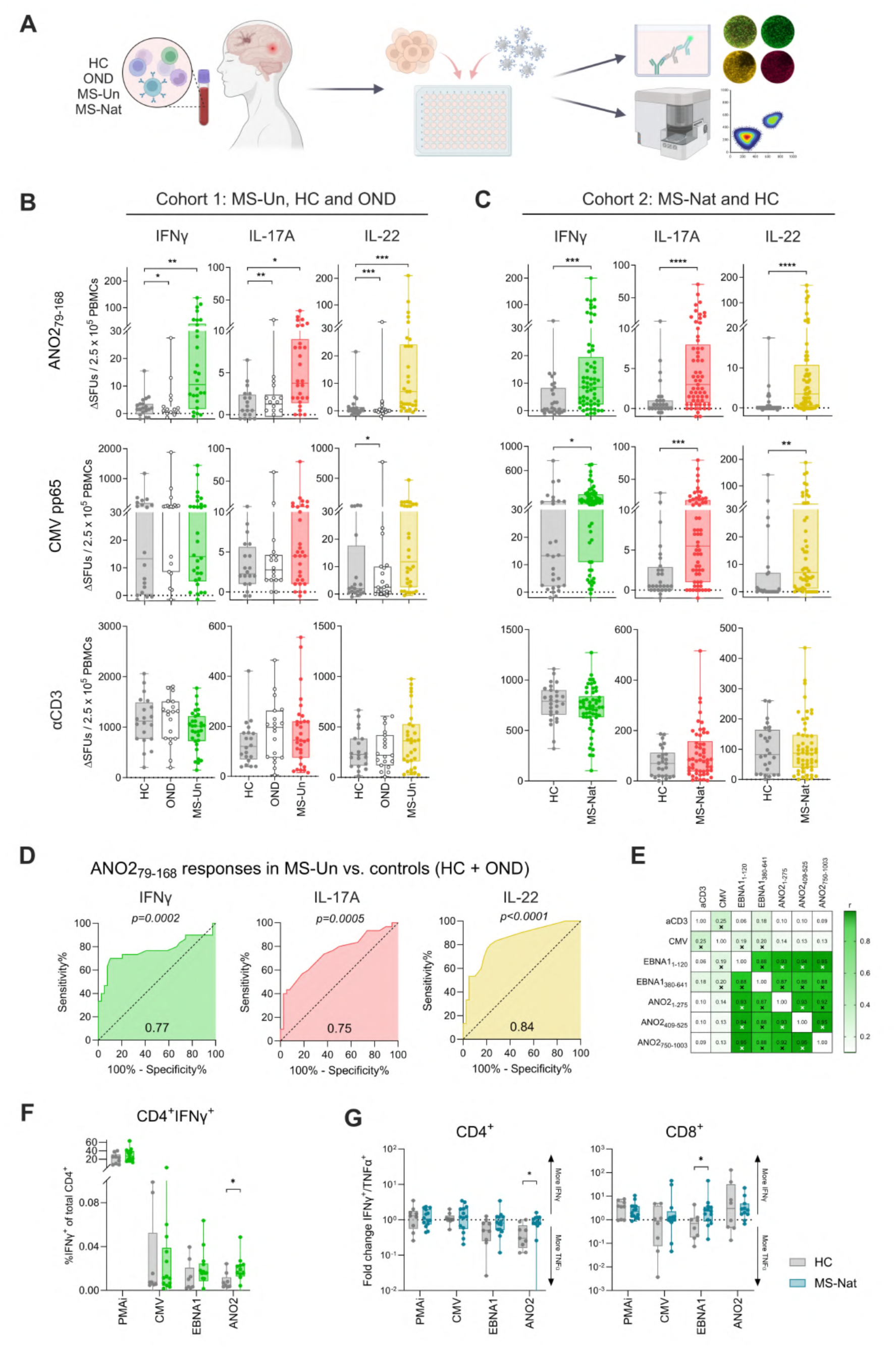
ANO2 is a target of T cell responses in MS. **A.** A schematic for investigating antigen-specific T cell responses in MS. Briefly, PBMC were isolated from donors and cryopreserved. After thawing, PBMC were stimulated with antigen-coupled beads or control stimuli in either FluoroSpot plates (IFNγ/IL-17A/IL-22) or 96-well tissue culture plates. Fluorospot plates were developed after 44 hours. PBMC stimulated in tissue culture plates were incubated for 16 hours and intracellular cytokine staining was performed before data collection on a spectral flow cytometer. Created with BioRender.com. **B.** Number of IFNγ (left), IL-17A (middle) and IL-22 (right) SFU after stimulation in a FluoroSpot assay with ANO2_79-168_, CMV pp65, or anti-CD3 in cohort 1: MS-Un (n=30), HC (n=20) and OND (n=19). Boxes represent median ± IQR and statistical significance was calculated with the Kruskall-Wallis test with Dunn’s multiple comparisons test. P-values are indicated where significant. **C.** Number of IFNγ (left), IL-17A (middle) and IL-22 (right) SFU after stimulation with ANO2_79-168_, CMV pp65, or anti-CD3 in cohort 2: MS-Nat (n=61) and HC (n=28). Boxes represent median ± IQR, statistical significance was calculated with the Mann-Whitney test and p-values indicated where significant. **D.** Receiver operating characteristic (ROC) based on IFNγ (left), IL-17A (middle) and IL-22 (right) responses to ANO2_79-168_ in control (HC + OND, n=20+19 respectively) and MS-Un donors (n=30). Area under curve indicated at the bottom of graphs and p-values indicated above graphs. **E.** Correlation matrix of IFNγ T cell responses to aCD3, CMV pp65, EBNA1_1-120_, EBNA1_380-641_, ANO2_1-275_, ANO2_409-525_, and ANO2_750-1003_ in individuals across HC (n=22), OND (n=20), MS-Un (n=6) and MS-Nat (n=68) groups by FluoroSpot. Spot-forming units (SFU). Linear regression r values are indicated significant linear regression values are indicated by crosses. **F.** Spectral flow cytometry analysis of autoreactive T cells using intracellular cytokine staining after autoantigen stimulation. CD4^+^IFNγ^+^ responses to autoantigens in MS-Nat (n=12, green bars) and HC (n=9, grey bars). Mann-Whitney statistical test. **G.** Ratios of IFNγ and TNFɑ-producing CD4^+^ T cells (left) and CD8+ T cells (right) after CMV pp65, EBNA1 and ANO2 stimulation. Group differences were evaluated using the Mann–Whitney U test. Untreated MS (MS-Un), natalizumab-treated MS (MS-Nat), healthy controls (HC), other neurological disease (OND), spot-forming units (SFU). \**p* < 0.05; \*\**p* < 0.01; \*\*\**p* < 0.001; \*\*\*\**p* < 0.0001. Created with https://BioRender.com.

Plates were read using an IRIS plate reader (Mabtech), and each individuals’ response to negative control (NC) beads was subtracted from the spot count in the antigen bead-stimulated wells.

For HLA-blocking FluoroSpots, the protocol was the same with the addition of 10μg/ml of anti-HLA-A/B/C, anti-HLA-DR or isotype control antibodies (BioLegend) to wells. Tests were excluded from analysis if unresponsive in the antigen-stimulated isotype control conditions (defined as <5 spot forming units (SFU) for IFNγ, IL-17A and IL-22).

FluoroSpots to assess antigen-specific responses in T cell clones (TCC) and bulk T cell (TCB) lines were performed in duplicate wells of human IFNγ/TNFɑ/Granzyme B or human IFNγ FluoroSpot plates (Mabtech) at approximately 5×10^4^ cells/well with 1×10^5^ HLA-DRB1*15:01-matched or mismatched irradiated PBMC (45 Grays) added to wells to act as antigen-presenting cells (APC). For some experiments, DR2a- (*HLA-DRB5*01:01*) or DR2b- (*HLA-DRB1*15:01*) expressing bare lymphocyte syndrome (BLS) cells (generated by Bill Kwok, Benaroya Research Institute, Seattle) were irradiated (100 Grays) and used as APCs. Peptides were added at 2μg/ml and antigen-coupled beads at a bead:cell ratio of 10:1.

For epitope mapping of TCC in Stockholm, TCC were first screened for reactivity to antigen-coupled ANO2_1-275_ beads, EBNA1_380-641_ beads, ANO2_1-275_ peptides or EBNA1_380-641_ peptides (Peptides & Elephants). TCC which responded to either ANO2, EBNA1 or both stimulations were then tested by IFNγ/IL22/IL17A FluoroSpot (Mabtech) for responses to overlapping ANO2_1-275_ and EBNA1_380-641_ peptide libraries (15-mers overlapping by 11aa) at final concentration of 2μg/ml and anti-CD3 as a positive control in the presence of 5×10^4^ HLA-DRB1*15:01+ irradiated PBMC (45 Grays).

For EBNA1-specific TCB lines from 2113HU and 2092PH in Zurich, CD45RA-PBMC were stimulated with EBNA1_380-641_-beads at a 5:1 bead-to-cell ratio for 7-8 days. Proliferating T cells were sorted by flow cytometry as CD3^+^CD4^+^CD8^-^CFSE_DIM_ and subsequently expanded. The cells were screened for ANO2 reactivity using IFN*γ* ELISpot assays (Mabtech, 3420-2APT-2) at a seeding density of 3.5 × 10^4^ cells/well in presence of irradiated (300 Grays) BLS cells expressing DR2a (HLA-DRB5*01:01) or DR2b (HLA-DRB1*15:01) at 5 × 10^4^ cells/well. Six ANO2 pools were generated by pooling 11 (pools 1-5) and 12 (pool 6) 20-mer peptides with overlaps of 5 amino acids. Cells were stimulated with each pool at a final concentration of 7.5μM per single peptide. Positive controls were an overlapping EBNA1_380-641_ peptide pool (20-mers overlapping by 5 aa, n=18, final concentration 1.35μM per peptide) and anti-CD2/3/28 beads (Miltenyi, 130-091-441, bead-to-cell ratio of 1:2). Cells were incubated in X-VIVO® 15 serum-free media (Lonza, 02-053Q) for 44h, and spots were developed according to the manufacturer’s protocol.

### Spectral flow cytometry

Cryopreserved PBMC from representative donors (natalizumab-treated pwMS (MS-Nat), n=12, healthy controls (HC), n=9) were stimulated and analysed by intracellular cytokine staining and spectral flow cytometry as previously described.^18^ Stimuli were beads at a bead:cell ratio of 10:1 and phorbol 12-myristate 13-acetate and ionomycin (PMA/I, Invitrogen) was used as a positive control stimulus. Antibody staining panel in Table S2 and samples were acquired on an Aurora 4L instrument (Cytek). Spectral overlap was calculated using SpectroFlow software (Cytek) and data was analysed in FlowJo v10 (BD Biosciences) and Cytobank (Beckman Coulter). Flow cytometry gating strategy is presented in Figure S5.

### Autoantibody suspension bead array

Assays to investigate antibody responses were performed by suspension bead array as previously described^15,18^. Proteins and peptides from EBNA1, ANO2 and CRYAB were selected based on previously reported immunoreactivity, association with MS and/or homology. Briefly, peptides and proteins (full list in Table S2) were coupled to biotin and an amino acid spacer (PEPscreen, Sigma-Aldrich) and coupled to NeutrAvidin beads (Luminex Corporation). To deplete tag-binding antibodies and to remove background, blinded plasma samples were diluted 1:50 in phosphate-buffered saline (PBS) supplemented with 5% bovine serum albumin (BSA), 0.1% Tween 20, 6xHis-ABD (160μg/ml) and NeutrAvidin (10μg/ml) before incubation for 1 hour. After 1hour, diluted plasma samples were incubated for a further 2 hours in the suspension bead array. Samples were fixed with 0.2% formaldehyde for 10 minutes and incubated for 30 minutes with secondary antibody (phycoerythrin F(ab’)2-goat anti-human IgG Fc secondary antibody, Invitrogen). Samples were analysed using a FLEXMAP 3D (Luminex Corporation). Mean fluorescence intensity (MFI) for each plasma sample was adjusted based on the assumption that 33% of antigens would be negative, and the 33^rd^ percentile MFI (_Per33_MFI) was subtracted from each response. A threshold for positive response was created on the basis of the whole cohort’s response to the tested negative peptides; negative controls were 6xHis-ABP (purification tag), rabbit anti-human IgG (Jackson Laboratories) and buffer only, and the threshold at 99.9^th^ percentile response was _Per33_MFI of 2484.85). Samples were run across four different plates.

### Expansion and flow cytometry-assisted sorting of antigen-specific T cells for T cell cloning and single cell sequencing

To isolate T cells which responded to ANO2 and EBNA1 for T cell cloning, single cell sequencing and epitope mapping, CD45RA-negative cells were isolated using CD45RA microbeads (Miltenyi) from cryopreserved MS-Nat PBMC according to the manufacturer’s instructions. CD45RA-negative cells were then stained with CFSE (Thermofisher Scientific) and stimulated with antigen beads at a ratio of 10:1 or peptide pools at concentration 2μg/ml peptide (Peptides & Elephants), and incubated for 8 days at 37°C, 5% CO_2_. After 8 days, cells were stained with Live/Dead Aqua viability dye (Thermofisher Scientific), CD3 APC-Cy7 and CD4 PerCP-Cy5.5 antibodies (BioLegend) and the LiveCD3^+^CD4^+^CFSE^DIM^ population sorted using a SH800S Cell Sorter (Sony).

For T cell cloning, single LiveCD3^+^CD4^+^CFSE^DIM^ T cells were sorted into wells of 96-well U-bottom plates containing 2×10^5^ irradiated allogeneic PBMC (45 Grays) per well in T cell media (IMDM 5% human serum (Merck), 100U/ml penicillin, 100μg/ml streptomycin, 2mM L-glutamine) containing PHA (Thermofisher Scientific) and 20U/ml human IL-2 (Peprotech) and expanded for 2 weeks. After 2-4 weeks, visible colonies were transferred to separate plates and rested in media without IL-2 supplementation for 5 days prior to antigen specificity testing.

For single cell TCR sequencing, after 8 days stimulated CD45RA-negative cells were washed once with FACS buffer (PBS 1% FCS) and barcoded by staining with anti-β2-microglobulin barcoding TotalSeq-C antibodies (BioLegend, Table S3) for 20 minutes at 4°C to identify the original stimulation. After 2 washes with FACS buffer, cells were stained with a surface receptor master mix of antibodies containing CD3 APC-Cy7 antibody (Biolegend), CD4 PerCP-Cy5.5 antibody (BioLegend) and TotalSeq-C antibodies (Table S4A) and incubated for 30 minutes at 4°C. After 2 washes with FACS buffer, cells were re-suspended in FACS buffer and the LiveCD3^+^CD4^+^CFSE^DIM^ population bulk sorted into cold RPMI in polypropylene FACS tubes (Corning) and counted 1:1 with Trypan blue (Gibco) before processing for single cell sequencing (Table S4B). The sorted cells were loaded onto Chromium Next GEM Chip K (10X Genomics) on Chromium Controller (10X Genomics) for cDNA sequencing and barcoding. Single cell library preparation was performed using the Chromium Next GEM Single Cell 5’ Reagent Kit v2 (Dual Index) with feature barcode technology for immune receptor mapping (10X Genomics) according to the manufacturer’s instructions (single cell sequencing hashtag antibodies in Table S4B). Stimulated cells were FACS sorted separately and T cells responding to antigens were loaded onto chips (Table S4B). Library quality control was determined using BioAnalyzer High Sensitivity DNA Analysis (Agilent) and KAPA Universal Library Quantification (Roche) kits according to the manufacturers’ instructions and sequenced on a NovaSeq Xplus (Illumina).

### Experimental animals

All animal experiments were performed at Karolinska Institutet and are approved and performed following the Swedish National Board on research animal ethics and the European Community Council Directive (86/609/EEC) and the local ethics committee of Stockholm North under the ethical permits 9328-2019, 23561-22 and 7029/2020. Male and female C57BL/6Ntac (Taconic), male NOD (Taconic), female SJL/J (Janvier), male and female SJL/J x C57BL/6Ntac F1 and SJL/J x C57BL/6-GFP (Jackson Laboratory Strain 004353) F1 mice were kept at the Comparative Medicine Department at Karolinska University Hospital, Sweden. Animals were maintained in a pathogen-free and climate-controlled environment with regulated 12-hour light/dark cycles. All mice used for experiments were adults between 2 and 4 months of age, with the exception of adult SJL/J which were between 6 and 7 months of age, weighing 20–35 g, and with access to chow and water ad libitum.

### Mouse antigen recall experiments

For immune priming and assessment of T cell cross-reactivity, animals were immunised subcutaneously at the tail base with ABD (irrelevant protein control), ANO2_1-275_ or EBNA1_380-641_ in 100μl of Freund’s complete adjuvant (CFA, Chondrex) containing 200μg of heat-killed Mycobacterium tuberculosis (MTB) per mouse. Antigen amounts per mouse varied between experiments (50-100μg) but were given at comparable doses within each experiment for all antigens. Unimmunised animals as well as CFA immunised animals were used as controls. Day 9-11 after immunisation, animals were sacrificed and spleen and draining (inguinal) lymph nodes were taken for assessment of antigen-specific responses. Lymph nodes and spleens were mechanically dissociated, red blood cells were lysed with ACK buffer (Sigma) or Hybri-Max Red Blood Cell lysis buffer (Sigma) and cells were spun and resuspended in RPMI-1640 complemented with 10% foetal bovine serum, 1% l-glutamine, 1% penicillin-streptomycin, 1% pyruvic acid, 1% MEM non-essential amino acids, 10mM HEPES (all from Sigma-Aldrich), and 50mM 2-mercaptoethanol (Gibco-BRL). Cells were plated at 0.2 × 10^6^ cells per well in U-bottom plates and restimulated with the immunisation antigen and a panel of control as well as putative cross-reactive antigens, all bead-conjugated: ABD, ANO2_1-275_, EBNA1_1-120_, EBNA1_380-641_, SNAP91, RASGRP2, SNCB, GlialCAM_262-418_. After 96 hours, cells were replated into V-bottom plates and restimulated for 5 hours with phorbol 12-myristate 13-acetate (PMA) (50 ng/ml; Sigma-Aldrich), ionomycin (1μg/ml; Sigma-Aldrich), and brefeldin A (GolgiPlug) (1μl/ml; BD Biosciences). Samples were subsequently stained with antibodies according to Table S3A and gating strategy in Figure S10A). LIVE/DEAD Fixable Near-IR or Yellow dead cell exclusion dyes (L34976 or L34959, Invitrogen) were used to exclude dead cells. Intracellular/intranuclear staining was performed after permeabilisation using a fixation/permeabilisation kit (eBioscience). Cells were acquired using a LSR Fortessa flow cytometer (BD) and analysed using Kaluza software (Beckman Coulter).

For priming and restimulation experiments with proteins, peptide pools and individual peptides, animals were immunised with ANO2_1-275_ or EBNA1_380-641_ proteins, an ANO2_1-275_ overlapping peptide pool of 15-mers (overlapping by 11aa) and covering the aa1-275 region, an EBNA1 overlapping peptide pool covering the aa380-641 region similar to above, the ANO2 peptide 35 (aa137-151) containing the EBNA1 mimicry region, the mouse counterpart of this minimal peptide, the EBNA1 peptide 13 (aa428-442) containing the ANO2 mimicry region and the HCMV IE1_162-173_ as an irrelevant control, all emulsified in CFA as described above. Animals were immunised, lymph nodes and spleens were processed and cells restimulated *in vitro* against the original immunogens as well as to all the other immunogens described above, all at 20μg/ml. For the mapping of responses against EBNA1, cells were additionally restimulated against smaller pools of 15-mer peptides (overlapping by 11aa, 10-11 peptides per pool) covering the aa380-641 region. After staining, samples were acquired using an Aurora flow cytometer (Cytek) and analysed using FlowJo (BD) software.

### Mouse immunisations and EAE assessment

For the induction and clinical evaluation of EAE in response to different antigens, we used a protocol adapted from Lanz *et al.*^16^. In brief, mice were immunised subcutaneously with 100μl of emulsion divided into two injections of 50μl on each flank containing ABD (irrelevant protein control), ANO2_1-275_, EBNA1_380-641_ or pools of overlapping peptides, GlialCAM_262-418_, or the specific cross-reactive peptide sequences EBNA1_428-442_ or ANO2_137-151_. Antigens were emulsified in Freund’s incomplete adjuvant (IFA; Sigma) containing 65-85μg CpG ODN 1826 (Invivogen), a TLR9 agonist. Amounts of proteins/peptides given per animal varied between 70μg and 200μg per animal in different experiments but were kept even between different groups of animals within the same experiment. After three weeks, mice were immunised subcutaneously on the tail base with 100μg PLP_139-151_ in 100μl CFA containing 200μg of heat-killed Mycobacterium tuberculosis (MTB) per mouse and their disease course followed for up to 60 days (Figure 2B). Pertussis injections were tried in initial experiments but proven only to hasten the onset and increase the general severity, without greatly modifying the disease course, and were not used for subsequent experiments. Mice were scored and weighed daily from day 10 after immunisation for any abnormalities in tail tonus and gait. The typical clinical score was graded as follows: 0, no clinical signs of EAE; 1, tail weakness or tail paralysis; 2, hind leg paraparesis or hemiparesis; 3, hind leg paralysis or hemiparalysis; 4, tetraplegia or moribund; 5, death. For atypical symptoms, grading was done as follows: 1, tail tonus but slight wobble; 2, tail tonus but strong wobble, head tilting and/or poor balance; 3, severe head tilting, walking along the edges of the cage, spinning when picked up and/or front leg paralysis whilst retaining hind limb and tail tonus; 4, spasms, inability to stand upright. Disease incidence did not consistently differ between control and MS-related antigens for any of the conditions compared (except for aged mice, see below) and thus the average clinical scores for each day only included mice who showed weight loss and displayed signs of EAE. For aged mice (Figure 3), a clear pattern of increased incidence was observed in the ANO2-preimmunised group and unaffected mice were included in the score analysis, as stated in the figure legend. Mice were euthanised if presenting with > 25% weight loss or prolonged tetraplegia for more than 24 hours without signs of recovery (humane endpoint).

**Figure 2.**
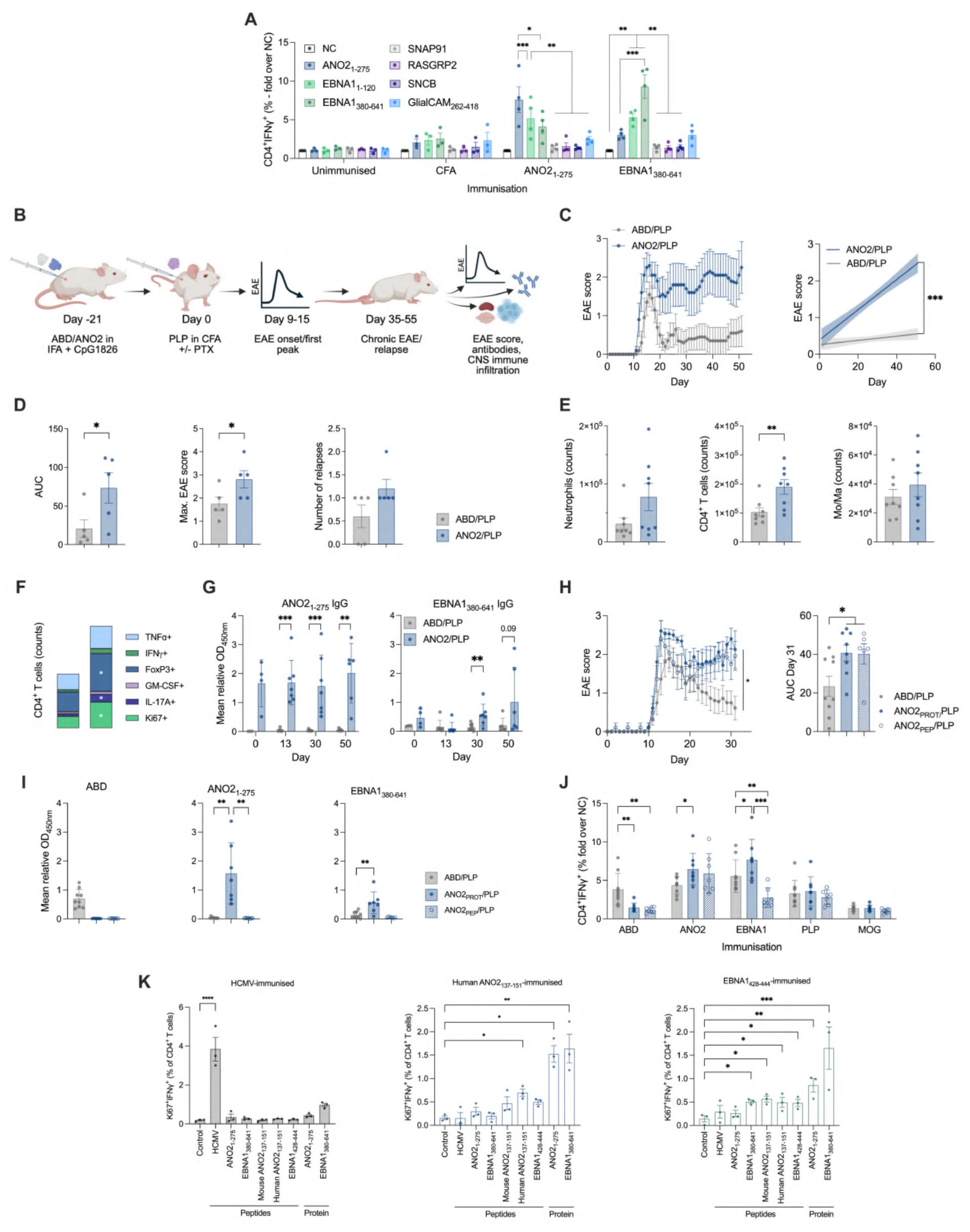
ANO2_1-275_ pre-immunisation leads to more severe EAE and primes T cell and antibody responses to EBNA1_380-641_. **A.** Draining inguinal lymph node lymphocytes from young SJL/J mice with no immunisation (n=3), complete Freund’s adjuvant (CFA) immunisation (n=3), ANO2_1-275_ immunisation (n=4) or EBNA1_380-641_ immunisation (n=4) were harvested at day 11 and rechallenged *in vitro* with antigen-coupled beads and processed by intracellular cytokine staining and flow cytometry. Data presented as the fold change of CD3^+^CD4^+^IFNγ^+^ T cells over negative control (NC) beads. Data analysed using a two-way ANOVA mixed effects model with Dunnett’s test for multiple comparisons. **B.** Schematic representation of EAE experimental procedure. Briefly, SJL/J mice were pre-immunised with ABD or ANO2_1-275_ protein with incomplete Freund’s adjuvant (IFA) + CpG1826. After 21 days, mice were immunised with PLP + CFA with or without pertussis toxin (PTX) and EAE onset was monitored through its peak at day 9-15 and up to 55 days post-immunisation. Tissue from animals at peak disease and the chronic phase were taken for analysis. **C.** (Left panel) Clinical course of EAE represented by EAE score of ABD/PLP (n=5) and ANO2_1-275_/PLP-immunised (n=5) mice up to day 55. (Right panel) Linear regression curve of EAE scores of ABD/PLP (n=5) and ANO2_1-275_/PLP-immunised (n=5) mice. Data from sick animals only, shaded area represents the 95% confidence interval. **D.** Clinical phenotype of mice sacrificed at day 51 post-immunisation represented by EAE score area under curve (AUC, left), maximum EAE score (centre), and number of relapses (right) in ABD/PLP (n=5) and ANO2_1-275_/PLP-immunised mice (n=5). Data was non-normally distributed and was analysed using the Mann-Whitney test. **E.** Frequency of neutrophils (left), CD4^+^ T cells (centre) in the brains, and monocytes and macrophages (Mo/Ma, right) in spinal cords of ABD/PLP (n=8) and ANO2_1-275_/PLP immunised mice (n=8) sacrificed at day 51. Mann-Whitney test. **F.** Cytokine production of spinal cord-infiltrating CD4^+^ T cells of ABD/PLP (n=7, pooled) and ANO2_1-275_/PLP-immunised (n=7, pooled) mice sacrificed at day 30 was analysed by ICS and flow cytometry (total cell number for each condition: 80,694). Mann-Whitney test. **G.** Sera from ABD/PLP and ANO2_1-275_/PLP-immunised mice were tested for ANO2_1-275_ IgG reactivity by ELISA at priming (ABD n=3, ANO2 n=4) and at days 13 (ABD n=7, ANO2 n=7), 30 (ABD n=9, ANO2 n=7) and 50 (ABD n=4, ANO2 n=6) post-immunisation. Data presented as absorbance at 450nm (OD_450nm_), statistical test is the multiple Mann-Whitney test. **H.** Similar to EAE experimental procedure described in Figure 2B, SJL/J mice were pre-immunised with ABD, ANO2_1-275_ protein or an ANO2_1-275_ peptide pool (15mers overlapping by 11aa) with incomplete Freund’s adjuvant (IFA) + CpG1826. After 21 days, mice were immunised with PLP + CFA with or without pertussis toxin (PTX) and EAE onset was monitored through its peak at day 9-15 and up to 31 days post-immunisation. Tissue from animals at peak disease and the chronic phase were taken for analysis. EAE score for pre-immunised animals with ABD (n=9), ANO2_1-275_ protein (n=8) or ANO2_1-275_ peptides (n=6) during the 31-day follow-up period. Area under curve (AUC) for EAE scores at day 31 following induction of EAE between groups (right panel). Unpaired T-test. **I.** Antibody reactivity to ABD (left, dilution 1:300,000), ANO2_1-275_ (middle, dilution 1:24,000) and EBNA1_380-641_ (right, 1:6000) was analysed in sera harvested from animals at day 31 post-EAE induction by in-house ELISA. Unpaired T-test. **J.** Splenocytes from SJL/J mice pre-immunised with ABD (n=9), ANO2_1-275_ protein (n=8) or ANO2_1-275_ peptides (n=7) were rechallenged with ABD beads, ANO2_1-275_ beads, EBNA1_380-641_ beads, PLP peptides or MOG peptides at day 30 and analysed for T cell reactivity by flow cytometry and ICS. Data is presented as the fold change (FC) %CD4^+^IFNγ^+^ T cells over the negative stimulation control (NC). 2-way ANOVA with Dunnet’s multiple comparisons test. **K.** Splenocytes from SJL/J mice which were immunised with HCMV IE1_162-173_ peptide (n=3), human ANO2_137-151_ peptide (n=3) and EBNA1_428-444_ peptide (n=3) were rechallenged *in vitro* with each of the antigens used for original immunisations to test for antigen-specific priming of T cell responses. Extended data in Figure S12D. Responses were determined by ICS and flow cytometry and indicated by Ki67 and IFNγ expression. One way ANOVA with Dunnet’s multiple comparisons test. Albumin binding domain (ABD), anoctamin-2 (ANO2), proteolipid protein (PLP), myelin oligodendrocyte glycoprotein (MOG), negative control (NC), complete Freund’s adjuvant (CFA), Pertussis toxin (PTX), incomplete Freund’s adjuvant (IFA), 91 kDa synaptosomal-associated protein (SNAP91), RAS guanyl-releasing protein 2 (RASGRP2), synuclein-β (SNCB), glial cell adhesion molecule (GlialCAM), optical density (OD). \**P* < 0.05; \*\**P* < 0.01; \*\*\**P* < 0.001. Created with https://BioRender.com.

**Figure 3.**
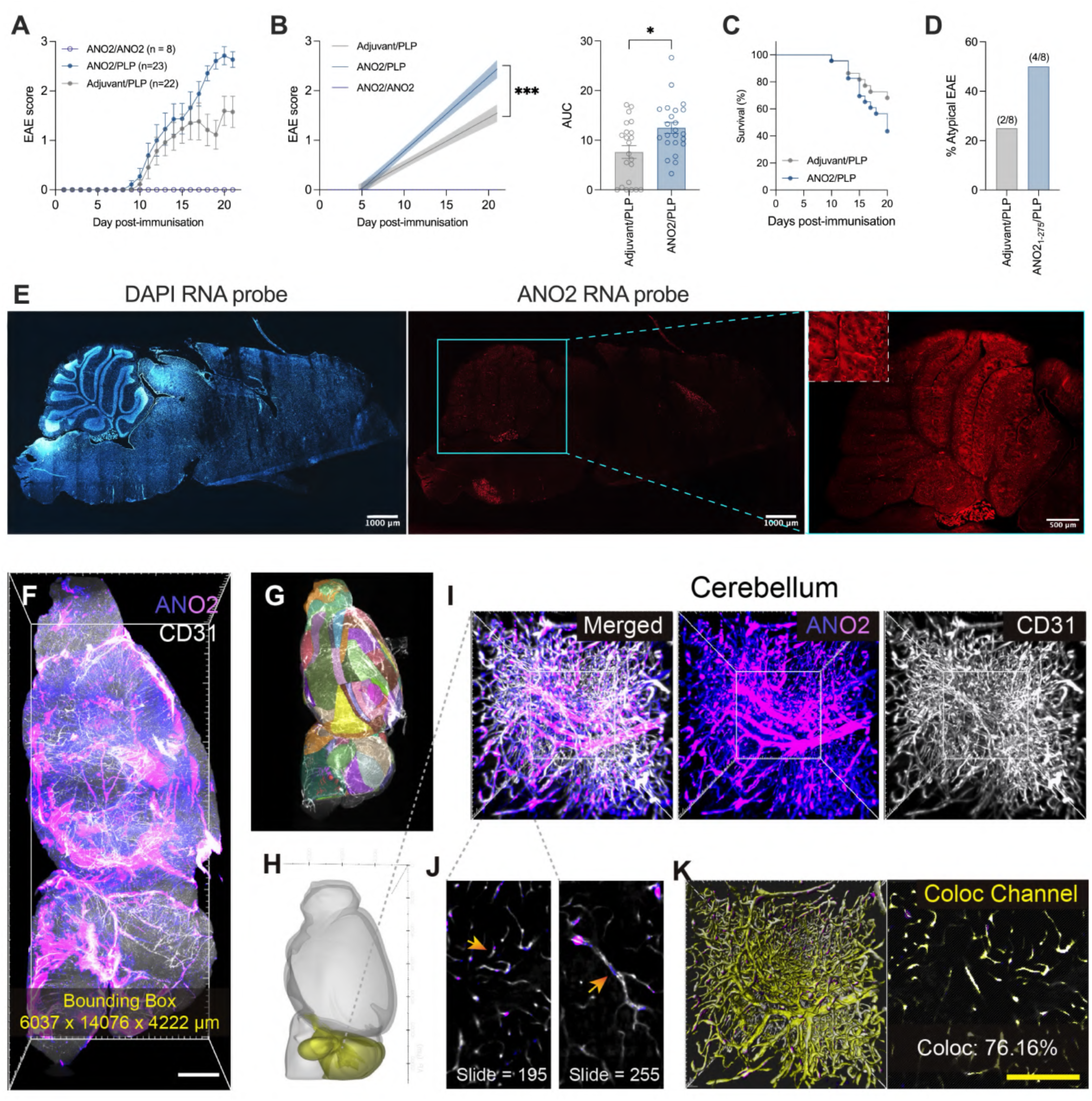
ANO2_1-275_ pre-immunisation of adult mice induces an atypical EAE phenotype and suggests targeting of brain areas with ANO2 expression. **A.** Six-month-old mice were immunised with adjuvant only (n=22) or ANO2_1-275_ (n=31) protein in adjuvant, and three weeks later EAE was induced by immunisation with PLP in CFA or ANO2_1-275_ in CFA (as schematic in Figure 2B). EAE score of mice immunised with adjuvant/PLP (n=22), ANO2_1-275_/PLP (n=23) or ANO2_1-275_/ANO2_1-275_ (n=8) including unaffected animals and deceased animals as score 5. Pooled data of 3 independent experiments. **B.** (Left panel) Linear regression curve of EAE scores for mice immunised with Adjuvant/PLP (n=22), ANO2_1-275_/PLP (n=23) or ANO2_1-275_/ANO2_1-275_ (n=8). Global F-test to test whether regression lines differ across groups. (Right panel) Area under the curve (AUC) of EAE scores for mice immunised with Adjuvant/PLP (n=22) or ANO2_1-275_/PLP (n=23). Mann-Whitney test. **C.** Survival curve of animals immunised with Adjuvant/PLP (n=22), ANO2_1-275_/PLP (n=23). **D.** Proportion of animals which presented with an atypical EAE phenotype; adjuvant/PLP (n=8), ANO2_1-275_/PLP (n=8). One representative experiment. **E.** RNAscope to determine expression of *Ano2* transcripts in mouse brains. *Ano2* indicated by white staining. **F.** Whole-mount visualisation and regional co-expression analysis of ANO2 and CD31. Volumetric rendering of the representative brain hemisphere of a SJL/J mouse (9 imaged overall). ANO2 signal is displayed using an intensity-based gradient from violet to pink (ULTRA colormap), and the classical endothelial marker CD31 is shown in white. Bounding Box: 6037 x 14076 x 4222 μm. Scale bar = 1000 μm, white. **G.** The whole-brain dataset registered to the Allen Mouse Brain Atlas at 25μm resolution. **H.** Detailed analysis of the cerebellum based on ANO2 and CD31 co-expression patterns (highlighted in yellow). **I-K.** Representative images and co-localisation analysis of ANO2 and CD31 expression in the cerebellum. Scale bar = 250 μm, yellow. \**P* < 0.05; \*\**P* < 0.01; \*\*\**P* < 0.001.

At different time points after EAE induction, animals were sedated with isoflurane, perfused transcardially with 50ml of PBS, and the brain, spinal cord, and spleens were dissected for further experiments. Splenocytes were mechanically dissociated, erythrocytes lysed with ACK and cells spun and resuspended in RPMI 1640 complemented with 5% foetal bovine serum, 1% L-glutamine, 1% penicillin-streptomycin, 1% pyruvic acid, 1% MEM non-essential amino acids, 10mM HEPES and 50mM 2-mercaptoethanol. Cells were plated at 0,2×10^6^ cells per well in U-bottom plates and restimulated with the immunisation antigen and a panel of control as well as putative cross-reactive antigens, both bead-conjugated as well as native purified proteins and peptides thereof (see figure legends). After 96 hours, cells were replated into V-bottom plates and restimulated for 5 hours with PMA (50ng/ml), ionomycin (1μg/ml), and GolgiPlug (1μl/ml). Samples were subsequently stained with antibodies according to Table S3. LIVE/DEAD Fixable Near-IR or Yellow dead cell exclusion dyes (L34976 or L34959, Invitrogen) were used to exclude dead cells. Intracellular/intranuclear staining and flow cytometry acquisitions were performed as above (Table S3B). Gating strategy in Figure S10B and S10C.

Brains and spinal cords were mechanically dissociated and resuspended in 3ml of an enzyme mix containing 0.2mg/ml DNAseI (Roche) and 1mg/ml Collagenase-D (Roche) in HBSS. Tissues were incubated twice for 30 minutes at 37°C, followed by further dissociation through repeated pipetting. After a total of 60 minutes of incubation the homogenate was filtered through a 40μm cell strainer, spun down and resuspended in 10ml of an isotonic 38% Percoll (Sigma-Aldrich) solution in HBSS and centrifuged at 800g (no break) for 10 minutes at 4°C. The pellet was carefully collected with a 1ml pipette and washed in 15ml HBSS by centrifugation at 400g for 10 minutes at 4°C. For flow cytometry analysis, cells were resuspended in ice cold PBS, transferred to a 96-well V-bottom plate and stainings were performed as described above. For antibodies see Table S3C. For analysis, samples were only excluded if presented with obvious signs of technical problems in perfusion, myelin clearance or flow cytometry acquisition-related problems.

### Adoptive transfer EAE

For the induction and clinical evaluation of EAE induced exclusively by T cells, donor SJL/J animals were immunised with either PLP_139-151_, ANO2_1-275_ peptide pool of 15-mers overlapping by 11aa or the HCMV IE1_162-173_ peptide as a control, all in CFA as described above. Spleen and lymph nodes were harvested 9-10 days post-immunisation, single-cell suspensions were generated and cells were restimulated *in vitro* in RPMI-1640 complemented with 10% foetal bovine serum, 1% l-glutamine, 1% penicillin-streptomycin, 1% pyruvic acid, 1% MEM non-essential amino acids, 10mM HEPES and 50mM 2-mercaptoethanol at a density of 3×10^6^ cells/ml in medium containing 10μg/ml of respective peptides and 20ng/ml of recombinant mouse IL-23 (R&D). After 3 days, cells were collected, counted, resuspended in 200μl PBS at a density of 25-50×10^6^ cells/ml and were transferred intraperitoneally into recipient mice. EAE development and scoring was performed as described above.

For T cell tracking experiments we made use of C57BL/6 animals that express GFP from the human 10ubiquitin C promoter (JAX, Strain 004353) crossed with SJL/J mice. The F1 of these strains readily develops EAE in response to both MOG_35-55_ and PLP_139-151_ -primed adoptively transferred T cells. Here we opted for MOG_35-55_ transfer since MOG-specific T cells primarily target the white matter regions of the CNS, including the spinal cord and optic nerves. SJL/J x C57BL/6-GFP animals were immunised with either MOG_35-55_ or ANO2_1-275_ peptide pool and cells were harvested, restimulated *in vitro* and finally adoptively transferred into non-GFP SJL/J x C57BL/6 F1 congenic animals. EAE development and scoring was performed as described above.

### ELISAs to detect recombinant antigen IgG responses in mice

ELISAs were optimised for detection of recombinant antigen IgG responses in immunised animals. Sera was collected from sacrificed animals by cardiac puncture. MaxiSorp 96-well plates were coated with respective antigens (5µg/ml, overnight incubation at 4°C). Each plate included wells for blank measurements and wells coated with BSA (5µg/ml) were used as controls. The plates were washed five times with wash buffer (PBS with 0.05% Tween-20) and subsequently blocked with blocking buffer (PBS with 3% BSA and 0.05% Tween-20) for 1 hour at room temperature (RT). After washing, sera was diluted (ABD 1:300,000, ANO2_1-275_ 1:24,000, EBNA1_380-641_ 1:6,000) in ELISA buffer (PBS with 0.2% BSA and 0.05% Tween-20) and incubated for 2 hours at RT. The plates were washed and secondary antibody (HRP-conjugated goat anti-mouse IgG, Jackson Laboratories, 1:2000) was added before incubation for 1 hour at RT in the dark. After washing, plates were developed with TMB-substrate (Sigma) for 10 minutes and the reaction was stopped with 0.5M H_2_SO_4_. Optical density (OD) values were read at 450nm using SpectraMax plus 384 ELISA reader (Molecular devices). For titration of antibody dilutions see Figure S11A and S11B.

### Plasma labeling and injection

Plasma proteins were labeled with NHS-ester fluorescent dye as previously described^43^. Briefly, 2mL EDTA-isolated plasma was reacted with 1mg NHS-ATTO647N (ATTO-TEC) at RT for 1.5h. Unreacted dye was removed by overnight dialysis in PBS at 4°C using a 10kDa MWCO membrane, followed by quenching with 50mM Tris (pH 8.0). Labeled plasma was washed once with Amicon Ultra Centrifugal filter unit (10kDa MWCO) followed by four washes on Zeba spin desalting columns (7kDa MWCO). Free dye removal (∼98%) was confirmed by processing plasma incubated with non-reactive carboxy-ATTO647N under the same conditions. Protein concentration was measured by nanodrop and plasma was kept refrigerated until intravenous injection. For injections, EAE mice were injected in the tail vein at 5.5µl plasma per gram of body weight and incubated two hours before the sacrifice.

### Tissue processing and sectioning

In all the experiments, anesthetized animals were transcardially perfused with PBS followed by either 4% PFA or 4% formaldehyde. The spinal cords and brains were post-fixed for 12h or overnight in 4% PFA or 4% formaldehyde, and subsequently kept in 30% sucrose overnight for cryoprotection. Tissues were embedded in Tissue-Tek O.C.T Compound (Sakura), frozen and sectioned at 20-25 µm using a Leica CM1850 cryostat before being mounted on Adhesion Microscope Slides (Epredia). Sections were subsequently stored at -20°C until further processing.

### Immunohistochemistry

Slides retrieved from -20°C storage were left at RT to air dry before washing with PBS. For sections stained with ANO2, antigen retrieval was performed by incubating slides in immunohistochemistry (IHC) antigen retrieval solution (Invitrogen, 00-4955-58) at 90°C for 10 minutes. Unbroken circles were then drawn around the samples with a PAP pen to create a hydrophobic barrier. Sections were subsequently blocked for 1h at RT with blocking solution (5% normal donkey serum with 0.3% Triton-X in PBS), followed by overnight incubation at 4°C in a humidified chamber with the following antibodies according to the experiment: donkey anti-goat CD31 antibody (1:250, R&D systems, AF3628), goat polyclonal Iba1 antibody (1:300, Abcam, ab5076), rabbit polyclonal anti-MBP antibody (1:200, ProteinTech, 10458-1-AP), rabbit anti-ANO2 (1:200, Alomone Labs, LTD #ACL-012), rabbit anti-CD4 (1:200, Abcam, #ab18365), goat anti-Podocalyxin (1:300, R&D Systems, AF1556). After rinsing with PBS, the sections were incubated for 1h at RT with the following secondary antibodies: donkey anti-goat Cy3 (1:500, Jackson ImmunoResearch, 705-165-003), donkey anti-goat IgG (H+L) Alexa Fluor Plus 647 (1:1000, Invitrogen, A32849), donkey anti-rabbit IgG (H+L) Alexa Fluor Plus 488 (1:1000, Invitrogen, A32790). All sections were stained with Hoechst (ThermoFisher Scientific, 62249) before mounting with antifade mounting medium (Invitrogen, P36984). Full brains and spinal cords, along with specific anatomical regions including the cortex, hippocampus, corpus callosum, brain stem, cerebellum, and the ventral and dorsal column of spinal cords were captured with a Zeiss LSM800 or Zeiss LSM 880 confocal microscope using the ZEN software.

### Image analysis

All images were analyzed using ImageJ (Fiji) software (version 1.54j). Plasma leakage quantification was obtained by calculating the percentage of Atto647-positive area relative to the total tissue area. A uniform threshold (Triangle) was applied to isolate the specific signal, using identical parameters across all samples to ensure consistency. The Iba1- and MBP-positive areas were quantified using the same thresholding conditions, focusing on defined regions of interest (ROIs). The Iba1-positive cell numbers were determined by manual counting of ROIs and expressed as cells per mm². In total, 4–5 brain sections and 5–8 spinal cord sections were analyzed per mouse.

### RNAscope

Sagittal brain sections from SJL/J x C57BL/6 F1 congenic animals at peak of disease were used for assessing expression of Ano2 using the Ano2-C3 probe conjugated to TSA Vivid Fluorophore 570 (Cy3). Brain sections were first treated with 1x target retrieval buffer supplied by the HiPlex v2 Assay (ACDbio) for 5min at 98°C, and subsequently processed with probe hybridization and fluorescence conjugation (RNAscope™ Multiplex Fluorescent Reagent Kit v2) according to the manufacturer’s instructions. Sections were subsequently counterstained with DAPI. Images were acquired using confocal microscope LSM 980 Airyscan (Zeiss).

### Sample collection, preparation and whole-mount immunostaining using iDISCO+

Nine mouse brains were harvested and fixed overnight at 4 °C in 4% paraformaldehyde (PFA). After fixation, a graded methanol dehydration series was performed at RT (60%, 80%, 100%, 100%; Sigma-Aldrich, Germany), and the samples were stored at −20 °C. For downstream process and analysis, the right hemispheres were carefully dissected using a sterile scalpel.

The samples were equilibrated at 4 °C and incubated overnight in 66% dichloromethane (DCM) in methanol (Sigma-Aldrich, Germany). This was followed by two 30-minute washes in 100% methanol and overnight bleaching at 4 °C in 5% hydrogen peroxide in methanol. Rehydration was performed through sequential 1-hour incubations in 80%, 60%, 40%, and 20% methanol, followed by 1 hour in PBS at RT. The samples were then washed for 1 hour in PBS containing 0.2% Triton X-100 (PTx.2; Sigma-Aldrich, Germany) and permeabilized by incubating for 2 days at 37 °C in a permeabilization solution composed of 20% DMSO and 2.3% glycine in PTx.2. To block nonspecific binding, samples were incubated at 37 °C for 2 days in a blocking buffer containing 6% normal donkey serum (Jackson ImmunoResearch) and 10% DMSO in PTx.2. Primary antibody labelling was carried out in PTwH buffer (PBS supplemented with 0.2% Tween-20 and 0.1% of a 10 mg/ml heparin stock solution, using Anti-ANO2 (1:100; Proteintech, 67638-1-Ig) and Anti-CD31 (1:100; R&D Systems, Cat#AF3628). Samples were incubated at 37 °C for 9 days, followed by 4–5 washes in PTwH over the course of the next day. Secondary antibody staining was performed with donkey anti-mouse IgG conjugated to Alexa Fluor™ 647 (Jackson ImmunoResearch, 715-606-150) and donkey anti-goat IgG conjugated to Alexa Fluor™ 488 (Jackson ImmunoResearch, 109-605-006), also for 9 days. Samples were rinsed in PTwH buffer 4–5 times over the course of a day. This was followed by stepwise dehydration through a graded methanol series (60%, 80%, 100%, 100%), with each step lasting 1 hour at RT. The dehydrated tissues were then incubated in 66% dichloromethane (DCM) in methanol for 3 hours. To remove residual methanol, samples were washed twice for 15 minutes in 100% DCM. Final tissue clearing was performed using dibenzyl ether (DBE; Sigma-Aldrich, Germany).

Samples were mounted in 100% dibenzyl ether for image acquisition. Data were acquired using an UltraMicroscope Blaze system (LaVision Biotec) equipped with a 4× immersion objective. Optical sectioning was performed with a z-step interval of 2 µm, generating a voxel resolution of 1.62 × 1.62 × 2 µm. Fluorophores corresponding to CD31 and ANO2 were excited using 488 nm and 647 nm laser lines, respectively. All specimens were imaged under identical acquisition settings to ensure consistency. Raw image stacks were subsequently stitched and converted into Imaris file format for downstream analysis.

For image registration and analysis^45^, half-brain registration was performed using the Brainreg module from the BrainGlobe suite, referencing the Allen Mouse Brain Atlas at 25 µm resolution. ROIs were localised using Brainrender. Whole-brain image stacks were imported into AmiraTM 3D (ThermoFisher, Version 2024.1) via a custom-developed Python script. After down-sampling to 8 x 8 x 8 µm, volumetric segmentation was performed on the entire dataset. To suppress nonspecific high-intensity signals along tissue boundaries, a 4-voxel erosion filter was applied. Segmentation results were subsequently resampled and applied across all fluorescence channels. The processed datasets were converted using ImarisFileConverter (Oxford Instruments, Version 10.2.0) and imported into Imaris (Oxford Instruments, Version 10.2.0) for volume rendering, region-based segmentation, image export, and animation. Colocalisation analysis between CD31 and ANO2 was performed using intensity threshold-based detection in Imaris, generating dedicated normalised ion channels for quantitative assessment.

### Coculture experiments

#### Primary oligodendrocyte precursor cell (OPC)/mature oligodendrocyte (MOL) cell culture

Mouse brains from P4-7 pups were removed and dissociated into single-cell suspensions using the Neural Tissue Dissociation Kit P (Miltenyi) and OPCs were obtained through MACS sorting with CD140a microbeads (Miltenyi) following the manufacturer’s protocols. Cells were seeded in poly-L-lysine (Sigma) plus fibronectin (Sigma) coated flat bottom 96-well plates and grown on proliferation media comprising DMEM F12 Glutamax, N2 media, Pen/Strep (all from ThermoFisher Scientific), NeuroBrew (Miltenyi), bFGF at 20ng/ml and PDGF-AA at 10ng/ml (both from Peprotech). For OPC differentiation, cells were left for 2 days in medium without bFGF and PDGF-AA. For differentiation into MOLs, cells were kept in the same medium with the exception of bFGF and PDGF-AA and the addition of T3 at 40ng/ml for 48h. For the induction of disease-associated OPC and MOL phenotype, cells were treated with 50ng/mL IFNγ (R&D) for 24 hours.

#### Primary microglia and astrocyte cell culture

PBS-perfused mouse brains from adult mice (2-3 months old) were removed, placed into Leibovitz L15 medium (ThermoFisher) containing 0,1mg/ml papain (Worthington) and dissociated into single-cell suspensions using 3 rounds of 5 min incubation at 37°C intercalated by pipetting. DnaseI (Roche) at 0.2mg/ml was added during the last step. The enzymatic digestion was stopped by adding HBSS containing 10% FBS and centrifuged at 200g for 7 min at RT. The pellet was resuspended in 20% isotonic Percoll (Sigma) in HBSS and once again centrifuged at 200g for 7 min RT with low acceleration and no brake, then washed and resuspended in DMEM/F12 (Gibco) with 10% FBS, Pen/Strep, Glutamine and 20ng/ml M-CSF (R&D). After 10 days, cells were trypsinized and cell suspension was MACS sorted using CD11b beads (Miltenyi) into a negative astrocyte fraction and a positive microglia fraction. Cells were counted and plated at 100 000 cells/well in complete DMEM/F12 medium in flat bottom 96-well plates for 24h.

#### Primary neuron cell culture

Neuronal cells for co-cultures were established from cells isolated from P5 mouse CNS dissociated using the Postnatal Neural Tissue Dissociation Kit (Miltenyi) according to the manufacturer’s instructions. Neuronal cells were isolated using the Neuron Isolation Kit (Miltenyi) and cultured in MACS Neuro Medium containing MACS NeuroBrew-21 (both from Miltenyi), 1% Pen/Strep and 0.5 mM L-glutamine on poly-L-lysine coated flat bottom 96 well plates for 10 days.

#### Primary bone marrow derived dendritic cell (BMDC) culture

Femurs from 2-4 months old mice were collected, briefly dipped in 70% ethanol (EtOH) and washed in PBS. The bone epiphysis was cut on both sides and the bone marrow was flushed out with 5ml PBS per bone into a petri dish using a 10ml syringe and a 23-gauge needle. The bone marrow of 2 bones was collected in a 15ml Falcon tube and dissociated by pipetting. Cells were centrifuged and erythrocytes lysed in ACK lysis buffer (Sigma). Cells obtained from 1-2 femurs were resuspended in 30ml DMEM (Sigma) including glutamine, 20% FBS, 1% Pen/Strep and 20ng/ml rmGM-CSF (R&D) for BMDCs in T75 culture flasks with low adherence growth surface. Medium was changed twice, returning non-adherent cells back to the flask. After 10 days, cells were stimulated with 500 ng/ml LPS O111:B4 (Invivogen). After 24h, activated DCs were thoroughly washed, detached with 0.05% Trypsin-EDTA and counted.

#### RNA extraction

RNA was isolated using RLT Lysis Buffer (#79216, Qiagen, Hilden, Germany) and purified by Rneasy Mini kit (74104, Qiagen), including Dnase digestion. The concentration was measured with Multiskan SkyHigh Spectrophotometer (Thermo Scientific, Waltham, MA).

### Quantitative PCR (qPCR)

For qPCR, total RNA was reverse transcribed using an iScript cDNA Synthesis Kit (#1708891, Bio-Rad). PCR amplification was performed in 384-well plates in a CFX Opus 384 Real-Time PCR System (Bio-Rad), using TaqMan® Fast Advanced Master Mix (#4444556, Fisher Scientific) and TaqMan® Gene Expression Assays (H2-Ab1 *Mm00439216_m1* and ANO2 *Mm00463894_m1*, Applied Biosystems). All samples were measured in duplicate. Results were normalised to the equal mass of total RNA as well as the Ct values of RPLP0 (Mm00725448_s1) housekeeping control. The relative amount of target gene mRNA was calculated either by the 2^-ΔCt method and presented as relative expression to the housekeeping gene or by the 2^-ΔΔCt^ method and presented as a fold change compared to the baseline expression in negative controls.

#### Incucyte time-lapse imaging

For coculture experiments, target cells (microglia, astrocytes, neurons, OPCs, MOLs) described above were obtained from SJL/J x C57BL/6-GFP F1 animals, while T cells and dendritic cells were obtained from SJL/J animals, to allow for discrimination between GFP-positive targets and GFP-negative effector cells. ANO2-specific effector T cells were generated as described for adoptive transfer experiments after cells were harvested from immunised mice, restimulated in vitro for 3 days with peptide pools and rested for 7 days in 20ng/ml rmIL-2 (R&D). While numbers of target cells could not be kept constant across different cell types due to plating/replating constrains and proliferation rates, they were constant within one single experiment and cell type. On the day of the assay, target cells were incubated with 50 000 DCs and 100 000 resting ANO2-specific T cells in RPMI-1640 complemented with 10% FBS, 1% l-glutamine, 1% Pen/Strep, 1% pyruvic acid, 1% MEM non-essential amino acids, 10mM HEPES and 50mM 2-mercaptoethanol containing Incucyte® Caspase-3/7 Dye for Apoptosis assessment (Sartorius). Combinations of targets cells with either DCs or T cells or both were included. Conditions including the addition of ANO2 peptide pools were included as positive controls for T cell activation. Plates were immediately transferred to an incubator connected to an Incucyte ZOOM instrument in which wells were analysed using 20x ocular magnification every 4-12 hours for up to 50 hours. Target cell numbers as well as the number of Caspase-3/7-positive cells was recorded and analysed using the Incucyte 2024A (Sartorius) software.

### Bioinformatic analysis of T cell receptor sequences

#### QC and demultiplexing of cells

Unfiltered-count matrices from CellRanger were loaded into R(4.2.3) for preliminary QC analysis. DropletUtils was used to remove empty droplets from the data with default parameters. Low-quality cells were excluded from the dataset on a per-sample basis by excluding cells which exhibited nUMIs or nGenes outside of 3 median absolute deviations from the sample. Cells that exhibited a fraction of mitochondrial reads greater than 15%, presumably due to damage and/or death, were also removed from the dataset. The HTODemux function in Seurat was called to classify cells by hashing-antibodies. Cells without sufficient hashtag or classified as multiplets were removed from downstream analysis. Filtered and de-hashed count matrices from each sample were aggregated and exported to an .h5 file with AnndataR.

#### QC of VDJ data

TCR matrices for each sample were loaded into python (3.10) with Scirpy and aggregated into a single anndata object. A multi-modal dataset containing TCR and transcriptome data was created using the muon-package. Only cells that contained both transcriptional data and TCR data were kept for downstream analysis. Selection of primary/secondary chains, and annotation of TCRs was performed with scirpy.pp.index_chains. TCRs containing an incomplete TCR pair (orphan VJ/VDJ), or that contain multiple VJ/VDJ (> 2. multichain) were removed from the dataset. Number of TCR clonotypes following data processing and QC in Table S5.

#### Clustering of clonotypes and identification of cross-reactive clonotypes

We assigned unique TCR groupings based on a strict-clustering of identical amino acid sequence of VJ/VDJ CDR3 regions and enforcement of identical V-gene using the Scirpy package scirpy.tl.clonotype_clusters. Briefly, a distance matrix is computed for each of the CDR3 regions, where 1 corresponds to an identical amino acid sequence and 0 between sequences with a mismatch (parameter: sequence = identity). All V-genes from both VJ and VDJ arms are enforced to match, as specified by the function parameters receptor arms = all, and v_gene = same. Clonotype clusters were enforced to be distinct for each donor separately. Clonotypes were considered expanded if >4 cells were found in a given condition (anti-CD3, ABD, ANO2, EBNA1 or CRYAB) or if >10% of cells in a given clone were derived from a specific stimulation. For each clonotypes, the larger threshold of the two were chosen. Cells that met classification criteria in multiple conditions were considered cross-reactive given expansion in multiple conditions. The cut-off of 4 was chosen considering a hypothetical scenario where 1 cell with a given TCR was initially seeded in the culture would undergo two cell divisions, and a cutoff of 10% of the composition of a clone was chosen to account for potential differences in initial seeding. Similarity of TCRs was assessed by clustering based on amino acid similarity with scirpy.tl.clonotype_clusters. A distance-matrix is calculated between the CDR3 regions using the BLOSUM62 matrix (parameter: sequence = alignment). V-genes from both VJ & VDJ were enforced to be identical, and similarity wascalculated within each donor group.

#### Clonal entropy and diversity

Shannon’s entropy and clonal diversity were calculated for each cell using functions implemented in the Scirpy package. Mean-diversity/entropy was calculated for each group for visualisation.

#### Jaccard similarity scoring

A presence/absence matrix was generated, where a 1 denoted sampling of a given TCR in a donor/condition combination. The vegan package was used to calculate jaccard dissimiliarty between conditions in each individual donor. Similarity scores were reported as 1 – jaccard dissimilarity for each comparison.

#### V(D)J gene-usage

A binomial model was fitted to each gene using the gamlss package. Trials was defined as the number of unique TCRs for each donor:classification, and successes were defined as the number of unique TCRs for each donor:classification that contained a given V(D)J gene. A fixed-effect for donor was added to the model to account for the paired-nature of the samples. Due to constraints on the number of donors, as well as the number of TCRs obtained for certain classifications, only v-genes which could be identified in >2 classifications in >2 donors were included.

#### Comparison with public datasets

TCR data from Gottlieb *et al*^42^ were retrieved from GEO [GSE250037], and samples were aggregated into one dataset. The dataset was reduced to unique CDR3βs. TCR data from Ballerini was accessed with the DOI included in the original publication. Similarly, .csv files containing CDR3 β regions from each sample were aggregated into a single dataset and reduced to unique CDR3β sequences. The number of identical CDR3bs overlapping between the three datasets was reported.

#### Transcriptome analysis

Cells that passed quality-control for VDJ sequences were further processed for transcriptome analysis. Genes with low-expression, detected in <3 cells, were removed from the count matrix. Ribosomal (RSP|RPL), mitochondrial (MT-) and MALAT1 were removed from the count matrix. Library normalization (scanpy.pp.normalise total: target_sum = 1e4) and log-transformation (scanpy.pp.log1p) were performed. Highly-variable features were calculated to use for dimensional reduction (sc.pp.highly_variable_genes: Flavor = Seurat_v3, n_top_genes = 2000), and data was scaled clipping values at 10. Principal Component Analysis (PCA) on the top 2000 features was performed. The python implementation of Harmony^46^, harmonypy was used to resolve differences in clustering due to donor origin. Harmonized principal components were used to construct a nearest-neighborhood graph (n_pcs = 15). The number of PCs was arbitrarily chosen, as clustering was robust across a wide-range of parameters. UMAP dimensional reduction and leiden clustering were computed using the constructed neighborhood-graph. Initial analysis resulted in nine clusters, however clusters 8 & 9 had low numbers of cells and expressed genes suggesting contamination (e.g. CD14) and were therefore removed from downstream analysis.

#### Cell-cycle scoring

Cell-cycle gene list was obtained from regev_lab^47^, which contains annotated S-genes and G2M genes. Using scanpy’s score_cell_cycle function, a phase score was assigned to each of the cells.

#### Differential Expression Analysis

Sum-aggregated pseudobulk profiles were generated from cells from each donor:cluster. To identify cluster-specific markers, each cluster was contrasted against the aggregate profile of the remaining cells. Differential expression was performed with the python implementation of DESeq2, PyDESeq2. A negative-binomial model was constructed for each gene, with donor as a fixed-effect to account for pairedness of our data with default parameters.

#### Geneset Enrichment Analysis

Following differential gene analysis, for each cluster GSEA was performed. Log2FoldChanges for each comparison were used to construct a gene-ranking using the fgsea package, and the hallmark-pathways obtained from the msigdbr package were used to calculate pathway enrichment.

#### Python Packages

Scirpy 0.19.0^45^

Scanpy1.10.3^46^

Muon0.1.7^47^

AnnData^48^

Harmonypy (https://github.com/slowkow/harmonypy)

Leiden^48^ (https://github.com/vtraag/leidenalg)

PyDESeq2^49^

Decoupler^50^

#### R Packages

DropletUtils1.18.1^43^

Seurat4.3.0^44^

AnndataR 0.99.0 (https://github.com/scverse/anndataR)

ggforce_0.4.1 (https://ggforce.data-imaginist.com)

VennDiagram_1.7.3^51^

ggalluvial_0.12.5^52^

vegan_2.6-4 (https://github.com/vegandevs/vegan)

ComplexUpset_1.3.3 (https://github.com/krassowski/complex-upset)

dplyr_1.1.1 (https://dplyr.tidyverse.org)

ggplot2_3.4.2

Cowplot

GGally (https://ggobi.github.io/ggally/)

Fgsea (http://biorxiv.org/content/early/2016/06/20/060012)

Msigdbr7.5.1 ( https://igordot.github.io/msigdbr/authors.html#citation)

#### Code availability

https://gitlab.com/jagodiclab and full code will be supplied upon request.

### Statistical analysis

Non-parametric statistical tests were used for all analyses. The Wilcoxon signed-rank test with Holm-Sidak correction for multiple comparisons were used to test comparison to baseline measurements in paired samples. The two-tailed non-parametric Kruskall-Wallis test with Dunn’s multiple comparisons test was used for analysis of multiple groups. Correlations were measured with a two-tailed non-parametric Spearman correlation (r) with Holm-Sidak correction. Two- and one-way analyses of variance (ANOVA) were used with Tukey’s multiple comparisons test to analyse T cell responses in EAE antigen recall experiments. Statistical analyses were performed in GraphPad Prism v10.

## Results

### T cell responses targeting Anoctamin-2 are elevated in persons with multiple sclerosis

We first investigated whether ANO2 is also the target of T cell responses in pwMS using stimulation with antigen-coupled beads followed by IFNγ, IL-17A and IL-22 FluoroSpot^40,42^ (Figure 1A). Recombinant proteins covering Cytomegalovirus (CMV) pp65, albumin binding domain (ABD, control protein) and ANO2_79-168_ (with homology to EBNA1) were coupled to paramagnetic beads as previously described^40^ (Figure S1A-B). For the analysis, PBMC were collected in two separate cohorts: cohort 1 comprised untreated pwMS (MS-Un), individuals with other neurological disease (OND) and healthy controls (HC), and cohort 2 comprised natalizumab-treated pwMS (MS-Nat) and HC; viability of thawed PBMC were comparable between cohorts (Figure 1C). Natalizumab-treated pwMS were chosen due to elevated levels of autoreactive T cells in peripheral blood^18,41^.

Whilst responses to anti-CD3, CMV pp65 and ABD beads in cohort 1 were not significantly changed between MS-Un and other groups (Figure 1B and S2A), ANO2_79-168_-specific T cell responses were significantly increased in MS-Un donors, as measured by higher frequency of cells producing IFNγ, IL-17A and IL-22 (Figure 1B). Overall, 56.7% of MS-Un donors had a T cell response to ANO2_79-168_, defined as >10 IFNγ-secreting cells per 2.5×10^5^ PBMC. We observed particularly increased IL-17A^+^ and IL-22^+^ T cell responses to ANO2_79-168_ in MS-Nat individuals from cohort 2 compared to MS-Un and HC, suggesting that natalizumab treatment may lead to accumulation of these cells in the periphery (Figure 1C). We also observed a slight increase in the number of ABD and CMV pp65-specific T cells in MS-Nat compared to HC (Figure 1C and S2B).

We performed receiver operating characteristic (ROC) tests based on IFNγ, IL-17A and IL-22 responses in MS-Un and control donors (HC + OND) from cohort 1 (Figure 1D). Analysis of cytokine responses to ANO2_79-168_ produced an area under the curve (AUC) of 0.77 for IFNγ (CI 0.64-0.90, *P*=0.0002), 0.75 for IL-17A (CI 0.63-0.87, *P*=0.0005) and 0.85 for IL-22 (CI 0.74-93, *P*<0.0001), respectively (Figure 1D). ROC for ANO2_79-168_ T cell responses in cohort 2 (MS-Nat and HC) also showed high ROC values for all cytokines (Figure S2C), indicating a potential value of ANO2-specific T cell detection in MS diagnostic workup alongside other measures.

### T cell responses to other regions of ANO2 are elevated in pwMS and are predominantly CD4^+^IFNγ^+^

Previous studies have identified amino acid sequence homology between ANO2_140-149_ and EBNA1_431-440_ and cross-reactive antibodies which target both epitopes in MS^15^. To validate results in Figure 1B and to investigate whether T cell responses to ANO2 were restricted to the region with homology to EBNA1, we produced recombinant ANO2_1-275_, ANO2_409-525_ and ANO2_750-1003_ protein-coupled beads (Figure S1A and S1B)^40^, omitting transmembrane domains due to the difficulty of producing these regions in bacteria. EBNA1_1-120_ and EBNA1_380-641_ were designed to exclude the glycine-alanine repeat region which is also difficult to produce recombinantly. T cell reactivities to ANO2 and EBNA1 were tested by FluoroSpot.

We detected T cell responses to ANO2_1-275_, ANO2_409-525_, ANO2_750-1003_, which were higher in MS-Un and MS-Nat compared to OND and HC donors (Figure S3A-D). IFNγ T cell responses to EBNA1_1-120_ and EBNA1_380-641_ were similarly elevated in MS-Un and MS-Nat groups compared to controls. T cell responses to ANO2, EBNA1 and CMV pp65 were highly heterogeneous with individuals displaying unique autoreactivity profiles (Figure S3E-G). Responses were partially blocked by both HLA-DR and HLA-A/B/C blocking antibodies, suggesting that both CD4^+^ and CD8^+^ T cell responses towards EBNA1 and ANO2 were present (Figure S4). Due to sequence homology and previously reported antibody cross-reactivity^15^, we analysed T cell responses to ANO2 and EBNA1 in individuals and found these to be positively correlated, whereas no correlation was observed for CMV pp65 (Figure 1E).

Intracellular cytokine staining (ICS) and flow cytometry analysis of ANO2, EBNA1, CMV pp65 and anti-CD3-stimulated T cells were performed in a subset of MS-Nat and HC individuals, with example staining from two representative donors shown in Figure S5A-E. ICS confirmed results from FluoroSpot (Figure 1E, 1F, S6A and S6B), and overall there was a trend towards higher EBNA1 responses and significantly more CD4^+^IFNγ^+^ T cells after ANO2 stimulation in MS-Nat individuals (Figure 1F). Correlation of T cell responses to each target measured by flow cytometry (Figure S7) reflected findings from FluoroSpot (Figure 1E).

### Epitope mapping of ANO2_1-275_ reveals targeting of multiple epitopes

To understand if T cell responses were restricted to ANO2_140-149_ – the epitope with sequence homology to EBNA1^15^ – we mapped responses in MS-Nat individuals using an overlapping ANO2_1-275_ peptide library. Briefly, T cell responses to 66 peptides (15-mers overlapping by 11aa) covering the ANO2_1-275_ region (divided into six pools) were analysed by FluoroSpot to understand which regions were immunodominant.

Whilst responses to peptides were overall lower than responses to antigen-coupled beads, we noted multiple responses to ANO2 peptide pools with no homology to EBNA1 (Figure S8A). We also identified several donors who responded to >1 peptide pool, which may indicate intramolecular epitope spreading within ANO2_1-275_. However, as we only analysed one timepoint, it is not possible to deduce if these responses developed simultaneously or sequentially (Figure S8A). One donor (P68) demonstrated responses to multiple ANO2 peptides when responses to individual peptides were tested by FluoroSpot, including regions spanning aa1-15, aa9-35, aa33-47, aa41-59, aa53-67, aa61-79 and aa73-87 (Figure S8B). Although IFNγ^+^ spots were overall fewer than those detected for TNFɑ and Granzyme B, T cell responses seemed to show a different pattern of cytokine production across regions of ANO2.

### Antibody and T cell responses to ANO2 are moderately correlated in pwMS

Antibody responses to ANO2 and EBNA1 epitopes were analysed in a cohort of MS and control individuals (MS n=713, Con n=722) as previously described^18^, and CRYAB was included as another MS autoantigen with sequence homology to EBNA1^18^ (Table S2). Antibody responses to ANO2_134-153_ with homology to EBNA1_430-440_ were significantly higher in pwMS, confirming previous studies^15,23^ (Figure S9A). Anti-EBNA1 IgG towards multiple regions including aa69-88, aa391-410, aa393-412, aa401-420 and aa425-444 were higher in MS, as well as CRYAB IgG to aa2-16 and aa3-17 (Figure S9A). Antibody and T cell responses to selected ANO2 and EBNA1 peptides were correlated in a subset of individuals (MS-Nat n=33) showing that ANO2_79-168_ IFNγ^+^ T cell responses (FluoroSpot) are moderately correlated with anti-ANO2_135-149_ IgG, the region of ANO2 with homology to EBNA1_430-440_ (Figure S9B and S9C). Although representing a small subset of donors, a significant correlation between ANO2 T cell and antibody responses to the same region nevertheless indicates that ANO2 IgG responses may be used to enrich for donors displaying pathogenic T cell responses.

### ANO2_1-275_ and EBNA1_380-641_ immunisation cross-primes CD4^+^ T cell responses to the reciprocal antigen *in vivo*

To explore cross-reactivity of EBNA1 and ANO2 adaptive immune responses *in vivo*, we immunised three different mouse strains – C57BL/6, NOD and SJL/J – with ANO2_1-275_ protein, EBNA1_380-641_ protein or adjuvant only. After 9-11 days, inguinal draining lymph nodes were harvested and single-cell suspensions restimulated *in vitro* for antigen reactivity to bead-bound antigen, protein and/or peptides by flow cytometry and ICS^18,40^ (gating strategies in Figure S10A-C). In-house ELISAs were optimised to detect EBNA1 and ANO2-specific IgG in mouse sera (Figure S11A-C).

ANO2_1-275_ T cell priming was absent in C57BL/6 mice, but some reactivity to EBNA1_380-641_ was observed in the CD4^+^IFNγ^+^ T cell compartment (Figure S12A). In contrast, NOD mice demonstrated significant CD4^+^ T cell priming to both EBNA1_380-641_ and ANO2_1-275_ after immunisation and tended to show cross-reactive T cell responses to homologous antigens but not to autoantigens with no sequence homology, although this did not reach significance (Figure S12B).

Immunisation of SJL/J mice with ANO2_1-275_ or EBNA1_380-641_ induced strong CD4^+^ T cell responses to both antigens with evidence of antigen cross-priming on the CD4^+^ T cell level *in vivo* (Figure 2A). Surprisingly, ANO2_1-275_-primed SJL/J mice showed CD4^+^IFNγ^+^ T cell responses not only to EBNA1_380-641_, but also to EBNA1_1-120_, which lacks cross-reactive residues deducible from the amino acid sequence only. However, this reactivity did not comprise other MS-associated autoantigens; synaptosomal-associated protein 91 (SNAP91), synuclein-β (SNCB)^53^, RAS guanyl-releasing protein 2 (RASGRP2)^54^, GDP-L-fucose synthase (GDPLFS)^55^ or glial cell adhesion molecule (GlialCAM)^16^ (Figure 2A). Similarly, EBNA1_380-641_-immunised mice had cross-reactive T cell responses to ANO2_1-275_ but not to other autoantigens (Figure 2A).

To dissect EBNA1 and ANO2 cross-priming of CD4^+^ T cell responses further, we repeated immunisation experiments of SJL/J mice with HCMV IE1162-173 peptide (irrelevant antigen control), human ANO2_137-151_ peptide that contains the EBNA1 cross-reactive sequence, ANO2_1-275_ peptide pool, ANO2_1-275_ protein, EBNA1_428-444_ peptide that contains the ANO2 cross-reactive sequence, EBNA1_380-641_ peptide pool and EBNA1_380-641_ protein. Splenocytes from immunised mice were subsequently rechallenged i*n vitro* with antigens used for the original immunisations as well as to all the other immunogens described above. EBNA1_380-641_ and ANO2_1-275_ protein-immunised animals demonstrated a strong T cell priming, both to the immunisation antigens as well as significantly increased T cell responses to the reciprocal antigen (Figure S12D). Interestingly, mice immunised with EBNA1_380-641_ protein also had significantly higher T cell responses to mouse ANO2_137-151_ peptide and to the EBNA1_428-444_ peptide (Figure S12D). These results indicate that EBNA1 protein immunisation elicits cross-reactive T cell responses both to full length ANO2_1-275_ protein and also to the peptide with homology to EBNA1, ANO2_137-151_. Immunisation of mice with the minimal EBNA1_428-444_ peptide with homology to ANO2_137-151_ demonstrated low but specific reactivity to human and mouse ANO2_137-151_ peptides, as well as to full length EBNA1_380-641_ and ANO2_1-275_ proteins (Figure 2K).

Similarly, mice immunised with ANO2_1-275_ protein demonstrated T cell priming to the original stimulus as well as to both the ANO2_1-275_ peptide pool and the EBNA1_380-641_ protein (Figure S12D). Likewise, ANO2_1-275_ peptide pool-immunised mice had significantly increased responses to the original stimulation as well as to ANO2_1-275_ protein and EBNA1_380-641_ peptide pool and protein demonstrating cross-reactivity (Figure S12D). Mice immunised with the minimal human ANO2_137-151_ peptide with homology to EBNA1 had significantly higher T cell responses to the original stimulus and to ANO2_1-275_ and EBNA1_380-641_ proteins, and non-significant increased reactivity to mouse ANO2_137-151_ peptide and EBNA1_428-444_ peptide (Figure 2K). These results confirm previously demonstrated cross-priming (Figure 2A and 2J) and also show that immunisation with EBNA1_428-444_ peptide can lead to cross-reactive T cell responses to its homologous ANO2_137-151_ peptide (both mouse and human).

The lower magnitude of priming, recall responses *in vitro* and cross-reactive responses against the minimal epitopes as compared to full proteins and peptide pools suggested that additional immunodominant epitopes might serve as response drivers. We mapped responses to smaller peptide pools throughout the EBNA1_380-641_ region (Figure S12E). The immunodominant response from EBNA1-immunised animals targeted EBNA1_468-522_, which does not contain the EBNA1/ANO2 homologous region identified as a cross-reactive antibody target (EBNA1_430-440_). This indicates that, although T cell responses to the EBNA1_430-440_ region can be elicited in SJL/J mice (Figure 2K), the EBNA1_468-522_ region is immunodominant in this mouse strain. However, we cannot exclude that our epitope mapping approach with a 15-mer peptide library misses the optimal sequence presented from natural trimming of EBNA1 by endogenous proteases. Overall, immunisation of two different mouse strains – NOD and SJL/J – led to cross-priming of CD4^+^ T cell responses to ANO2_1-275_ and EBNA1_380-641_ and indicates cross-reactivity between these antigens *in vivo*.

### ANO2_1-275_ pre-immunisation cross-primes EBNA1-reactive T cell and antibody responses and leads to a more severe EAE phenotype

SJL/J mice are known to demonstrate stronger T cell priming to multiple classical myelin MS autoantigens including MOG, MBP and PLP, as well as displaying increased susceptibility to EAE as compared to the C57BL/6 strain^56^. We therefore selected the SJL/J strain to investigate if ANO2-immunisation would affect the course of EAE. Eight-week-old SJL/J mice were pre-immunised with either an irrelevant protein (ABD) or ANO2_1-275_ protein in combination with CpG1826 in incomplete Freund’s adjuvant (IFA). After 3 weeks, EAE was induced by injection of PLP in complete Freund’s adjuvant (CFA), and the ensuing clinical phenotype was followed for up to 55 days (Figure 2B).

Immunised mice from both groups presented with a classical EAE phenotype characterised by ascending paralysis, but ANO2_1-275_/PLP-immunised mice had more severe disease (Figure 2C) with significantly higher maximum EAE scores, longer disease duration, elevated cumulative disability scores and trends towards earlier disease onset and a higher number of relapses (Figure 2D and Figure S12C). Higher disease scores in ANO2_1-275_/PLP-immunised mice may be attributed to significantly higher CD4^+^ T cells counts and trends towards higher neutrophils and monocytes/macrophages in the spinal cord (Figure 2E). We also ran experiments on mice immunised solely with ANO2, but these mice presented with no EAE disease phenotypes, nor overt observable behavioural changes (Figure 3A).

To explore underlying mechanistic differences in EAE phenotype in greater detail, eight mice from each group were sacrificed at the peak of relapse (around day 30), and spinal cord-infiltrating T cells were analysed by flow cytometry. Spinal cord CD4^+^ T cells from ANO2_1-275_/PLP-immunised mice had greater numbers of FoxP3^+^, IL-17A^+^ and Ki67^+^ cells than ABD/PLP-immunised mice (Figure 2F).

To analyse antibody responses, sera from animals were tested for IgG reactivity to ANO2_1-275_ and EBNA1_380-641_ (optimisation for ELISAs is shown in Figure S11). ANO2_1-275_/PLP-immunised mice demonstrated antibody responses to ANO2_1-275_ at induction of EAE (priming) that were maintained throughout the experiment up to day 50 (Figure 2G). EBNA1_380-641_ cross-reactive IgG antibodies were only found in ANO2_1-275_/PLP-immunised mice and not in the ABD/PLP group. In addition, emergence of EBNA1_380-641_-specific IgG in ANO2_1-275_/PLP-immunised mice was delayed to day 30 (Figure 2G) and required lower serial dilution (1:6000) for detection. A strong correlation was also observed between ANO2_1-275_ and EBNA1_380-641_ IgG (Figure S13A) at day 30. Interestingly, there were non-significant negative trends between EAE scores and ANO2_1-275_ IgG at sacrifice on day 30 and day 50 (Figure S13B) and a significant positive correlation between CD4^+^ T cell counts and EAE scores at day 50 (Figure S13C). Correlation matrices of clinical and immunological factors revealed that high EAE scores were associated mostly with T cell factors – but not B cells or autoantibodies – at both the peak (day 30) and the chronic phase (day 50) of disease in ANO2_1-275_/PLP-immunised mice (Figure S13D). Together these data demonstrate that EAE is exacerbated by ANO2_1-275_ CD4^+^ T cells but suggest that ANO2_1-275_ antibodies are not necessary for disease in this model.

### Protein versus peptide ANO2_1-275_ pre-immunisation demonstrates that CD4^+^ T cell responses are responsible for increased EAE severity

We compared ANO2_1-275_ protein and peptide pool (15-mers overlapping by 11 amino acids) pre-immunisation in the same SJL/J model (Figure 2H). As before, mice pre-immunised with either ANO2_1-275_ protein or peptides had higher EAE scores throughout the experiment and higher AUC values at day 31 than ABD pre-immunised mice, indicating a more severe disease course (Figure 2H).

Remarkably, despite similar disease severity and course, mice pre-immunised with ANO2_1-275_ peptide pool had no detectable ANO2_1-275_-specific antibodies in sera at day 31 nor did they respond to EBNA1_380-641_ (Figure 2I). In contrast, mice pre-immunised with ANO2_1-275_ protein had ANO2_1-275_ IgG responses and cross-reactive EBNA1_380-641_ antibodies (Figure 2I). IgG responses to ABD, but not ANO2_1-275_ and EBNA1_380-641_, could be detected in the ABD group. In contrast, splenic CD4^+^ T cell responses to ANO2 could be detected in mice immunised either with ANO2_1-275_ protein or peptides by flow cytometry, with evidence of cross-priming of EBNA1_380-641_-specific T cells in the ANO2_1-275_ protein group (Figure 2J). ABD pre-immunised mice showed CD4^+^ T cell responses to ABD beads but not to ANO2_1-275_ or EBNA1_380-641_, and no significant differences were observed in reactivity to MOG or PLP between groups (Figure 2J).

We further investigated the minimal epitope needed to affect EAE disease course by pre-immunising mice with ANO2_137-151_ and EBNA1_428-442_ peptides (Figure 2B). A trend towards higher EAE scores at disease onset was observed for ANO2_137-151_ peptide pre-immunised animals compared to control or EBNA1_428-442_ groups (Figure S14A) although AUC for EAE scores did not reach significance (Figure S14B). These data indicate that pre-immunisation with the minimal epitope ANO2_137-151_ may induce more severe disease, however pre-immunisation with ANO2_1-275_ protein led to the highest EAE scores, likely due to stronger and/or broader induction of autoreactive CD4^+^ T cell responses *in vivo* as suggested by our T cell priming data comparing minimal peptides, peptide pools and recombinant proteins (Figure 2K and S12D-E).

Given the EBNA1 homology with ANO2 and GlialCAM^16^, we wanted to understand if pre-immunisation with other antigens would likewise lead to a more severe EAE phenotype. Using the same approach (Figure 2B), SJL/J mice were pre-immunised with recombinant ABD, GlialCAM_262-418_ protein, EBNA1_380-641_ protein or EBNA1_380-641_ overlapping peptides. Mice pre-immunised with EBNA1_380-641_ protein or peptides had similar EAE scores and were higher than those observed for ABD or GlialCAM_262-418_ pre-immunised animals (Figure S14C and E) but the AUC for EAE scores at both day 30 as well as day 51 were not significantly different between groups (Figure S14D and F). A trend towards more severe disease and higher AUC in EBNA1_380-641_ protein or EBNA1_380-641_ overlapping peptide pool pre-immunised mice was observed compared to the ABD group up to 31 days post-immunisation (Figure S14F), and EBNA1_380-641_ protein pre-immunised mice developed antibody responses to both EBNA1_380-641_ and ANO2_1-275_ at priming which were maintained for 50 days post EAE induction (Figure S14G). These data suggest that EBNA1 pre-immunisation also leads to more severe EAE than with GlialCAM or ABD, possibly due to homology with several autoantigens contributing to worsened CNS autoimmune disease.

### ANO2_1-275_ pre-immunisation of adult mice gives an atypical EAE phenotype and increases the number of activated microglia and macrophages in the CNS

Given a recent report that adoptive transfer of T_H_17 T cells in EAE can give rise to an atypical clinical phenotype in adult mice (8-15 months old)^57^, we also investigated if age would impact the clinical phenotype following ANO2_1-275_ pre-immunisation. Six-month-old SJL/J mice – at the developmental stage equivalent to around 30 human years – were pre-immunised with either adjuvant only or adjuvant plus ANO2_1-275_. After 21 days EAE was induced by PLP immunisation (Figure 2B).

By 20 days after EAE induction, most mice had reached their humane endpoint and were sacrificed leading the experiment to end prematurely. ANO2_1-275_/PLP-immunised adult animals showed increased incidence, had more severe EAE scores than the control/PLP-immunised group (Figure 3A and 3B) and decreased survival (Figure 3C). In addition, twice as many mice in the ANO2_1-275_/PLP group compared to the control displayed atypical EAE phenotypes characterised by descending paralysis, incoordination, balance disturbance and seizure-like spasms (Figure 3D, Videos S1-3). Immunofluorescence staining analysis of CNS tissue revealed trends towards increased spinal cord and brain demyelination in the ANO2_1-275_/PLP-immunised group and significantly increased total CNS demyelination (Figure S14H).

Immunofluorescence staining for nuclei (DAPI), myelin (MBP) and activated macrophages/microglia (ionised calcium-binding adaptor molecule 1, Iba1) of brain and spinal cord of adult mice was performed to understand whether inflammation was localised to certain brain regions which may account for the higher proportion of atypical EAE. Adjuvant/PLP-immunised animals showed overall weak Iba1 staining across the brain. In contrast, the ANO2_1-275_ pre-immunised group showed activated microglia/macrophages in several regions, as demonstrated by significantly increased Iba1 staining in the hippocampus (Figure S14I and J).

Further immunofluorescent staining of adjuvant/PLP-immunised mouse brains with DAPI, and with anti-ANO2 and anti-podocalyxin antibodies showed that ANO2 expression was mostly localised to vessel epithelial cells overlapping with podocalyxin (Figure S14K). In contrast, vascular ANO2 expression seemed weaker and relatively absent on endothelial cells in the ANO2_1-275_/PLP-immunised group (Figure S14K). CD4 staining of hippocampi demonstrated T cells which were located next to vessels in ANO2_1-275_/PLP-immunised mice indicating immune infiltration (Figure S14L). Overall, these data show that a higher proportion of ANO2_1-275_ pre-immunised mice have an atypical EAE phenotype which could be driven by increased activated microglia/macrophages in the hippocampus and demyelination in the brain stem.

### ANO2 co-localises with CD31 staining in the vascular endothelium and is expressed in the cerebellum, inferior olivary nucleus and septum of mouse brains

Expression of *Ano2* in mice is not well-characterised in the literature, since bulk and scRNAseq are often biased by cell type and region, as well as efficiency of isolation of certain cell types, a similar pitfall as for *ANO2* expression in human samples. To understand how ANO2 autoimmunity in EAE exacerbates disease and leads to an atypical phenotype, we investigated RNA and protein expression of ANO2 in mouse brains by RNAscope and immunolabeling-enabled three-dimensional imaging of solvent-cleared organs (iDISCO) respectively.

Whole brains stained with Ano2-specific probes and Ano2 transcripts were detected by RNAscope in three main areas: the septum, the inferior olivary nucleus and in the cerebellum (Figure 3E). These results were in concordance with Ano2 regional expression in these sites of the mouse brain reported in the Allen Brain Atlas (https://mouse.brain-map.org/) (Figure S15H).

To verify that protein could also be detected in these areas, we applied iDISCO-based whole-mount imaging to study ANO2 protein expression in mouse brains. This method involves tissue clearing and whole mount immunolabelling, enabling detailed analysis of both antigen expression and their co-localisation with other markers overlayed onto intact brain architecture. Results reflected what we observed in RNAscope findings, with several foci of ANO2 protein expression in the mouse brain in the areas of the cerebellum, inferior olivary nucleus and the septum (Figure 3F-K and S15A-G). Imaging of ANO2 demonstrated a clear vasculature-like staining pattern (Figure 3I) and, when overlayed with the classical endothelial marker CD31, co-expression of these markers was 76% in the cerebellum (Figure 3K, Video S4), 85% in the septal region (Figure S15E, Video S5) and 72% in the inferior olivary nucleus (Figure S15G, Video S5). Therefore, in regions of the adult mouse brain where ANO2 was highly expressed, ANO2 appears to be expressed mostly in association with the vasculature.

These findings demonstrate that ANO2 expression in the mouse brain is not uniform. Given the higher incidence of atypical EAE disease in ANO2_1-275_ pre-immunised mice (Figure 3D), the atypical phenotype may be driven by anti-ANO2 responses directed against the vasculature or adjacent cells in the cerebellum, inferior olivary nucleus and septal regions.

### Adoptive transfer of ANO2_1-275_-specific T cells leads to greater plasma extravasation in areas of high ANO2 expression

To understand whether ANO2-specific CD4^+^ T cells could exacerbate EAE in the context of PLP-induced EAE, we performed adoptive transfer of either ANO2_1-125_-specific T cells or HCMV IE1 antigen-specific control T cells together with T cell specific for PLP_139-151_ peptide (Figures 4A and 4B). Animals that developed disease were aligned by day of onset, and the results clearly replicated our previous findings of disease exacerbation following ANO2_1-125_ pre-immunisation (Figures 2 and 3). There were no differences in EAE scores between mice adoptively transferred with PLP T cells or PLP+HCMV-specific T cells (Figure S15J).

**Figure 4.**
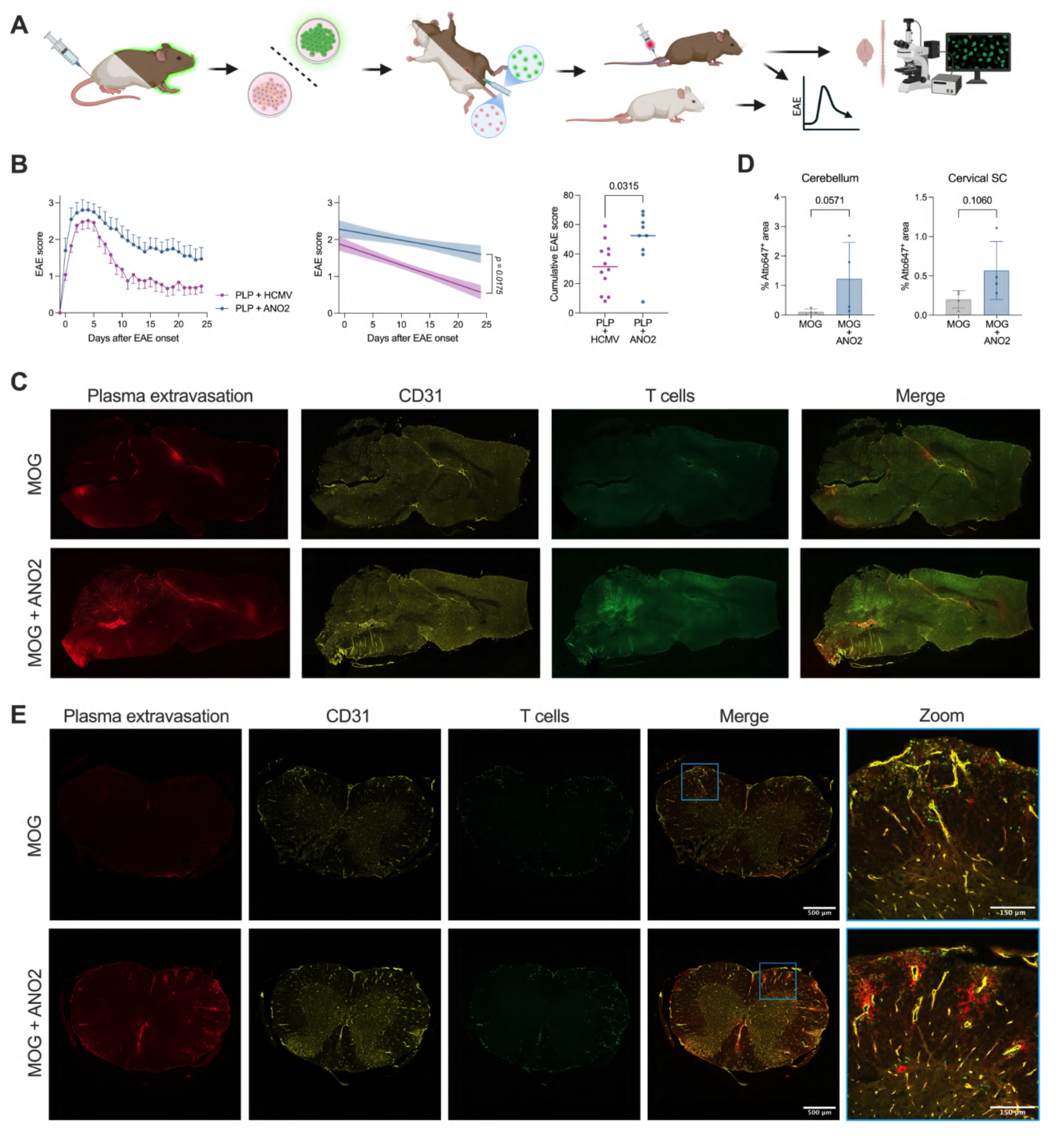
Adoptive transfer of ANO2-specific T cells leads to extravasation in areas of the mouse brain where ANO2 is expressed. **A.** Schematic of adoptive T cell transfer experiments. (TOP) SJL/J mice (white) were immunised with PLP_139-151_ peptide, HCMV IE1_162-173_ peptide or ANO2_1-125_ peptide pool. After 9 days spleens and draining inguinal lymph nodes were collected and cells were restimulated *in vitro* with the immunisation peptides in the presence of IL-23 to expand antigen-specific T cells. After 3 days, PLP only, PLP + HCMV, or PLP + ANO2-specific T cells were adoptively transferred into recipient SJL/J mice and the disease course was followed for up to 30 days. (BOTTOM) Schematic of adoptive T cell transfers followed by plasma extravasation and *ex vivo* tracking of T cells by immunofluorescence imaging. SJL/J x C57BL/6 F1 cross animals with ubiquitous expression of GFP (brown/green) were immunised with MOG_35-55_ or ANO2_1-125_. After 9 days spleens and draining inguinal lymph nodes were collected and cells were restimulated *in vitro* with the immunisation peptides in the presence of IL-23 to expand antigen-specific T cells. After 3 days, MOG only or MOG + ANO2-specific T cells (all GFP positive) were adoptively transferred into GFP-negative SJL/J x C57BL/6 F1 cross recipient mice (brown). After 11 days at disease onset, mice were anaesthetised and injected with Atto647-labelled plasma which was perfused through the animals. Mice were subsequently sacrificed and their brains and spinal cords analysed for CD31 expression, plasma extravasation and the presence of GFP-positive T cell infiltration. **B.** EAE scores of mice adoptively transferred with T cells specific for PLP only (n=3), PLP+HCMV (n=5) or PLP+ANO2 (n=3) over 30 days post transfer (left panel). Given the serendipitous EAE onset, scores were plotted by day of onset. Linear regression curve of EAE scores of PLP only, PLP+HCMV or PLP+ANO2 in adoptive T cell transfer mice. The shaded area represents the 95% confidence interval (middle panel). Cumulative EAE scores of adoptively transferred mice, statistical test is the Welch T test (right panel). **C.** Whole brain immunofluorescent imaging of mice that received adoptive transfer of MOG T cells only or MOG + ANO2-specific T cells (all GFP-positive) as described in **A** (brown mice). Plasma extravasation was most pronounced in the cerebellum and in regions surrounding the olivary nucleus and the septum. CD31 was used as a marker for vascular endothelium. **D.** Quantification of the Atto647^+^ area in the cerebellum (left) and cervical spinal cords (right) of mice receiving adoptive transfer of MOG only (n=4) or MOG + ANO2 T cells (n=4). The atto-647^+^ area indicates plasma extravasation. Statistical test is the Mann-Whitney. **E.** Immunofluorescent staining of spinal cords of mice which have received adoptive T cell transfer of MOG only or MOG+ANO2 T cells (all GFP-positive) as described in **C**. Spinal cords were stained with CD31 to identify the vascular endothelium, and Atto-647 staining indicates plasma extravasation. Created with https://BioRender.com.

To further dissect the mechanism of action of these cells and track them in the CNS we used an F1 cross between SJL/J mice and C57BL/6 animals with ubiquitous expression of GFP as donors and non-GFP SJL/J x C57BL/6 F1 cross as recipients. These F1 mice develop EAE in response to both MOG_35-55_ and PLP_139-151_ peptide immunisation and, given that MOG-specific T cells are generally concentrated in the lower portions of the spinal cord and the optic nerve and give overall milder disease, we chose MOG in this context so that exacerbation of disease initiated by ANO2 T cells could be better distinguished. Given that expression of ANO2 could be observed associated to vasculature in mouse brains (Figure 3K), at EAE onset animals were additionally intravenously injected with fluorescently-labelled mouse plasma to assess blood-brain barrier (BBB) integrity followed by imaging of sections of the brain and spinal cord (Figure 4D-F).

Interestingly, animals transferred only with ANO2-specific T cells neither developed disease nor showed any signs of T cell infiltration or evidence of BBB leakage (Figure S15I); findings which recapitulated mice immunised with ANO2 protein alone which did not develop EAE (Figure 3A). MOG T cell-transferred animals on the other hand had a clear presence of both T cells and BBB leakage which was exacerbated by co-transfer of ANO2 T cells, specifically in the cervical spinal cord as well as in cerebellum (Figure 4D-F, Figure S15K). Immunofluorescent imaging of brains from mice which had received co-transfer of GFP^+^ ANO2-specific T cells showed that these were present both in the cerebellum and in the cervical region of the spinal cord (Figure S15L), in accordance to the sites where plasma extravasation (Figure 4D and 4F) and ANO2 expression (at least for the cerebellum) were previously observed (Figure 3E-K).

### ANO2_1-275_-specific T cells can directly target ANO2-expressing glial cell targets *in vitro*

To address whether ANO2-specific CD4+ T cells exert an effect on target cells expressing ANO2, we cultured neurons, OPCs, MOLs, astrocytes and microglia with ANO2-specific resting T cells in vitro. These CD4+ T cells could be reactivated by ANO2-derived peptides in two ways: either by direct presentation for those cells capable of MHC class II expression or by indirect presentation though APCs that could pick up cell debris containing ANO2. To allow for the latter, we thus included conditions in which different glial cells were cocultured in presence or absence of LPS-activated bone marrow-derived dendritic cells (DCs) (Figure S16B). As a positive control, ANO2_1-125_ peptides were added in separate wells containing the glial target, T cells and DCs. In vitro-cultured cells were assessed for *Ano2* expression by qPCR revealing expression for MOLs, followed by OPCs and astrocytes (Figure S16A).

Incucyte tracking of the target cells in the presence of a Caspase 3/7 activation dye revealed that MOLs were susceptible to T cell mediated killing, even in the absence of DCs or exogenous peptide, and that this effect was most prominent for MOLs after 24h of co-culture (Figure S16B). We have previously shown that OPCs and MOLs upregulate MHC class II in response to IFNγ and are able to present antigens to T cells^58^, an effect we confirmed with exogenous IFNγ stimulation on both OPC and MOL cells in vitro (Figure S16C). Increased but transient Caspase3/7 signal could only be detected early in co-cultures with astrocytes, neurons or microglia but did not lead to reduced target cell counts at later timepoints, suggesting this effect to be a transient response to cell stress^59,60^. Additionally, an increase in MOL and OPC cell numbers could be observed in cocultures with activated DCs and suggests that this interaction could potentially deliver tonic signals akin to pathways proposed by their interaction with other myeloid cells^61,62^.

In summary, our experiments show that ANO2-specific T cells are able to cause disease exacerbation in conjunction with myelin antigen-specific T cells. In addition, ANO2 T cells migrate to areas of the CNS that express ANO2, directly target ANO2-expressing cells in vitro and lead to increased BBB leakage.

### Dual-specific T cell clones with reactivity to ANO2 and EBNA1 are found in HLA-DRB1*15:01 carriers with MS

Given our findings that ANO2 and EBNA1 immunisation in mice can cross-prime CD4^+^ T cell responses to the reciprocal antigen (Figure 2A, 2J, 2K and S12D), we further investigated whether human cross-reactive ANO2_1-275_ and EBNA1_380-641_ T cells can be detected in pwMS by isolating and expanding T cell bulk (TCB) and T cell clones (TCC) responding to these antigens. Briefly, PBMC from MS-Nat individuals were subjected to CD45RA negative depletion (to remove naïve T cells and B cells), labelled with CFSE and stimulated with bead-bound ANO2_1-275_ or EBNA1_380-641_ antigens (Figure 5A-B)^63^. After 8 days, responding cells were sorted and cultured for a further 2 weeks, and the resulting TCC were screened for reactivity to EBNA1 and ANO2 by FluoroSpot (Figure S17A and B).

**Figure 5.**
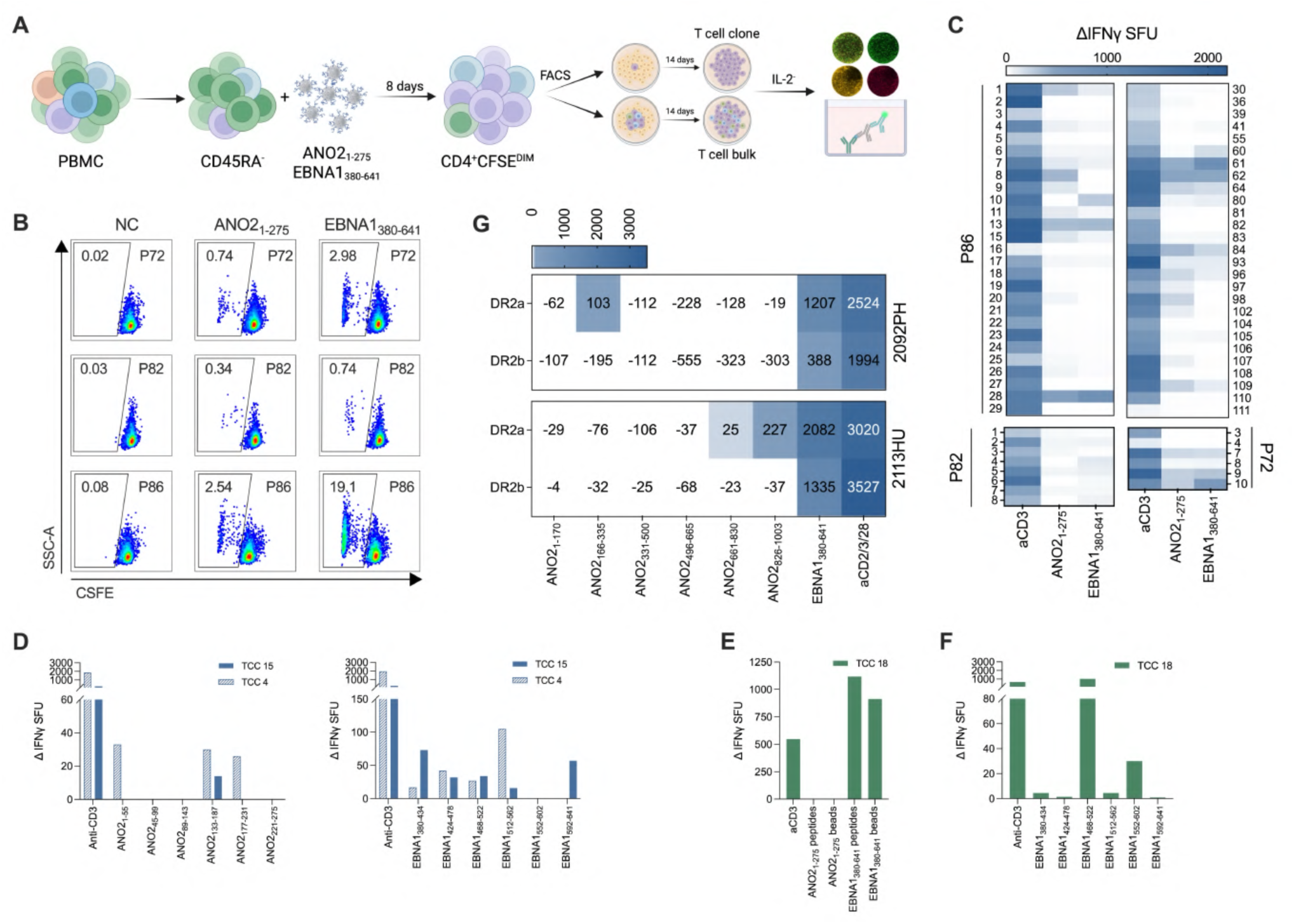
ANO2-specific CD4^+^ T cells respond to EBNA1 in MS. **A.** Schematic of expansion ANO2_1-275_ and EBNA1_380-641_-specific T cells from PBMC of pwMS. Briefly, PBMC from MS-Nat were subjected to CD45RA-negative selection, labelled with CFSE and stimulated with antigen-coupled beads. After 8 days, cells were surface stained for CD3 and CD4 expression and responding LiveCD3^+^CD4^+^CFSE^DIM^ T cells were sorted by FACS. For T cell clones (TCC), single cells were sorted into wells of a tissue culture plate containing irradiated allogeneic feeder PBMC plus PHA and IL-2. For T cell bulk (TCB) lines, 100 cells/well were sorted into plates containing irradiated allogeneic feeder PBMC plus PHA and IL-2. Sorted cells were expanded in culture for 14 days, and IL-2 was withdrawn for 5 days prior to testing for antigen specificity by IFNγ FluoroSpot or ELISpot. **B.** Proliferating LiveCD3^+^CD4^+^CFSE^DIM^ T cell populations after 7 days’ stimulation with negative control (NC) beads, ANO2_1-275_ and EBNA1_380-641_ beads from three representative HLA-DRB1*15:01^+^ MS-Nat donors: P72, P82 and P86. Percentage of CFSE^DIM^ cells of the total LiveCD3^+^CD4^+^ population indicated in the top left corner of gates. **C.** T cell clones generated from ANO2_1-275_-stimulated LiveCD3^+^CD4^+^CFSE^DIM^ cells from MS-Nat donors P72, P82 and P86 were screened for reactivity to ANO2_1-275_ and EBNA1_380-641_ beads by IFNγ FluoroSpot, anti-CD3 was used as a positive control. Data presented as background corrected IFNγ SFU by subtracting SFU for the negative control (NC) beads from anti-CD3, EBNA1 and ANO2 bead stimulations. T cell clones showed a range of specificities with some responding to ANO2_1-275_ or EBNA1_380-641_ or both targets, indicating that ANO2_1-275_ T cell can also respond and produce IFNγ in response to EBNA1_380-641_. **D.** TCC 4 and TCC 15 from a HLA-DRB1*15:01^+^ natalizumab-treated pwMS (P56) were expanded following stimulation with the ANO2_1-275_ overlapping peptide pool. TCC were re-challenged with an overlapping peptide library (15-mers overlapping by 11aa) covering ANO2_1-275_ (left panel) and EBNA1_380-641_ (right panel) in a FluoroSpot assay. TCC 4 and TCC 15 both responded to multiple peptide pools from ANO2 and EBNA1. **E.** TCC 18 was generated from a HLA-DRB1*15:01^+^ natalizumab-treated pwMS (P56) following stimulation with ANO2_1-275_ overlapping peptide pool. When screened for reactivity, TCC18 did not respond to ANO2_1-275_ beads or peptide, but did respond to EBNA1_380-641_ beads and peptide. **F.** TCC 18 was re-stimulated with an overlapping peptide library (15-mers overlapping by 11aa) covering EBNA1_380-641_ (right panel) in a FluoroSpot assay, and was found to respond to EBNA1_468-522_. **G.** T cell bulk (TCB) lines from two untreated HLA-DRB1*15:01^+^ pwMS (top panel: donor 2092PH and bottom panel: donor 2113HU) were generated from CFSE^DIM^ (proliferating, responding) and CFSE^HI^ (non-proliferating, non-responding) sorted after EBNA1_380-641_ stimulation were screened for ANO2 reactivity by IFNγ ELISpot using overlapping 20-mer peptide pools covering the entire sequence of ANO2 (aa1-1003). Reactivity to an overlapping EBNA1_379-641_ peptide pool was also used to establish the specificity of the T cell bulk line. Anti-CD2/3/28 beads were used as a positive control and BLS cells expressing DR2a and DR2b were used as antigen presenting cells. Data is presented as background subtracted ΔIFNγ SFU using the BLS DR2a and BLS DR2b no peptide control conditions. Data presented as background subtracted ΔIFNγ SFU, negative values presented as 0. T cell clones (TCC), spot-forming units (SFU). \**P* < 0.05; \*\**P* < 0.01; \*\*\**P* < 0.001; \*\*\*\**P* < 0.0001. Created with https://BioRender.com.

Multiple TCC with reactivity to both ANO2_1-275_ and EBNA1_380-641_ were isolated from donors P72 and P86, carriers of the risk *HLA-DRB1*15:01* haplotype, indicating that there is cross-reactivity on the single cell level between these antigens (Figure 5C); the same experiment was repeated with the same result (Figure S19). A range of responses was also demonstrated, with some having a higher number of IFNγ-secreting cells when stimulated with ANO2 than EBNA1 and vice versa (Figure 5C). These findings support our hypothesis that some EBNA1-specific CD4^+^ T cells may contribute to MS pathology by targeting ANO2 *in vivo*.

TCC4 and TCC15 were generated from a *HLA-DRB1*15:01^+^* natalizumab-treated pwMS (P56) following stimulation with the ANO2_1-275_ peptide pool. When the epitopes were mapped against overlapping ANO2 and EBNA1 peptide libraries, both were found to respond to multiple pools from ANO2_1-275_ and EBNA1_380-641_, suggesting that TCC4 and TCC15 cross-react with multiple epitopes within both antigens (Figure 5D). TCC18 was also generated from P56 in the same T cell cloning following stimulation with the ANO2_1-275_ peptide pool. However, TCC18 did not respond to the original stimulation when screened but responded strongly to EBNA1_380-641_ beads and peptides (Figure 5E). When responses were deconvoluted using the EBNA1 overlapping peptide library, TCC18 was found to respond strongly to EBNA1_468-522_ and to a lesser extent EBNA1_552-602_ (Figure 5F). This is similar to what was observed for ANO2-immunised SJL/J mice (Figure S12D).

Next, we screened polyclonal EBNA1-specific CD4^+^ TCB lines from two *HLA-DRB1*15:01* carriers for specificity to 20-mer peptide libraries spanning the full sequence of ANO2 (amino acids 1-1003) using BLS cells expressing DR2a (*HLA-DRB5*01:01*) or DR2b (*HLA-DRB1*15:01*) as antigen-presenting cells (APCs). Both TCB lines showed high reactivity to the original antigen stimulus EBNA1_380-641_ but displayed reactivity to different parts of ANO2. EBNA1_380-641_-specific TCB from donor 2092PH was found to have DR2a-restricted reactivity to peptides spanning ANO2_166-335_ (Figure 5G). In contrast, EBNA1_380-641_ TCB from donor 2113HU had DR2a-restricted reactivity to ANO2_826-1003_ and to a lesser extent ANO2_661-830_ (Figure 5G), indicating that other regions of full-length ANO2 may be targeted by DR2a-restricted EBNA1-specific CD4^+^ T cells. This experiment was repeated for consistency and demonstrated similar results (Figure S19A), also aligning with the observation that ANO2-immunised SJL/J mice respond to regions of ANO2 without clear amino acid sequence homology (Figure S12D).

T cell clones (TCC1-100) generated from a pwMS (2113HU) with reactivity to both EBNA1 and ANO2 were also screened for reactivity to both EBNA1_380-641_ and ANO2_79-168_ beads by incubating with autologous PBMC in an IFNγ ELISpot assay (Figure S19B). TCC showed a range of reactivities, but several T cell clones clearly responded to both EBNA1_380-641_ and ANO2_79-168_ beads. For cross-reactive TCC, higher reactivity was almost always observed for EBNA1 than for ANO2, suggesting that these cells have greater avidity for EBNA1 than for ANO2 and responses may have originally been stimulated *in vivo* by viral antigen (Figure S19B).

Overall, our data indicate that T cells with reactivity to ANO2 can be readily detected in the EBNA1-specific CD4^+^ compartment of pwMS. Data from donors 2092PH and 2113HU also suggest that these responses may be restricted by *HLA-DRB1*15:01* and *HLA-DRB5*01*01*, and that cross-reactivity between these repertoires exists outside of the ANO2 region with strict amino acid sequence homology to EBNA1, findings which are similar to what we observed in SJL/J mice (Figure S12D). These data provide direct evidence that EBV infection primes T cell responses with the ability to target the MS-associated autoantigen ANO2 in humans.

### Single cell TCR sequencing of EBNA1- and ANO2-reactive CD4^+^ T cells reveals a significant overlap between TCR repertoires

To investigate the extent of overlap between EBNA1 and ANO2-specific CD4^+^ T cell repertoires, we performed single cell sequencing of CD4^+^ T cells from natalizumab-treated pwMS responding to anti-CD3 (control), ABD (irrelevant bacterial antigen), EBNA1_380-641_, ANO2_1-275_, and CRYAB antigens (Table S4A). CRYAB was added as an additional target due to the previously reported sequence homology between CRYAB and EBNA1^18^. Proliferating LiveCD3^+^CD4^+^CFSE^DIM^ cells were sorted (gating strategy in Figure S20A) and labelled with hashtag antibodies to enable identification of the original stimulation (Figure 6A and 6B). Donors who had high background proliferation in the unstimulated control condition (due to autoproliferation^54^) or no response to ANO2 or CRYAB were not taken forward, leaving four MS-Nat individuals (two *HLA-DRB1*15:01* carriers) who met these criteria. Anti-CD3 was used as a non-specific expansion control and ABD was used as a non-homologous, irrelevant antigen stimulation control.

**Figure 6.**
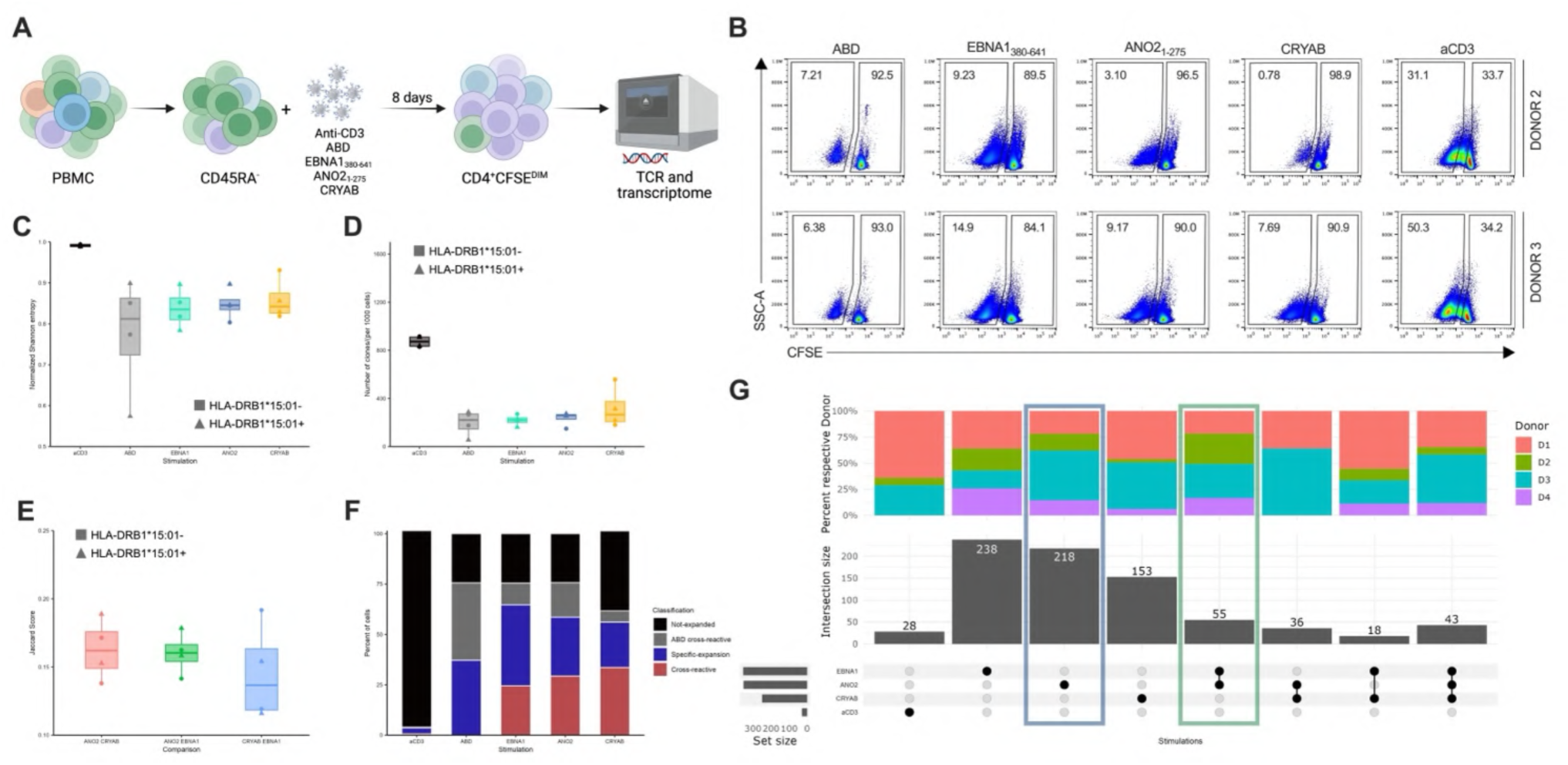
Single cell sequencing reveals a significant overlap between EBNA1 and ANO2-reactive TCR repertoires. **A.** Schematic of T cell expansions following stimulation with anti-CD3, ABD, ANO2_1-275,_ EBNA1_380-641_ and CRYAB-specific T cells from PBMC of four natalizumab-treated pwMS. Briefly, PBMC from natalizumab-treated pwMS were subjected to CD45RA negative selection, labelled with CFSE and stimulated *in vitro* for 8 days. After 8 days, cells were stained for viability, CD3 and CD4, and the responding LiveCD3^+^CD4^+^CFSE^DIM^ T cell populations were bulk sorted by FACS and processed for 10X single cell TCR sequencing. **B.** Proliferating LiveCD3^+^CD4^+^CFSE^DIM^ T cell populations after 8 days’ stimulation with ABD, EBNA1_380-641_, ANO2_1-275_ or CRYAB beads or anti-CD3 from two donors: donor 2 (top) and donor 3 (bottom). Percentage of CFSE^DIM^ and CFSE^HI^ cells of the total LiveCD3^+^CD4^+^ population are indicated in the corner of gates. **C.** Shannon entropy measure of TCR clonal diversity of repertoires from each stimulation from 10X single cell sequencing for all donors. Each symbol represents one donor. **D.** Number of unique TCR transcripts in each stimulation per 1000 CD4^+^ T cells, each symbol represents one donor. **E.** Jaccard distance score representing the overlap of expanded TCR repertoires between stimulations with MS-relevant antigens (EBNA1, ANO2 and CRYAB); a high Jaccard score represents a high degree of overlap between TCR libraries. Jaccard distance scores for each individual donor for ANO2:EBNA1, CRYAB:ANO2 and CRYAB:EBNA1. **F.** The percentage of TCR sequences in each stimulation which were unique or were also found in other repertoires, indicating their cross-reactivity. Expanded TCRs are defined as having ≥4 identical transcripts, non-expanded TCRs were classified as those with ≤3 identical transcripts. TCRs shared with ABD (irrelevant antigen, grey), TCRs only found in one stimulation (not cross-reactive, blue), cross-reactive TCRs which proliferated in response to ≥2 MS-associated antigens (EBNA1, ANO2 and/or CRYAB) and are therefore cross-reactive (red). **G.** Upset plot to indicate the intersection size of expanded TCRs in response to each stimulation in each individual donor (D1, D2, D3 and D4). Set size indicates the number of expanded TCR sequences after QC. Black dots and lines indicate stimulation identity and overlap between stimulations, sets with only one dot indicate TCRs which are unique to one stimulation and are not shared with any other condition. The blue box indicates TCRs which were only found in ANO2 stimulations, indicating T cells which are ANO2-specific but are not cross-reactive. The green box indicates expanded TCRs which are shared between EBNA1 and ANO2 stimulations only, indicating cross-reactive T cells which respond to both antigens. For the whole figure, HLA-DR*15:01^+^ donors are indicated by triangles and HLA-DR*15:01^-^ donors by squares. Albumin binding domain (ABD), T cell receptor (TCR). Created with https://BioRender.com.

After quality control, 27,524 total TCR sequences were available for analysis (Figure S20B), with details of number of cells per donor and condition shown in Figure S19C and S19D. Clonal diversity was greatest in anti-CD3 stimulated TCR repertoires and lowest in ABD stimulations, as measured by normalised Shannon entropy (Figure 6C). Expanded TCR clones were defined as detecting ≥4 identical cells indicating specific expansion in response to antigen stimulation *in vitro*, and non-expanded TCR clonotypes were classified as those with ≤3 cells. The number of unique expanded clonotypes in each condition per 1000 cells was highest in anti-CD3 stimulations, with fewer TCRs observed in antigen-specific expansions (Figure 6D), and the total number of TCR clonotypes identified per condition are shown in Figure S20E.

Promiscuous binding of TCRs to more than one antigen is a crucial feature of T cell immunity, and this fundamental aspect of T cell biology has evolved to enable TCR repertoires to respond to a vast array of potential pathogens^64^. Therefore, due to the nature of TCR degeneracy, a degree of overlap between antigen-specific TCR repertoires is expected. We therefore used stimulation with an irrelevant antigen (ABD) to distinguish cross-reactive T cells of interest from cross-reactive T cells which are not relevant for MS pathology. By removing T cells that cross-react with ABD, we can identify with a higher degree of certainty cross-reactive T cells between EBV EBNA1 and homologous MS autoantigens ANO2 and CRYAB. Expanded clonotypes in each stimulation were used as a further way of identifying T cells which were actively proliferating in response to antigen-specific stimulation.

The Jaccard similarity score is used to assess the cross-reactivity between TCR repertoires. Interestingly this measure was highest between EBNA1 and ANO2 indicating more shared TCRs, while lower scores were observed for the overlap between EBNA1 and CRYAB, indicating fewer shared TCRs despite donor variability (Figure 6E). An overlap between ANO2 and CRYAB repertoires was also detected (Figure 6E). Overall, the degree of cross-reactivity between TCR repertoires in our analysis varied between stimulations and donor. For EBNA1, around 40% of TCRs were expanded and were unique to this stimulation (not found in any other condition), but around 25% of total TCRs were expanded and were shown to be shared with ANO2 or CRYAB stimulations (Figure 6F). Likewise, around 28% of TCRs from ANO2 stimulations were unique to this condition, whereas around 26% were shared with EBNA1 and CRYAB (Figure 6F). For CRYAB, almost 30% of TCRs were unique to this stimulation, while around 20% were shared between EBNA1 and ANO2 (Figure 6F). The number of TCRs from EBNA1, ANO2 and CRYAB stimulations which were shared with ABD (the irrelevant antigen) varied between 5-15% (Figure 6F).

Next, we looked at the intersection size between EBNA1, ANO2 and CRYAB expanded TCR repertoires (after removing ABD-overlapping irrelevant cross-reactive TCRs) (Figure 6G). In total, 238 expanded TCRs were detected across all donors which were unique to the EBNA1 stimulation and were not found in other conditions, i.e. they were EBNA1-specific and not cross-reactive. Likewise, 218 expanded T cell clonotypes which were unique to ANO2 stimulations could be detected in all donors, with the highest number from donor 3 (Figure 6G, blue box). Donors 1 and 3 had the highest number of CRYAB-unique TCR clonotypes with fewer observed in donors 2 and 4 (Figure 6G). These results indicate that ANO2 and CRYAB are targeted by EBNA1-reactive T cells, but also by autoreactive T cells independently of molecular mimicry.

In total, 55 expanded TCR clonotypes were identified which were specifically shared only between EBNA1 and ANO2 stimulations (green box, Figure 6G), suggesting that these T cells are EBNA1/ANO2 cross-reactive. Likewise, 18 expanded TCR clonotypes were shared between EBNA1 and CRYAB stimulations indicating cross-reactivity in all four donors, with around 50% of these TCR clonotypes coming from donor 1. Several expanded TCRs (36 in total) were shared between ANO2 and CRYAB suggesting cross-reactivity between autoantigens, and interestingly 43 tri-cross-reactive TCR clonotypes were shared between EBNA1, ANO2 and CRYAB stimulations (Figure 6G).

Overall, these data demonstrate that the majority of EBNA1, ANO2 and CRYAB-reactive T cells are not cross-reactive, however a significant number of TCR clonotypes were shared between other stimulations indicating that they are cross-reactive – including T cells which can respond to both EBNA1 and ANO2. Therefore, our results indicate a dynamic TCR repertoire comprised of both virus and autoreactive T cells which recognise only one specific target, but comprising also cross-reactivity between two and even three antigens.

### TCRVβ CDR3b chains from expanded EBNA1/ANO2 cross-reactive TCRs are occasionally observed in independent datasets

We next investigated whether the 55 EBNA1/ANO2-expanded TCR clonotypes were more frequent in EBNA1 or ANO2 stimulations and found that the majority were equal between stimulations, but some dominated in either EBNA1 or ANO2 (Figure S20A). This suggests that some EBNA1/ANO2 cross-reactive T cells expand more in response to EBNA1 or ANO2. The number of cross-reactive versus specific TCRs for ANO2 and EBNA1 stimulations is shown in Figure S20G.

Public TCRs are defined as shared clonotypes (TCRs with identical amino acid sequences) across multiple individuals and are observed in a range of antigen-specific CD4^+^ and CD8^+^ T cell responses. By comparing repertoires between donors, we found that a small proportion of TCRs (<3%) of EBNA1, ANO2 and CRYAB stimulations were shared between multiple individuals (Figure S20H). In contrast, up to 20% of ABD-cross-reactive clones had been previously identified as public in one donor, with <3% of specific and <10% of cross-reactive TCRs being identified as public respectively (Figure S20I).

To understand whether ANO2/EBNA1 cross-reactive TCRs have distinct features compared to non-cross-reactive TCRs, we analysed their gene usage. Whilst some genes were overrepresented in the ANO2/EBNA1 cross-reactive TCRs, none reached significance (Figure S21A and B). However, a trend towards higher frequency of TRVB12-4 was observed in the cross-reactive cells, and when TRAV, TRAJ and TRBJ were analysed in expanded cells expressing TRBV12-4, we saw that there was slightly more diversity in the specific EBNA1 and ANO2 TCRs compared to cross-reactive cells (Figure S21C). Despite detecting some TCRs in our dataset which were shared between donors (public), network plot analysis of TCR repertoires revealed little amino acid similarity, meaning that cross-reactive TCRs were not conserved and did not group discretely (Figure S21D).

To understand whether the identified expanded EBNA1/ANO2 cross-reactive TCRs could be detected in other MS datasets, we overlapped our data with two recent studies^65,66^ which analysed TCR repertoires in the MS blood and cerebrospinal fluid (CSF) from pwMS. We found that 20 out of 55 expanded EBNA1/ANO2 cross-reactive clonotypes express a CDR3β sequence detected in other studies: seven overlapped with Amoriello *et al*., five overlapped with Gottlieb *et al.* and seven were found in both studies (Figure S21C). Gottlieb *et al.* bulk sequenced TCRs from EBV and autologous lymphoblastoid cell line (LCL)-expanded T cells from both CSF and blood of pwMS, and therefore shared TCRVβ CDR3b clonotypes between these datasets suggest that EBNA1/ANO2 cross-reactive T cells may also expand in response to EBV-transformed B cells^66^ (Table S6B). In contrast, Amoriello *et al.* sequenced non-expanded TCRVβ repertoires from the blood and CSF of pwMS and control subjects, and the overlap of EBNA1/ANO2 cross-reactive T cells with CSF repertoires of pwMS may suggest that these can enter the CNS (Table S6C). However, as these datasets did not analyse single cells, we cannot establish if the overlapping TCR clonotypes also have the same TCRVɑ chain and therefore no conclusions can be drawn regarding whether these cells have the same antigen specificity.

### EBNA1-, ANO2-specific and cross-reactive T cells have distinct cytotoxic and activated phenotypes

To understand if the identified antigen-specific and cross-reactive T cells (Figure 6A) had distinctive phenotypes, we also analysed the transcriptomes of responding CD4^+^ T cells in parallel with their TCRs. Clustering analysis of 27,076 cells revealed seven distinct CD4^+^ T cell subtypes which we describe based on their gene expression profiles as *activated, chronic cytotoxic, cytotoxic, inflammatory, metabolic, non-activated* and *proliferative* (Figure 7A). Number of unique molecular identifiers (UMIs) (Figure S22A), mitochondrial reads (Figure S22B), and the number of genes (Figure S22C) were all similar between clusters, and the number of cells in clusters by stimulation, classification and donor are shown in Figures S22D-F.

**Figure 7.**
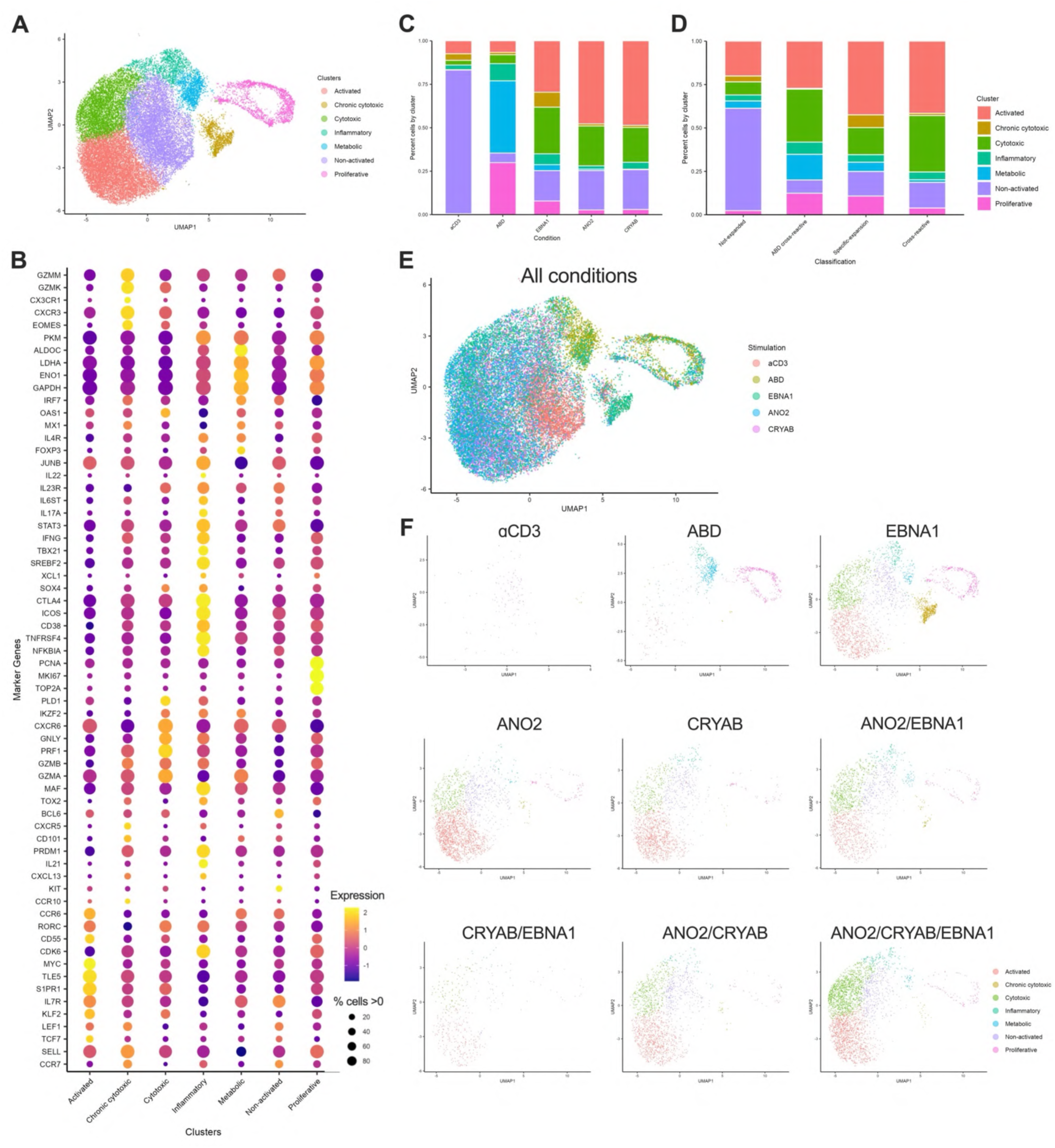
ANO2-specific and EBNA1/ANO2 cross-reactive CD4+ T cells have activated and cytotoxic phenotypes. Transcriptomes of single cells from the TCR analysis in Figure 6 (all donors). **A.** Global clustering of the transcriptomes of 27,524 T cells responding to different stimulation conditions: anti-CD3, ABD, EBNA1, ANO2 and CRYAB. T cells separated into 7 clusters: activated, chronic cytotoxic, cytotoxic, inflammatory, metabolic, non-activated and proliferative. UMAP dimensionality reduction. **B.** Canonical marker genes for cluster definitions in **A**, with relative expression (colour scale) and percent of cells expressing each marker (dot size). **C.** Cluster frequencies in relation to original stimulation conditions: anti-CD3, ABD, EBNA1, ANO2 and CRYAB (percent of total cells in each given condition). **D.** Cluster frequencies in relation TCR classifications: non-expanded, ABD cross-reactive, specifically expanded or cross-reactive (percent of total cells in each given classification). **E.** UMAP dimensionality reduction of T cells with original stimulation highlighted: anti-CD3, ABD, EBNA1, ANO2 and CRYAB. **F.** Individual UMAPs displaying T cells which are specifically expanded (only found in one condition) or cross-reactive between conditions, colour coded for the cluster they fall into. Specifically expanded T cells: anti-CD3 (top left), ABD (top middle), EBNA1 (top right), ANO2 (middle left) and CRYAB (middle). Cross-reactive expanded T cells: ANO2/EBNA1 (middle right), CRYAB/EBNA1 (bottom left), ANO2/CRYAB (middle bottom), and ANO2/CRYAB/EBNA1 (bottom right). Albumin binding domain (ABD), T cell receptor (TCR).

When looking at the top expressed genes in each cluster, cells in the *activated* cluster expressed high levels of *S1PR1*, *CCR6*, *JUNB*, *CD55*, *TCF7* and *RORC* (RORγt), whereas cells in the *chronic cytotoxic* cluster were dominated by expression of granzyme K (*GZMK*), granzyme M (*GZMM*), *CX3CR1*, *CXCR3* and *Eomes* (Figure 7B). Also of note was the *cytotoxic cluster* with high expression of CXCR6, granzyme B (GZMB), granzyme A (*GZMA*), perforin 1 (*PRF1*) and granulysin (*GNLY*) which are hallmarks of T cells with killing capacity (Figure 7B). Cells in the *inflammatory* cluster expressed high levels of *IFNγ*, *TBX21* (T-bet), *CTLA4*, *ICOS*, *CD38*, *TNFRSF4*, *IL-21*, and *PRDM1* (Blimp-1) (Figure 7B). Overall, a wide range of phenotypes was observed across responding CD4^+^ T cells in response to stimuli, and full details of top genes can be found in Figure 7B.

The original stimulus was highly linked to phenotype, with the most notable observation was the different transcriptional profiles between anti-CD3 and ABD stimulations compared with EBNA1, ANO2 and CRYAB, as shown by the unequal distribution of phenotypes across stimulations (Figure 7C) and TCR reactivity classifications (Figure 7D). Most anti-CD3-stimulated cells were in the *non-activated* cluster, reflective of cells which had not continued to actively proliferate *in vitro*. In contrast, most ABD-stimulated cells belonged to the *metabolic* and *proliferative* clusters, likely reflecting cells which were actively clonally expanding in response to this antigen. In comparison, T cells responding to the viral antigen EBNA1 showed the most diversity in transcriptional profile, with most cells fitting into the *activated, chronic cytotoxic, cytotoxic* and *non-activated* clusters (Figure 7C and F). Interestingly, the *chronic cytotoxic* cluster came almost exclusively from the EBNA1-stimulated condition (Figure 7C and F). The profiles of T cells responding to ANO2 and CRYAB were similar to that of EBNA1, with most cells in the *activated* and *cytotoxic* clusters suggesting a pathogenic phenotype *in vivo*.

We next compared the phenotypes of cells classified as not expanded (<4 TCR copies and not highly proliferating), ABD cross-reactive (shared with the irrelevant antigen stimulation ABD), specific (responding to one MS-relevant antigen only) and cross-reactive (cells shared between any combination of MS-relevant antigen stimulations: EBNA1, ANO2 and CRYAB). Non-expanded cells were dominated by the non-proliferating phenotype, whereas ABD cross-reactive cells had a highly varied and *proliferative* profile (Figure 7E). Specific expansion (not cross-reactive) cells were diverse in phenotype (Figure 7E), with absolute numbers and percentage of non-cross-reactive specific cells shown in Figure S22G and S22H respectively. Interestingly, cross-reactive T cells were similar in phenotype and were predominantly found in *activated* and *cytotoxic* clusters (Figure 7E), with EBNA1/ANO2 cross-reactive T cell population also being made up of cells from every cluster (Figure S22I and J). When looking at the cell cycle stage of cells, the *proliferative* cluster cells were demonstrated to be in either G2M or S phase, whereas most cells in all other clusters were in G1 phase (Figure S22K). No enrichment was observed for *HLA-DRB1*15:01* carriers and non-carriers across clusters (Figure S22J).

We also performed analysis of the hallmark pathways of clusters, demonstrating enrichment of the IFNγ response pathway in the *chronic cytotoxic* cluster which was overrepresented in EBNA1-specific T cells. As expected, many inflammatory pathways were upregulated in the *inflammatory* cluster, and other strong hits such as for proliferation and metabolism were upregulated in the *metabolic* and *proliferative* clusters (Figure S22J).

Together, these findings suggest that autoantigen-specific, and EBNA1/autoantigen cross-reactive T cells, have transcriptomes which are characterised by markers of activation and cytotoxic capabilities which are also linked to their antigen-specificity. Our results therefore support a role for these ANO2-specific and EBNA1/ANO2 cross-reactive CD4^+^ T cells in the immune pathogenesis of MS.

## Discussion

Here, we demonstrate elevated T cell responses to ANO2_79-168_ in both untreated and natalizumab-treated pwMS compared to healthy controls and individuals with other neurological disease, using two independent cohorts. Findings further show that, as well as being the target of antibody responses in MS^15^, ANO2 is also a T cell target in pwMS and that this response is highly associated with disease, as shown by the high AUC values in ROC analysis. Notably, the proportion of pwMS displaying ANO2-reactive T cells is much higher (approximately 57% of untreated pwMS) than the previously reported proportion of pwMS displaying anti-ANO2 antibodies (approximately 15%)^15^, further reinforcing the potential relevance of ANO2 cellular immunity in MS.

Autoreactive TCRs generally have lower avidity for their target than those which are specific for pathogens^67^ and a range of potential T cell fates is highly dependent on the strength of TCR:peptide:HLA interactions including clonal deletion, anergy or development of regulatory T cells. Lower avidity has complicated studies of autoreactive T cells as they cannot be easily detected with HLA tetramers or stimulated by overlapping peptide pools where optimal epitopes may not be in frame. Instead, our system of full-length antigen delivery into APCs via coupling to beads^40,42^ allows us to investigate T cell responses to natively processed and presented proteins with low background, especially where immunodominant epitopes are unknown or may be restricted by different HLA molecules. This method has allowed us for the first time to accurately estimate T cell responses to EBNA1_380-641_ in the context of MS – and what proportion of this repertoire can respond to two homologous MS autoantigens ANO2^15^ and CRYAB^18^ – with significant implications for the relevance of these responses to MS.

Further investigation of the ANO2-specific T cell compartment by flow cytometry confirmed FluoroSpot results and demonstrated that these cells are more likely to be CD4^+^ T cells and to produce IFNγ in natalizumab-treated pwMS, suggesting that they have a pro-inflammatory T_H_1 phenotype and therefore pathogenic potential. In addition, we observed reactivity to other regions of ANO2 with no amino acid sequence similarity to EBNA1^68,69^. These data indicate that, whilst ANO2-specific T cell responses may have been initially primed during exposure to homologous sequences in EBNA1, ANO2 T cell reactivity likely spreads to other epitopes outside the EBNA1-homologous region, although we cannot know in which order responses to multiple epitopes within ANO2 developed in humans. This could occur when naïve T cells encounter APCs which have phagocytosed ANO2 itself, myelin or other cellular debris and are presenting an array of self-peptides following CNS injury, as has been previously demonstrated in EAE^56,70,71^. This observation is also interesting given that ANO2-reactive T cells are maintained in the peripheral blood of pwMS despite long-term natalizumab therapy preventing them from entering the CNS, potentially via exposure to antigen in deep cervical lymph nodes or by sustained exposure to peripheral antigens – although maintenance of memory T cell responses does not require the presence of or repetitive stimulation by cognate antigen.

As there are cross-reactive EBNA1/ANO2-specific antibodies, there should be B cells which have surface expression of such immunoglobulins. Therefore, these B cells would lead to efficient capture and internalisation of both full-length EBNA1 and ANO2, followed by antigen presentation of a variety of epitopes on the B cells’ surface, which in turn could lead to intra-molecular epitope spreading and the multitude of reactivities against ANO2 observed in this study. Increased efficiency of T cell priming by B cells specific for the same cognate antigen has previously been demonstrated^72^. A recent study by Sattarnezhad *et al.*^17^ demonstrated that around 80% of pwMS have antibodies to at least one EBNA1 molecular mimic – ANO2, CRYAB or GlialCAM – indicating that these individuals would have B cells capable of such triggering. Interestingly, recent reports which identified autoantibodies to another chaperone protein the membrane protein modulator of VRAC current 1 (MLC1) noted that these responses may have developed as a result of epitope spreading from GlialCAM^73^. The remarkable effectiveness of anti-CD20 therapies in MS could be in part due to the depletion of these molecular mimicry B cells which propagate highly efficient autoreactive T cell priming in this context.

We previously demonstrated molecular mimicry between EBNA1 and CRYAB-specific T cell responses with cross-priming of T cells to these homologous antigens upon vaccination of SJL/J mice^18^, and we now demonstrate a similar cross-reactivity in the CD4^+^ compartment between EBNA1 and ANO2 in two different mouse strains prone to CNS autoimmunity. Experiments which compared pre-vaccination with ANO2 protein or 15-mer peptides corresponding to the same region revealed that peptide-immunised animals had similar disease severity but interestingly did not develop ANO2 autoantibodies – demonstrating that EAE severity is mediated by ANO2-specific T cells independently of humoral responses. The T cell mediated effect in adoptive transfer experiments further suggests a possible direct recognition of ANO2-expressing MOLs and OPCs by inflammation-derived IFNγ-mediated MHC class II upregulation. However, ANO2 expression, as we show at least in mice, is likely not restricted to cells of the oligodendrocyte lineage, and autoimmune attack could arise through secondary uptake of cell debris from ANO2-bearing target cells and presentation to ANO2-specific T cells. Together, these observations support that ANO2-specific CD4^+^ T cell responses are driving increased disease severity in this EAE model and facilitate disease rather than cause it. This interpretation aligns with evidence that EBV infection is required but not sufficient to cause disease, and therefore ANO2-specific and other cross-reactive T cells elicited by EBV may enable encephalitogenic T cells to gain entry into the CNS. Thus, EBV infection may lower the threshold for myelin-specific T cells to cause disease by making certain regions of the CNS more susceptible to immune infiltration, consistent with what we observe in our model.

Experiments where older mice were pre-immunised with ANO2 led to a more severe disease phenotype with higher prevalence of atypical EAE, characterised by descending paralysis, incoordination, balance problems and immune activation and infiltration in areas other than the spinal cord. This is interesting given that adult animals have been previously shown to have decreased CNS resilience^57^, and more closely reflects the median equivalent age (20-40 years) at which humans first develop MS symptoms. Development of atypical EAE in SJL/J mice has been previously demonstrated following adoptive transfer of T_H_17 cells and our study replicates this effect with direct immunisation^57^. Our results suggest that by using older animals and coupling the pre-immunisation (or adoptive transfer of T cells) against novel autoantigens to standard immunisation against myelin antigens, might reveal features of disease which are not readily achieved by immunisation against these novel antigens alone, for reasons we do not yet quite understand.

From the mechanistic perspective, our findings demonstrate that ANO2 RNA and protein are predominantly expressed in the olivary nucleus, septum, and cerebellum, with substantial co-localisation with CD31-positive vascular structures. Among these regions, the cerebellum stands out due to its marked plasma extravasation and pronounced accumulation of GFP-labelled CD4⁺ T cells following adoptive transfer. This is particularly relevant given the cerebellum’s critical role in motor coordination and balance; functions that are frequently impaired in mouse models that present with atypical EAE features. Interestingly, despite strong sharing of ANO2 expression patterns in these three brain regions, T cell aggregation is observed most prominently in the cerebellum. The reason for this regional preference remains unclear but may have important implications for disease presentation. Although we have not investigated this directly, we hypothesise that cytotoxic CD4⁺ T cells may directly target the BBB in this region, increasing its permeability and promoting local parenchymal inflammation. This targeted disruption of cerebellar integrity could account for the characteristic motor deficits seen in atypical EAE, underscoring the importance of regional vascular and immune interactions in disease development.

To further explore the role of EBNA1 and ANO2 cross-reactive T cell responses, we present the first evidence that T cells from *HLA-DRB1*15:01* carriers can target both ANO2 and EBNA1, demonstrated via both T cell clones and single cell TCR sequencing. T cell responses to EBNA1 have previously been demonstrated to be elevated and to have broader epitope specificity in MS^32^. Using EBNA1-stimulated polyclonal T cell lines from two *HLA-DRB1*15:01* carriers, we screened for reactivity to overlapping 20-mer peptide libraries presented by either DR2a^+^ (*HLA-DRB5*01:01*) or DR2b^+^ (*HLA-DRB1*15:01*) and demonstrated reactivity to two separate regions of ANO2 which differed between the donors. These results suggest that there may be further EBNA1 T cell cross-reactivity in other regions of ANO2, in addition to the aa140-150 region identified by antibody cross-reactivity^15^. These findings are corroborated by our priming experiments in the SJL/J mouse strain, showing that while animals can be primed against the aa140-150 region, immunodominant responses that include robust cross-reactivity may lie outside regions of strict sequence homology. In humans, both *HLA-DRB1*15:01* and *HLA-DRB5*01:01* are involved in autoproliferative mechanisms in MS^54^, and it is therefore likely that both genes contribute to MS disease pathology by presenting complementary peptide repertoires and by priming CD4^+^ T cells that are capable of recognising multiple antigens (including ANO2 and EBNA1) on either DR2 variant.

We performed single cell TCR sequencing of CD4^+^ cells which proliferated in response to EBNA1, CRYAB and ANO2 to determine the extent of overlap between these antigen-specific repertoires. Analysis of pwMS TCR repertoires after antigen stimulation revealed that up to 17% of EBNA1-specific TCRs were shared with the ANO2-specific compartment and a lower overlap with CRYAB. This was observed in all donors irrespective of *HLA-DRB1*15:01* carriage suggesting that cross-reactive EBNA1 CD4^+^ T cells may be a general feature of MS. This was further supported by our direct *ex vivo* analysis by FluoroSpot which indeed revealed drastically increased numbers of ANO2-reactive T cells in pwMS. Nevertheless, the extent of shared TCR sequences between EBNA1 and ANO2 is striking and, alongside evidence generated from T cell clones, strongly supports the notion that a proportion of T cells generated in response to EBNA1 are also able to target ANO2 and CRYAB. However, it is important to note that we also detected a much greater number of TCR clonotypes which were exclusively found in either ANO2 or CRYAB stimulations and are therefore not cross-reactive, suggesting that not all of these autoreactive T cells are cross-reactive with EBNA1.

Although detecting EBNA1/ANO2 cross-reactive CD4^+^ T cells in pwMS in multiple independent cohorts is not direct proof of their involvement in MS, their ability to target both viral and autoantigens suggests that a response originally primed by EBV infection leads to responses with the ability to target the CNS. However, we detected ANO2/EBNA1 cross-reactive TCRVβ chains in two other publicly available MS datasets suggesting that these T cells may be a shared feature of some pwMS, although single cell analysis of both TCRɑ and β chains is needed to confirm this. Interestingly, our single cell sequencing data showed that EBNA1/ANO2 cross-reactive T cells are present in individuals irrespective of *HLA-DRB1*15:01* carriage, which is relevant given that 40-50% of pwMS do not carry the allele^74^ and therefore at least part of the cross-reactive EBNA1/ANO2 T cells are not restricted to the *HLA-DRB1*15:01* risk haplotype. Detection of tri-cross-reactive TCRs in our dataset which could respond to EBNA1, ANO2 and CRYAB also raises further questions as to whether there is a pool of T cells with pathogenic potential which are capable of molecular mimicry between several disease-relevant proteins. T cell clones with multiple antigen reactivities have been detected in MS previously^63^, and further studies are needed to understand how widespread these degenerate T cells are.

Phenotypic analysis of autoreactive and cross-reactive CD4^+^ T cells demonstrated that these are predominantly activated and cytotoxic, also demonstrating expression of several genes which have been previously identified as associated with MS such as CXCR3 and CCR6, and demonstrating the capacity of these cells to migrate to the CNS and in autoproliferation^54,75,76^. In addition, these cells expressed granzymes and granulysin which contribute to local inflammation and tissue breakdown, leading to lesion formation^77^. It is likely that these molecules are involved in mechanisms of CD4^+^ T cells to target glial cells *in vitro* as we demonstrated in the present study. Despite culture of T cells for 8 days prior to single cell sequencing, their phenotype was maintained and was significantly different between stimulation conditions and likely gives a realistic picture of the phenotype they would have had *in vivo*. Therefore, these findings support a cytotoxic role for autoreactive and cross-reactive T cells in pwMS.

Limitations of our study is that we only sequenced expanded T cells from natalizumab-treated pwMS, and future studies should also investigate HC to understand whether these cells are a unique feature of MS. In addition, natalizumab treatment in general gives rise to a more activated phenotype of both T and B cells, which could potentially affect results in FluoroSpot and flow cytometry assays, potentially by preventing T cell migration to the gut and CNS leading them to accumulate in the peripheral blood^78,79^. However, our study nevertheless demonstrated high levels of circulating ANO2 T cells in untreated pwMS *ex vivo*, and comparison of TCR repertoires with two public datasets^65,66^ demonstrated almost no overlap of ANO2/EBNA1 cross-reactive TCRs in HC.

A further limitation is that our study only analysed EBNA1, ANO2 and CRYAB-specific CD4^+^ T cell repertoires in 4 individuals with MS, and future work should analyse a greater number of individuals (*HLA-DRB1*15:01* carriers and non-carriers) plus also TCR repertoires from both healthy controls and samples from other EBV-associated autoimmune diseases such as rheumatoid arthritis and systemic lupus erythematosus. This would provide further insights into antigen-specific TCR repertoire dynamics and degeneracy in health and disease.

Despite these limitations, we demonstrate that over half (around 57%) of pwMS have T cell responses to ANO2, which highlights the potential of these cells in aiding diagnosis, their targeted depletion for MS therapy or other methods to induce immune tolerance against ANO2. As ANO2-specific antibodies have previously been shown to appear 5-10years before MS disease onset and are associated with 26% higher sNFL in MS indicating more severe neuroaxonal damage^19^, ANO2 autoreactivity therefore has the potential to also serve as a biomarker for disease severity, and future studies should explore detection of both ANO2-reactive T cells and antibodies for diagnosis and prognosis.

Overall, we provide evidence that ANO2 is a target of EBNA1-specific T cell responses in MS, and that there is a significant overlap between these two antigen-specific TCR repertoires. More than that, we demonstrate that pre-immunisation of mice with ANO2 and EBNA1 leads to more severe MS-like disease, and that this effect is T cell-dependent. Adoptively transferred ANO2-specific T cells were tracked to areas where ANO2 is expressed in the brain, such as the cerebellum. In addition, ANO2 and EBNA1/ANO2 cross-reactive T cells from pwMS were demonstrated to have highly activated and cytotoxic phenotypes. These conclusions are underscored by the findings of increased sNFL levels in ANO2 IgG^+^ pwMS, suggesting that ANO2 responses contribute to greater neuroaxonal injury. Thus, our data supports the role of ANO2 as a T cell autoantigen in MS and also as a target of cross-reactive EBNA1 responses, providing direct mechanistic evidence for the role of EBV in MS.

## Acknowledgements

We thank P. Nilsson, C. Hellström, and the SciLife Lab Infrastructure Unit for Autoimmunity and Serology profiling for the serological analyses and their support. The PrESTs used in the suspension bead array were provided from The Human Protein Atlas project. We also thank the Neurology Clinic at Karolinska University Hospital and Neuroimmunology and MS Research Section (NIMS), Department of Neurology, University Hospital Zürich for study participant recruitment and blood sampling, B. Acharjee for PBMC preparation, L. Notari, T. Hessa and C. Quieroz for help with reagent preparation. We also thank Judith Kreutzmann for help with performing the iDISCO protocol. At the Centre for Molecular Medicine (CMM, Karolinska Institutet), we thank the Flow Facility for help with spectral flow cytometry experiments, Peri Noori and the Single Cell Facility for their guidance and help with the single cell library preparation, and the IT department for the storage and computing resources. The computations were enabled by resources provided by the Swedish National Infrastructure for Computing (SNIC) through the Uppsala Multidisciplinary Centre for Advanced Computational Science (UPPMAX), partially funded by the Swedish Research Council through grant agreement no. 2018-05973. The authors acknowledge support from the National Genomics Infrastructure in Stockholm funded by Science for Life Laboratory, the Knut and Alice Wallenberg Foundation, and the Swedish Research Council (through grant agreement no. 2018-05973). We also thank Fabian Sivnert for his continued support throughout the project.

## Disclosures

OT has received compensation for lectures and an advisory board from Merck and Pfizer respectively, which have no relation to the content of this study.

ON, GG and HG hold positions at NEOGAP Therapeutics AB.

KF has received compensation for lectures and advisory boards from Biogen, Roche, Novartis, and Merck. She has received research grants from UCB, Novartis and Merck.

FP has received research grants outside of this study from Denka, Janssen, Merck KGaA, Novartis, Pfizer and UCB, and fees for serving on DMC in clinical trials with Lundbeck and Roche, and preparation of expert witness statement for Novartis.

LA has received lecture honoraria from Teva, Biogen and Merck.

IK has received lecture honorarium from Merck and research grant from Neurogene INC which has no relation to the content of this study.

RM has received unrestricted grants from Biogen, Novartis, Roche, Third Rock and honoraries for advisory roles and lectures from Roche, Novartis, Biogen, Genzyme, Neuway, CellProtect, Third Rock, Teva. He is a patent holder and co-holder on patents of: daclizumab in MS (held by NIH), JCV VP1 for vaccination against PML; JCV-specific neutralizing antibodies to treat PML; antigen-specific tolerisation with peptide-coupled cells, novel autoantigens in MS, and designer neoantigens for tumour vaccination (all held by the University of Zurich). He is a co-founder of Abata Therapeutics, Watertown, MA, USA, and co-founder and employee of Cellerys AG, Schlieren, Switzerland, none of which have contributed to this study.

TO has received Lecture/advisory board honoraria from Merck, Novartis, Biogen and Sanofi, and unrestricted MS research grants from the same companies, none of which have contributed to this study.

All other authors declare no competing interests.

