## Supplementary Figures for "Anoctamin-2-specific T Cells Link Epstein-Barr Virus to Multiple Sclerosis"

##### Affiliations
[1] *Department of Clinical Neuroscience, Karolinska Institute, Centre for Molecular Medicine, Karolinska University Hospital, 171 76 Stockholm, Sweden.*
[2] *Division of Rheumatology, Department of Medicine, Solna, Karolinska Institutet, Karolinska University Hospital, Stockholm, Sweden.*
[3] *MS Center Amsterdam, Department of Molecular Cell Biology and Immunology, Amsterdam University Medical Center, Amsterdam, The Netherlands.*
[4] *Department of Neurology, Karolinska University Hospital, Stockholm, Sweden.*
[5] *Institute of Experimental Immunology, University of Zurich, Winterthurerstrasse 190, 8057 Zurich, Switzerland.*
[6] *Cellerys AG, Wagistrasse 21, 8952 Schlieren Switzerland*
[7] Division of Molecular Neurobiology, Department of Medical Biochemistry and Biophysics, Karolinska Institutet, Stockholm, Sweden.
[8] Department of Physiology and Pharmacology, Karolinska Institutet, Stockholm, Sweden
[9] *Division of Vascular Biology and Teaching Unit, Department of Medical Biochemistry and Biophysics, Karolinska Institutet, 171 65 Solna, Sweden.*
[10] *NEOGAP Therapeutics AB, Cancer Centre Karolinska, Visionsgatan 56, 171 64 Solna, Sweden*
[11] *Department of Medical Biochemistry and Microbiology, Uppsala University, Uppsala, Sweden*
[12] *SciLifeLab, Uppsala University, Uppsala, Sweden*
[13] *Centre for Neurology, Academic Specialist Centre, Stockholm, Sweden.*
[14] *Department of Protein Science, KTH–Royal Institute of Technology, 114 21 Stockholm, Sweden.*
[15] *Institute of Environmental Medicine, Karolinska Institute, 17177 Stockholm, Sweden.*

#### Supplementary Videos

**Video S1.** Typical EAE phenotype

**Video S2.** Atypical EAE phenotype

**Video S3.** Atypical EAE phenotype

**Video S4.** 3D staining and imaging of ANO2 and CD31 in the mouse cerebellum by iDISCO.

**Video S5.** 3D staining and imaging of ANO2 and CD31 in the mouse inferior olivary nucleus and septal region by iDISCO.

#### Supplementary Figures

##### Figure S1

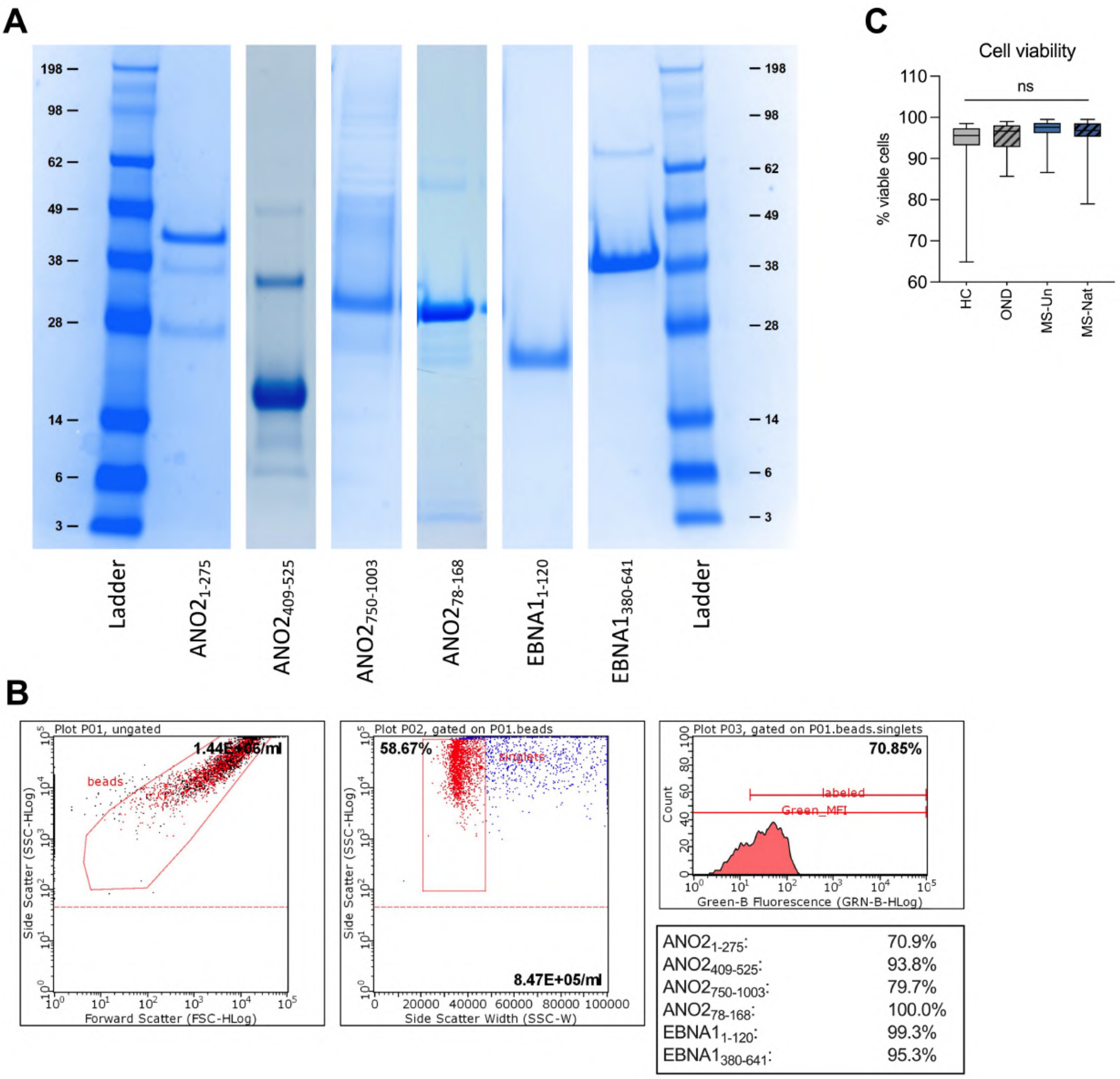


**Figure S1. Synthesis of ANO2 and EBNA1 antigen-coupled beads and PBMC viability.**

**A**. SDS-page gel of purified ANO2$_{1\text{-}275}$, ANO2$_{409\text{-}525}$, ANO2$_{750\text{-}1003}$, EBNA1$_{1\text{-}120}$ and EBNA1$_{380\text{-}640}$. Image spliced from different gels run with the same reference ladder; placing of gels in reference to the ladder is kept consistent. Numbers indicate ladder step size (kDa). Linear colour correction and brightness adjustment have been made to original image and irrelevant lanes have been cropped out.

**B**. Antigen bead coupling quality control using flow cytometry. Representative plots show gating strategy.

**C**. Viability after thawing of cryopreserved PBMC. Statistical differences were calculated using a one-way ANOVA. Healthy control (HC, n=22), other neurological disease (OND, n=20), untreated MS (MS-Un n=6) and Natalizumab-treated MS (MS-Nat n=68).

#### Figure S2

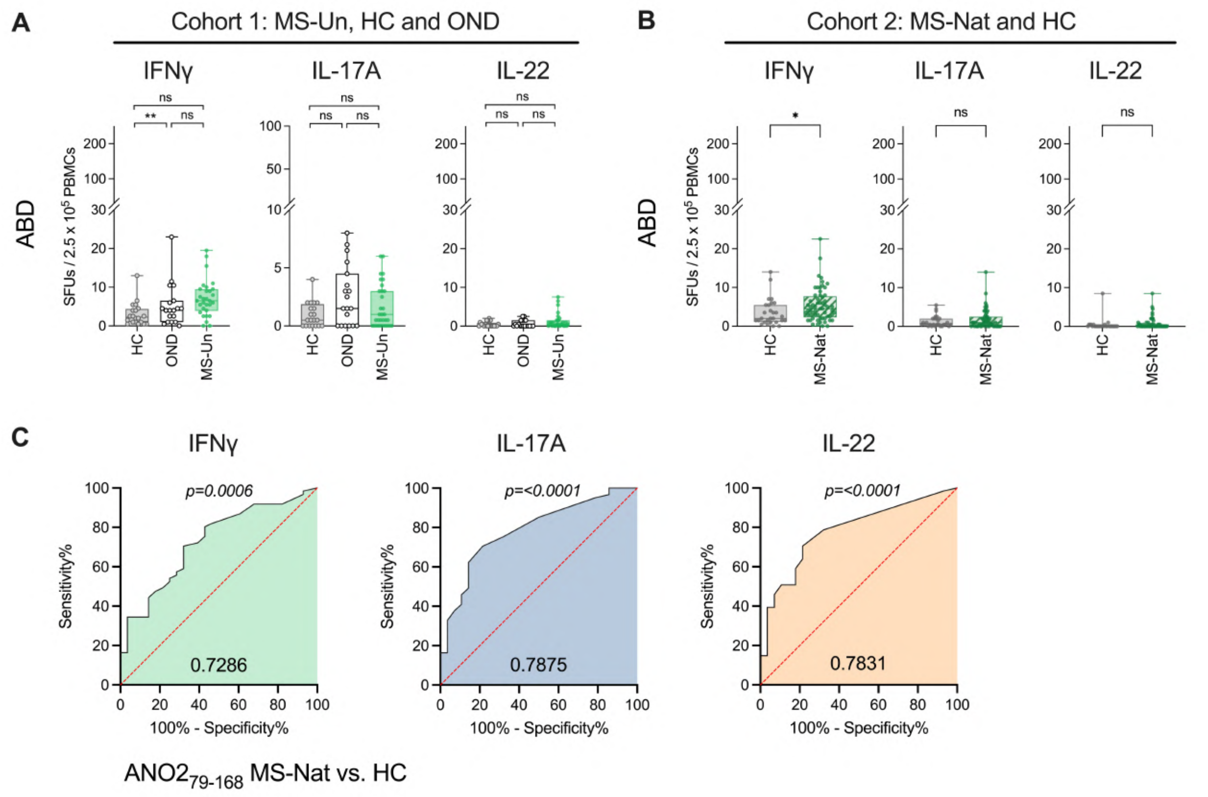


**Figure S2. Negative control bead stimulations and ANO2$_{79-168}$ ROC curves for MS-Nat donors.**

**A.** Number of IFNγ (left), IL-17A (middle) and IL-22 (right) SFU after stimulation in a FluoroSpot assay with albumin binding domain (ABD) negative control beads in cohort 1: MS-Un (n=30), HC (n=20) and OND (n=19). Boxes represent median ± IQR and statistical significance was calculated with the Kruskall-Wallis test with Dunn's multiple comparisons test with p-values indicated where significant.

**B.** Number of IFNγ (left), IL-17A (middle) and IL-22 (right) SFU after stimulation with NC beads in cohort 2: MS-Nat (n=61) and HC (n=28). Boxes represent median ± IQR, statistical significance was calculated with the Mann-Whitney test and p-values indicated where significant.

**C.** Receiver operating characteristic (ROC) based on IFNγ (left), IL-17A (middle) and IL-22 (right) responses to ANO2$_{79-168}$ beads in control (HC n=28) and MS-Nat donors (n=68). Area under curve indicated at the bottom of graphs and p-values indicated above graphs.

#### Figure S3

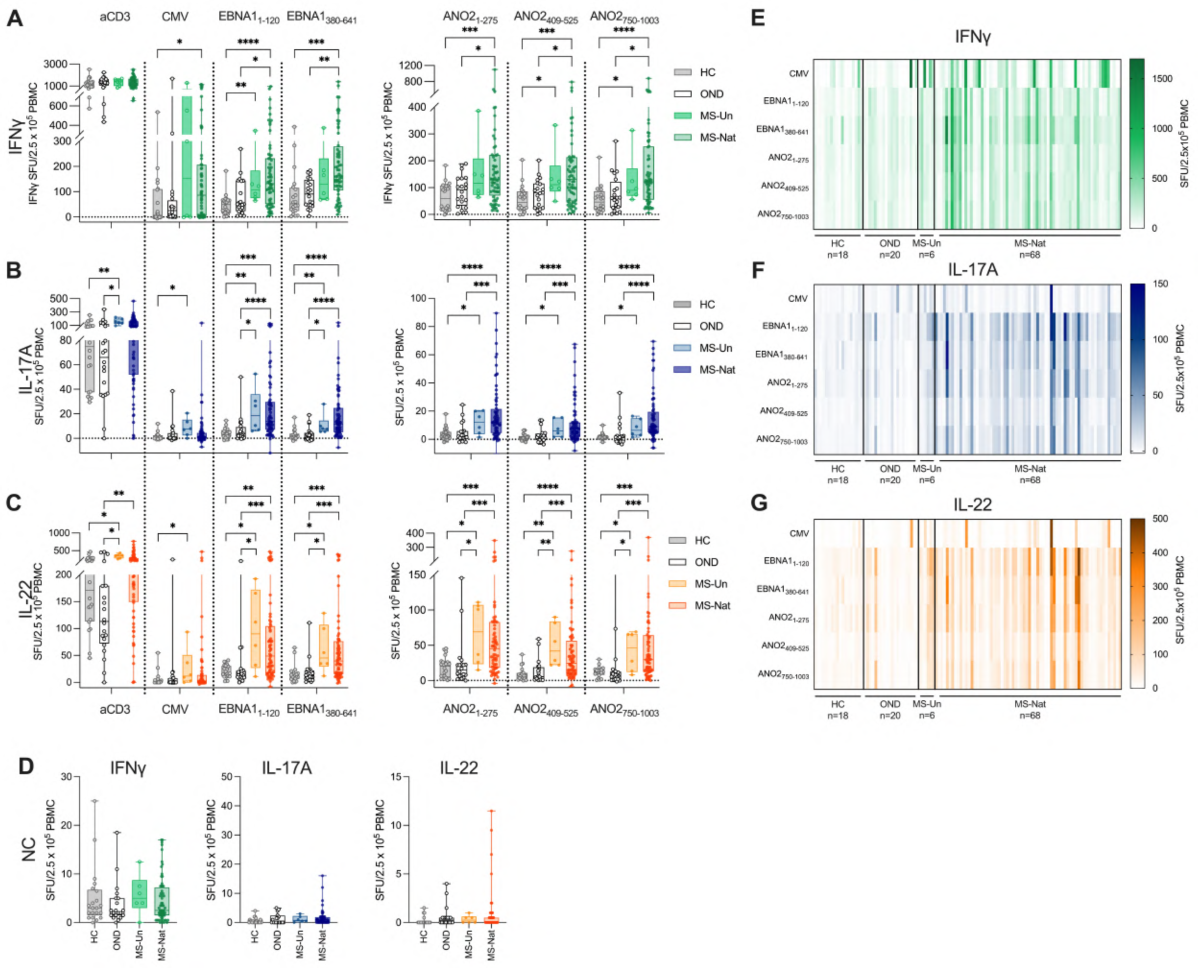


**Figure S3. ANO2 and EBNA1 are targets of T cell responses in MS.**

**A.** Number of IFNγ,

**B.** IL-17A and

**C.** IL-22 SFU after stimulation in a FluoroSpot assay with anti-CD3, CMV pp65, $EBNA1_{1\text{-}120}$, $EBNA1_{380\text{-}641}$ (left panel), $ANO2_{1\text{-}275}$, $ANO2_{409\text{-}525}$ and $ANO2_{750\text{-}1003}$ (right panel) in a cohort of healthy control (HC, n=22), other neurological disease (OND, n=20), untreated MS (MS-Un n=6) and Natalizumab-treated MS (MS-Nat n=68). Background subtracted SFU using the negative control (NC) bead stimulation.

**D.** Number of SFU after stimulation with NC beads in the same cohort as A-C. Boxes represent median ± IQR and statistical significance was calculated with the Kruskall-Wallis test with Dunn's multiple comparisons.

**E.** Heatmap of IFNγ SFU responses in individuals (data from A).

**F.** IL-17A SFU responses in individuals (data from B).

**G.** IL-22 SFU responses in individuals (data from C).

Spot-forming units (SFU). P-values indicated where significant, $*p < 0.05$; $**p < 0.01$; $***p < 0.001$; $****p < 0.0001$.

#### Figure S4

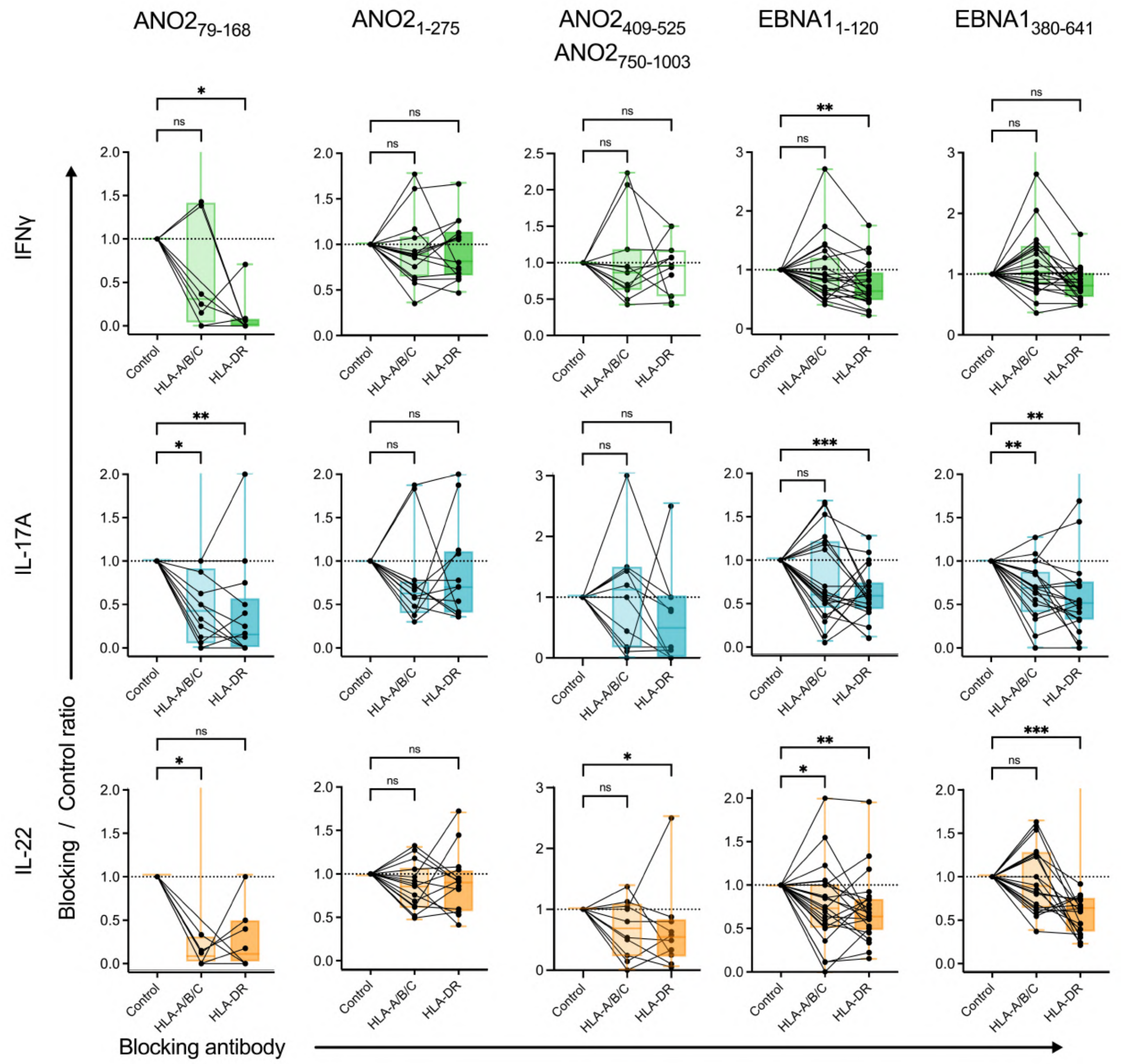


**Figure S4. HLA blocking of T cell responses by FluoroSpot.**

PBMC from Natalizumab-treated MS donors were stimulated in a FluoroSpot assay with ANO2$_{79-168}$ (n=23), ANO2$_{1-275}$ (n=15), ANO2$_{409-525}$ and ANO2$_{750-1003}$ (n=12), EBNA1$_{1-120}$ (n=23), EBNA1$_{380-641}$ (n=23) beads. Cytokine production of IFNγ (top), IL-17A (middle) and IL-22 (bottom) was measured after addition of isotype control, HLA-A/B/C and HLA-DR-blocking antibodies to wells to understand whether T cell responses were restricted by HLA class I or II. Unresponsive individuals (<6 SFU) were excluded from the analysis. Results plotted as the ratio of blocking antibodies versus isotype control. Boxes represent median and IQR, with the range in brackets. *P* values were calculated using two-tailed Wilcoxon signed-rank test.

*$p < 0.05$; **$p < 0.01$; ***$p < 0.001$.

**Figure S5**

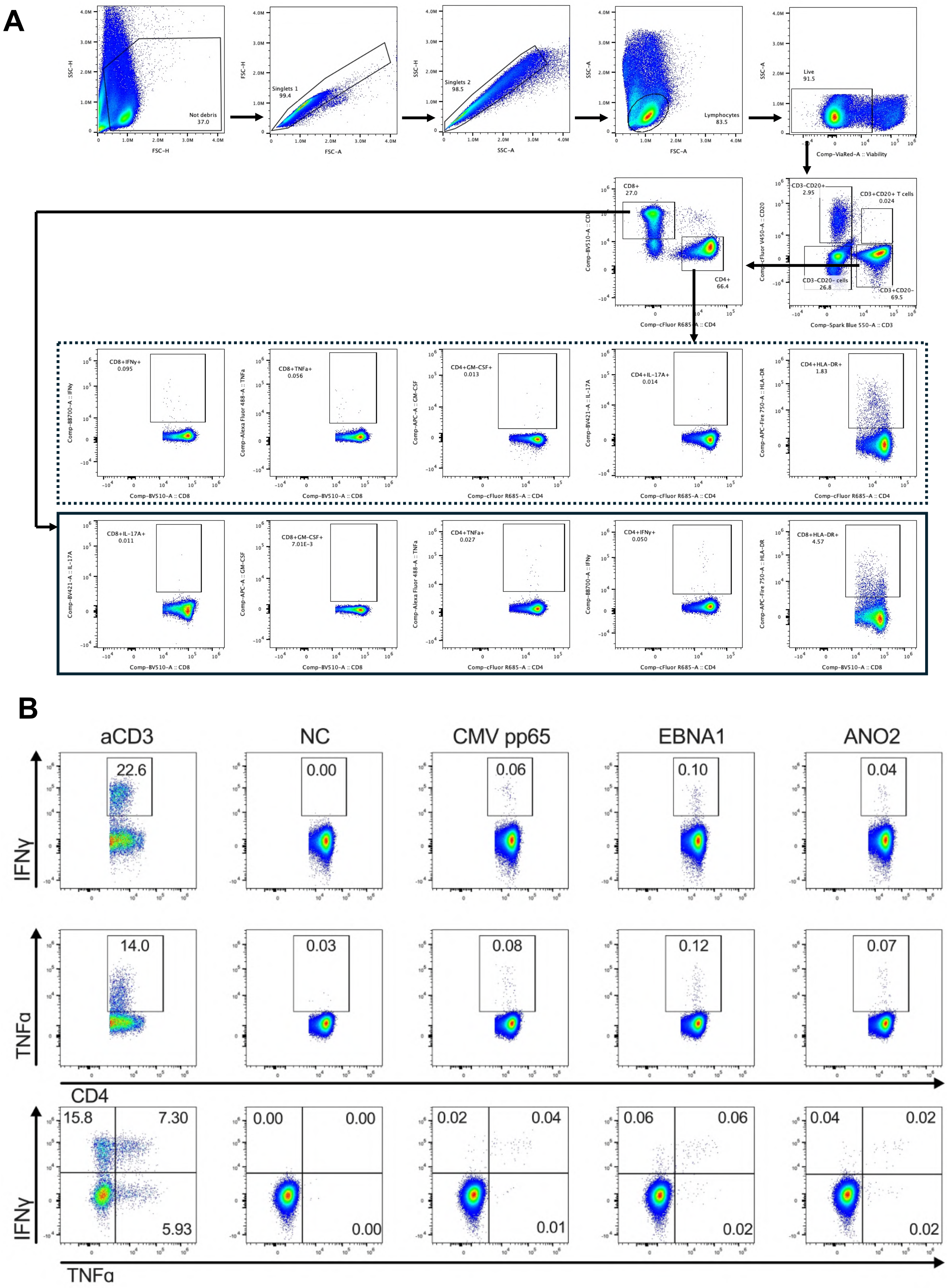
A
Not debris
37.0
Singlets 1
99.4
Singlets 2
98.5
Lymphocytes
83.5
Live
91.5
Comp-ViaRed-A :: Viability
CD3-CD20+
2.95
CD3+CD20+ T cells
0.024
CD3-CD20- cells
26.8
CD3+CD20-
69.5
Comp-Spark Blue 550-A :: CD3
CD8+
27.0
CD4+
66.4
Comp-cFluor R685-A :: CD4
CD8+IFNγ+
0.095
CD8+TNFa+
0.056
CD4+GM-CSF+
0.013
CD4+IL-17A+
0.014
CD4+HLA-DR+
1.83
CD8+IL-17A+
0.011
CD8+GM-CSF+
7.01E-3
CD4+TNFa+
0.027
CD4+IFNγ+
0.050
CD8+HLA-DR+
4.57
B
aCD3
NC
CMV pp65
EBNA1
ANO2
IFNγ
TNFα
CD4
TNFα
22.6
0.00
0.06
0.10
0.04
14.0
0.03
0.08
0.12
0.07
15.8
7.30
5.93
0.00
0.00
0.00
0.02
0.04
0.01
0.06
0.06
0.02
0.04
0.02
0.02

**Figure S5 continued**

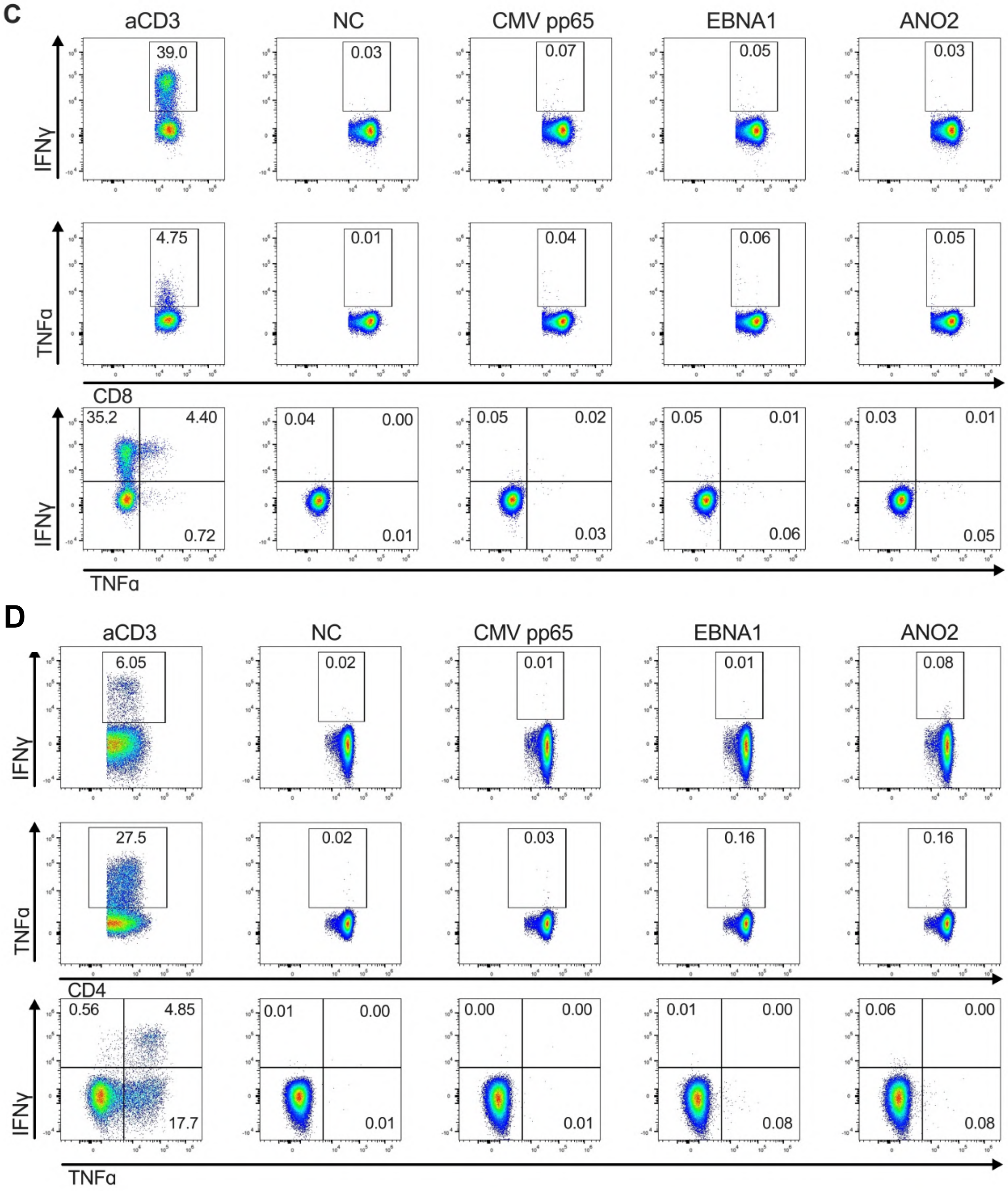

**Figure S5 continued**

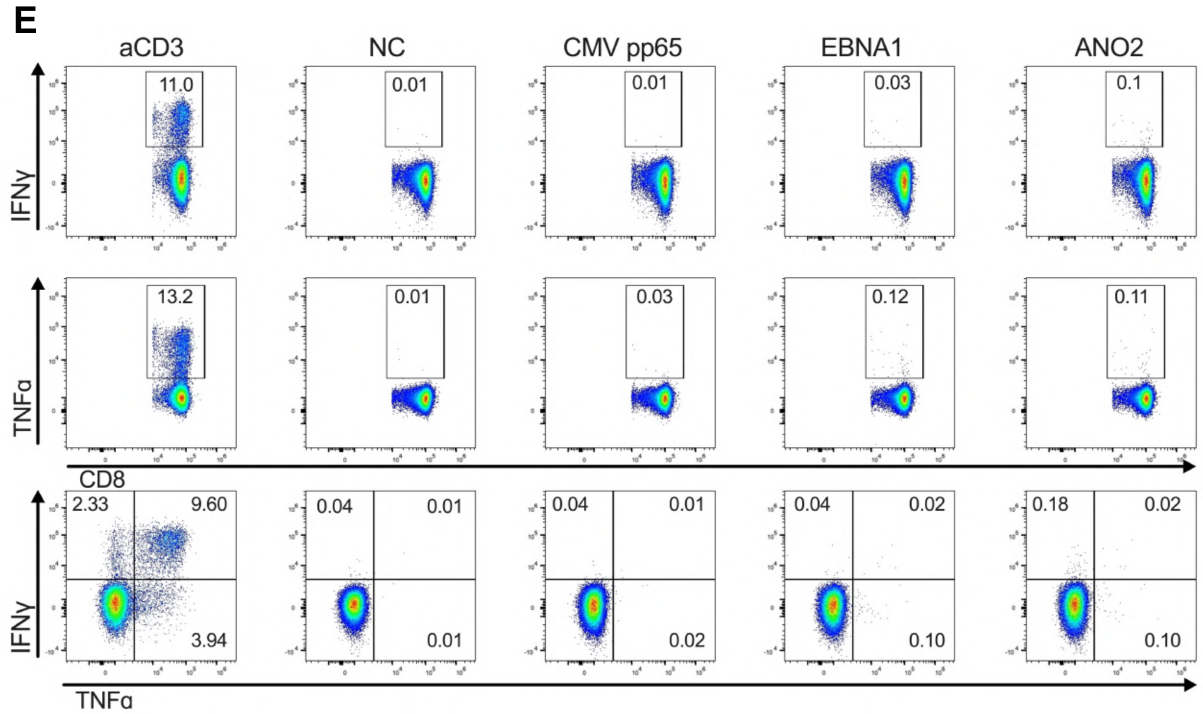


**Figure S5. Intracellular cytokine staining and spectral flow cytometry gating strategy for detection of *ex vivo* antigen-specific T cell responses in PBMC.**

**A.** Spectral flow cytometry analysis of autoreactive T cells using intracellular cytokine staining after stimulation with PMA/ionomycin, CMV pp65, EBNA1 or ANO2 beads.

**B.** Example $CD4^+$ T cell of intracellular cytokine staining from a representative MS-Nat donor (P68) after PBMC stimulation with anti-CD3, negative control (NC) beads, CMV pp65 beads, EBNA1 beads or ANO2 beads. Staining shows the $SingleLiveCD3^+CD20^-CD4^+$ population.

**C.** Example of $CD8^+$ T cell intracellular cytokine staining from a representative MS-Nat donor (P68) after PBMC stimulation with anti-CD3, negative control (NC) beads, CMV pp65 beads, EBNA1 beads or ANO2 beads. Staining shows the $SingleLiveCD3^+CD20^-CD8^+$ population.

**D.** Example $CD4^+$ T cell of intracellular cytokine staining from a representative HC donor (C36) after PBMC stimulation with anti-CD3, negative control (NC) beads, CMV pp65 beads, EBNA1 beads or ANO2 beads. Staining shows the $SingleLiveCD3^+CD20^-CD4^+$ population.

**E.** Example of $CD8^+$ T cell intracellular cytokine staining from a representative HC donor (C36) after PBMC stimulation with anti-CD3, negative control (NC) beads, CMV pp65 beads, EBNA1 beads or ANO2 beads. Staining shows the $SingleLiveCD3^+CD20^-CD8^+$ population.

### Figure S6

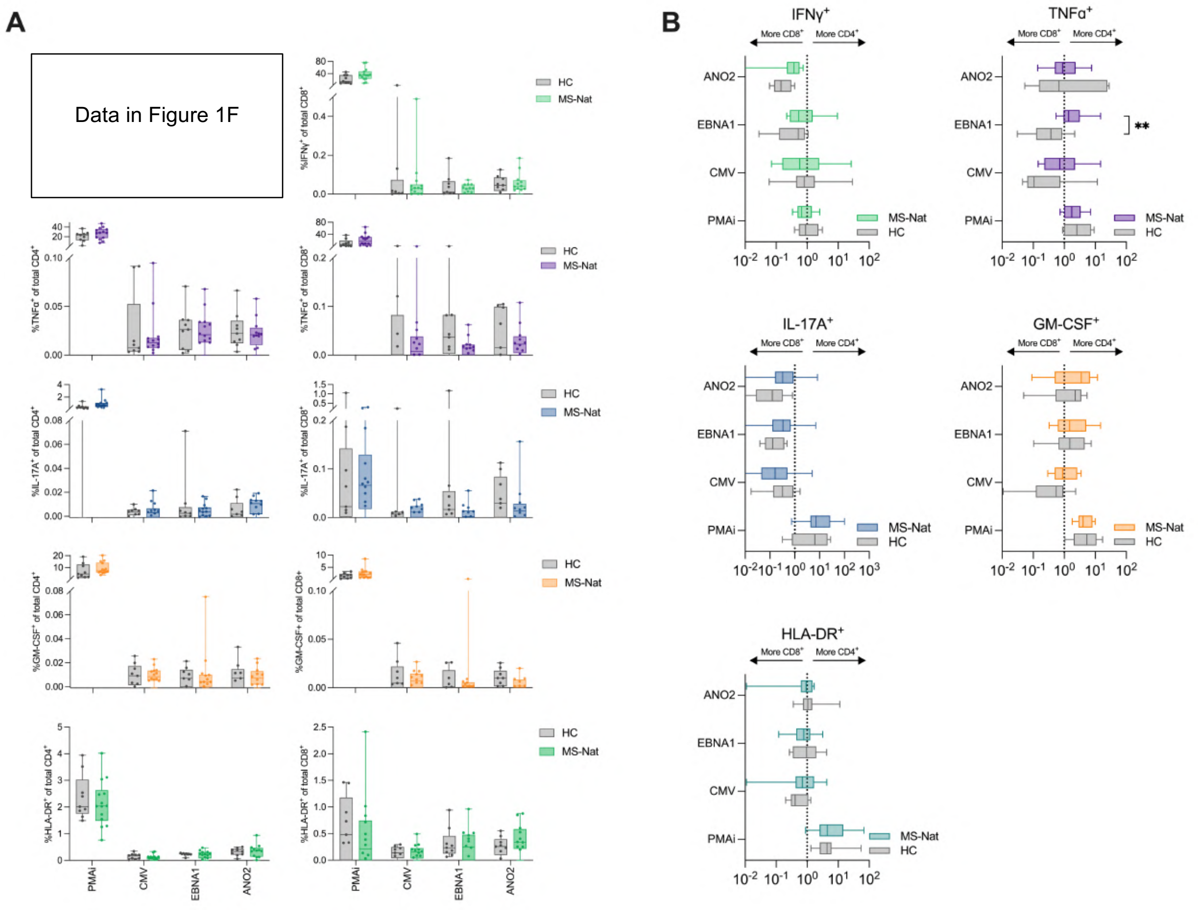


**Figure S6. *Ex vivo* T cell responses by flow cytometry.**

Spectral flow cytometry analysis of autoreactive T cells using intracellular cytokine staining after stimulation with PMA/ionomycin, CMV pp65, EBNA1 or ANO2 beads.

**A.** $CD4^+$ T cells producing TNFα, IL-17A and GM-CSF or expressing HLA-DR after stimulation with autoantigens. $CD4^+IFN\gamma^+$ responding T cells can be found in Figure 1F. $CD8^+$ T cells producing IFNγ, TNFα, IL-17A and GM-CSF or expressing HLA-DR after stimulation with autoantigens. MS-Nat (n=12) and HC (n=9) donors.

**B.** Ratio of % activation marker-expressing $CD4^+$ and % $CD8^+$ after stimulation in both MS-Nat and HC. Boxes represent median and IQR, and brackets represent 1.5 × IQR (Tukey). *P* values were calculated using two-tailed Mann-Whitney *U* test and indicated when significant.

$*P < 0.05$ and $**P < 0.01$.

#### Figure S7

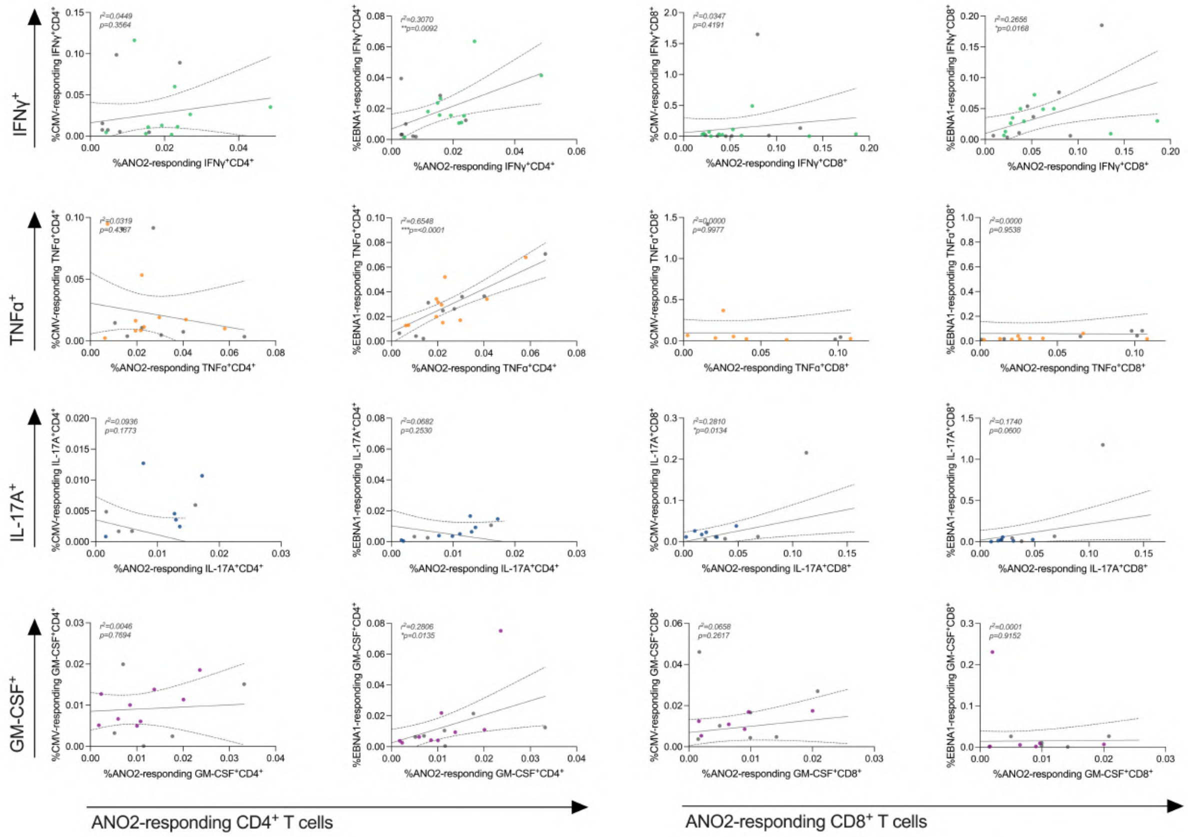


**Figure S7. Correlation of *ex vivo* T cell responses by flow cytometry.**

Spectral flow cytometry analysis of autoreactive T cells using intracellular cytokine staining after autoantigen stimulation. $CD4^+$ and $CD8^+$ T cells producing IFNγ, TNFα, IL-17A and GM-CSF after stimulation with antigen beads in MS-Nat (n=12) and HC (grey circles, n=9). Correlation between $CD4^+$ and $CD8^+$ T cell responses to CMV pp65, EBNA1 or ANO2 beads. Lines represent the best-fit nonlinear regression slope, and dashed lines represents the 95% confidence interval. Grey dots represent HC and coloured dots represent MS-Nat individuals. *P*-values were calculated using two-tailed non-parametric Spearman correlation tests and indicated where significant.

$^*P < 0.05$; $^{**}P < 0.01$; $^{***}P < 0.001$.

### Figure S8

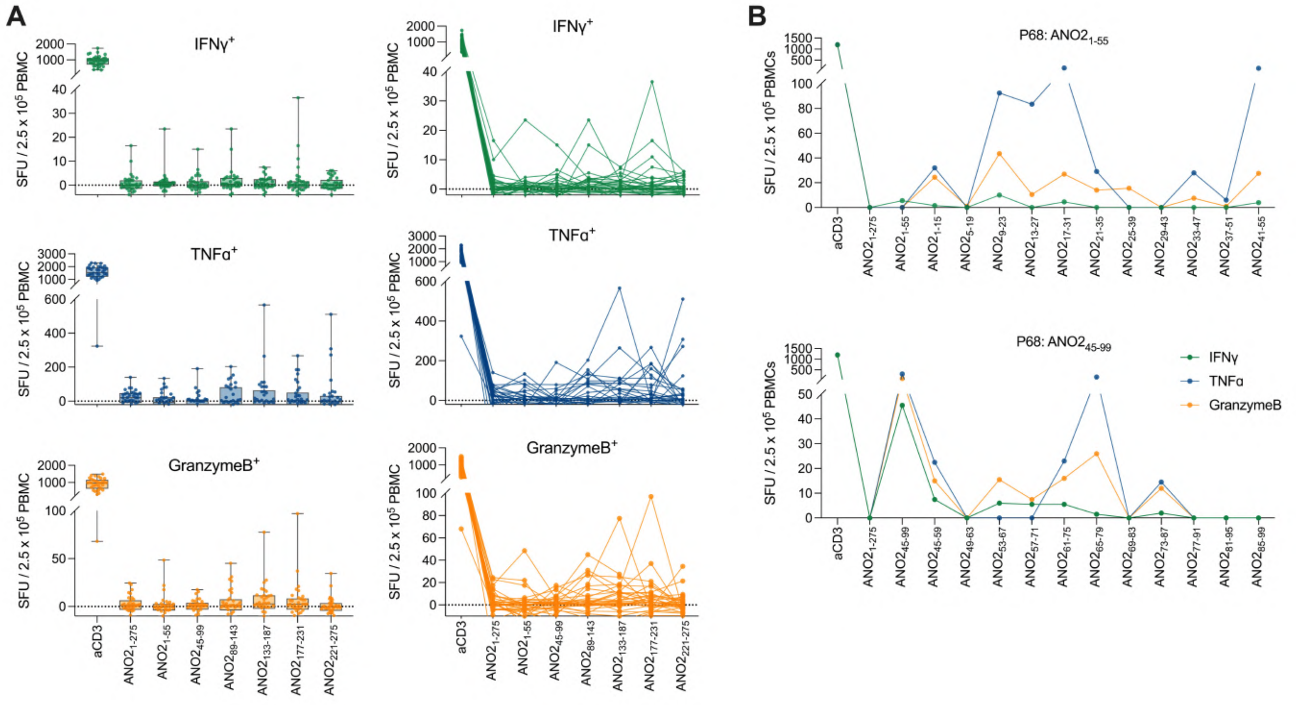


**Figure S8. Mapping of ANO2$_{1\text{-}275}$ T cell epitopes by FluoroSpot.**

**A.** A library of 15-mer peptides overlapping by 11 amino acids spanning ANO2$_{1\text{-}275}$ were used to map T cell epitopes in a IFNγ/TNFα/Granzyme B FluoroSpot assay. 6 pools of 11 peptides spanning ANO2$_{1\text{-}275}$were initially screened for reactivity in MS-Nat PBMC (n=35) and reactivity was detected to all pools including regions without homology to EBNA1. Bar charts (left column) showing overall data and lines representing each donor's responses to all pools (right column).

**B**. Epitope mapping of ANO2$_{1\text{-}55}$ (top) and ANO2$_{45\text{-}99}$ (bottom) using 15-mer peptides in an MS-Nat donor P68 by IFNγ/TNFα/Granzyme B FluoroSpot. Multiple epitopes distinct epitopes are the targets of T cell responses in this individual.

Natalizumab-treated MS (MS-Nat).

#### Figure S9

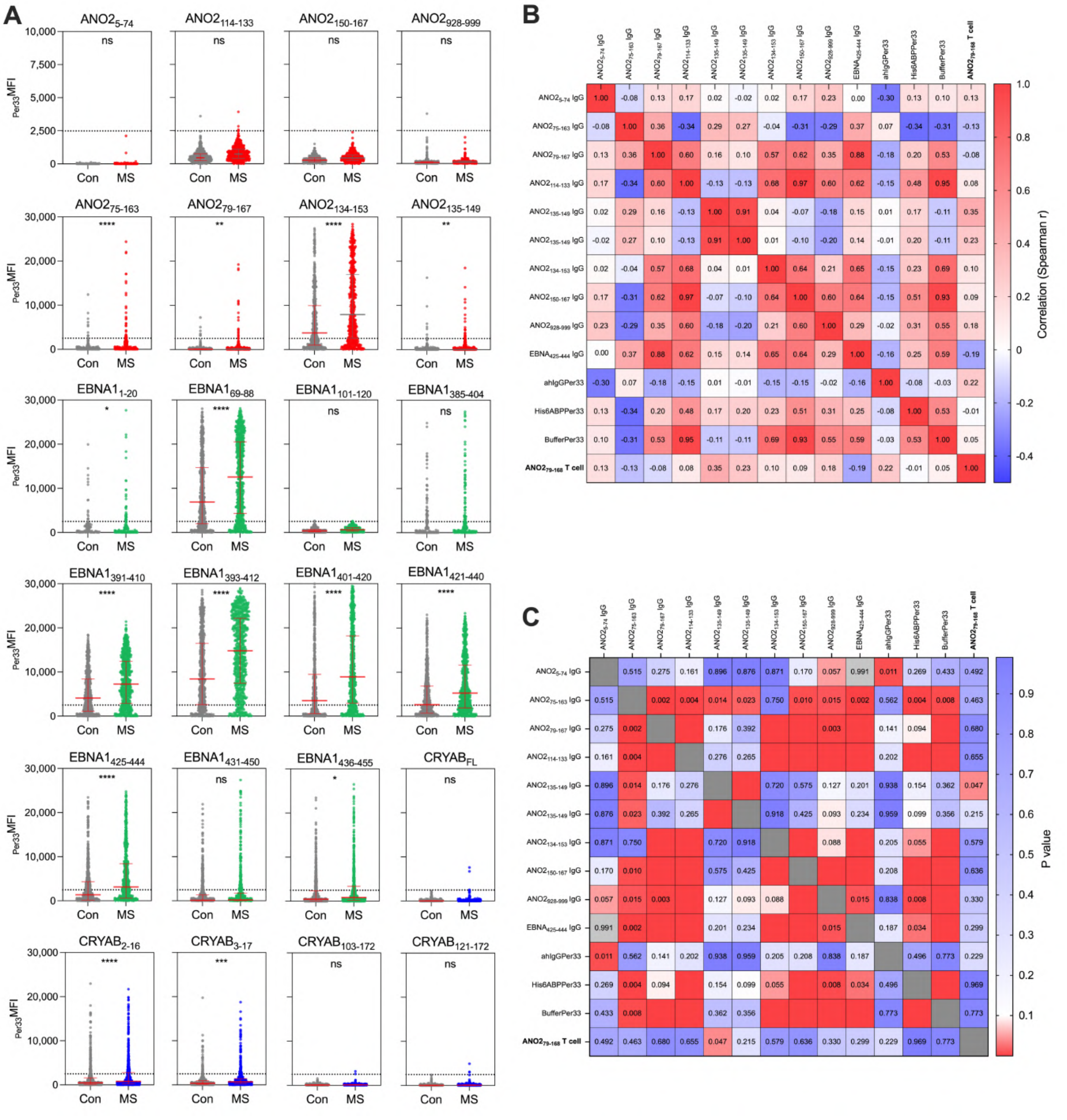


**Figure S9. Antibody responses to EBNA1, ANO2 and CRYAB and correlations with T cell responses.**

**A.** Suspension bead array measuring immunoglobulin G (IgG) against ANO2, EBNA1 and CRYAB peptides and proteins using plasma from pwMS (n =713) and controls (Con) (n =722). Background-adjusted mean fluorescent intensity [33rd percentile MFI ($_{Per33}$MFI)] values of ANO2 (red), EBNA1 (green) and CRYAB peptide and protein responses. Each dot represents one individual and staples denote the median and interquartile range (IQR). A threshold for positive response was calculated based on the cohort's responses to negative peptides (threshold at 99.9$^{th}$ percentile response $_{Per33}$MFI of 2484.85 (indicated by the horizontal dotted line). Fisher's exact statistical test was used to compare the number of positive responses between groups.

**B.** Correlation between selected ANO2$_{79-168}$ IFNγ$^{+}$ T cell responses by FluoroSpot (bold text) and selected ANO2 and EBNA1 peptide IgG responses by suspension bead array (log10 $_{Per33}$MFI) in 33 individuals (MS n=27, Con n=6). Spearman correlation coefficient (r) represented by positive correlation (red) and negative correlation (blue).

**C.** P-values of Spearman correlation coefficient (r) in **B**. Significant P-values ($P<0.05$) and are indicated by red colour, blank red squares indicate $P<0.001$.

*P<0.05; **P<0.01; ***P<0.001; ****P<0.0001 (adjusted P values).

**Figure S10**

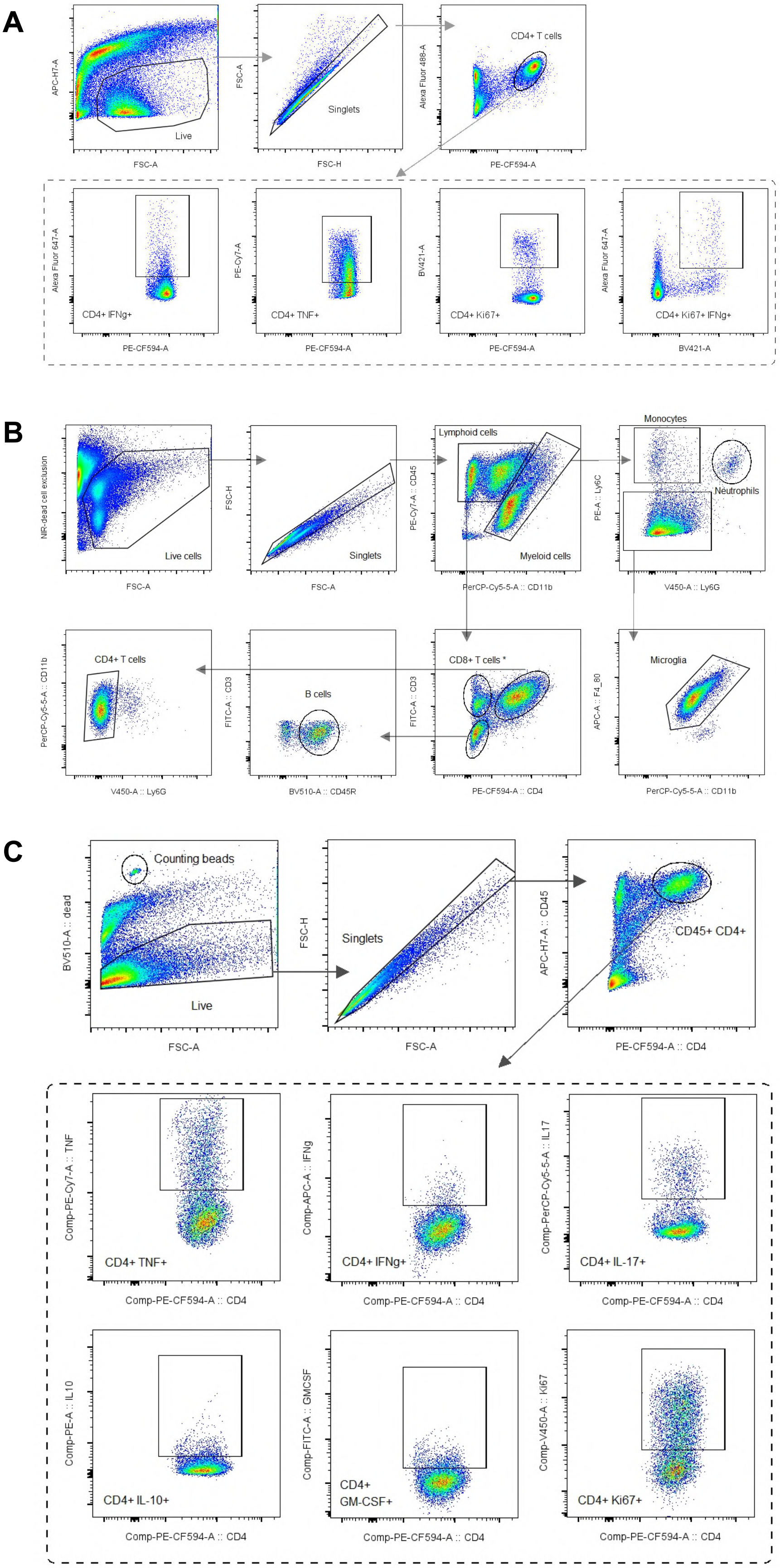
A
APC-H7-A
FSC-A
Live
FSC-A
FSC-H
Singlets
Alexa Fluor 488-A
PE-CF594-A
CD4+ T cells
Alexa Fluor 647-A
CD4+ IFNg+
PE-Cy7-A
CD4+ TNF+
BV421-A
CD4+ Ki67+
CD4+ Ki67+ IFNg+
BV421-A
B
NIR-dead cell exclusion
Live cells
FSC-H
Singlets
PE-Cy7-A :: CD45
PerCP-Cy5-5-A :: CD11b
Lymphoid cells
Myeloid cells
PE-A :: Ly6C
V450-A :: Ly6G
Monocytes
Neutrophils
PerCP-Cy5-5-A :: CD11b
CD4+ T cells
FITC-A :: CD3
BV510-A :: CD45R
B cells
CD8+ T cells *
PE-CF594-A :: CD4
APC-A :: F4_80
Microglia
C
BV510-A :: dead
Counting beads
Live
Singlets
APC-H7-A :: CD45
PE-CF594-A :: CD4
CD45+ CD4+
Comp-PE-Cy7-A :: TNF
CD4+ TNF+
Comp-PE-CF594-A :: CD4
Comp-APC-A :: IFNg
CD4+ IFNg+
Comp-PerCP-Cy5-5-A :: IL17
CD4+ IL-17+
Comp-PE-A :: IL10
CD4+ IL-10+
Comp-FITC-A :: GMCSF
CD4+
GM-CSF+
Comp-V450-A :: Ki67
CD4+ Ki67+

**Figure S10. Flow cytometry gating strategies for mouse antigen recall experiments in lymph nodes and brain populations and cytokines.**

**A.** Flow cytometry gating strategy for mouse lymph node *in vitro* antigen recall experiments.

**B.** Flow cytometry gating strategy for EAE mouse brain populations *ex vivo*.

**C.** Flow cytometry gating strategy for EAE mouse brain cytokines *ex vivo*.

#### Figure S11

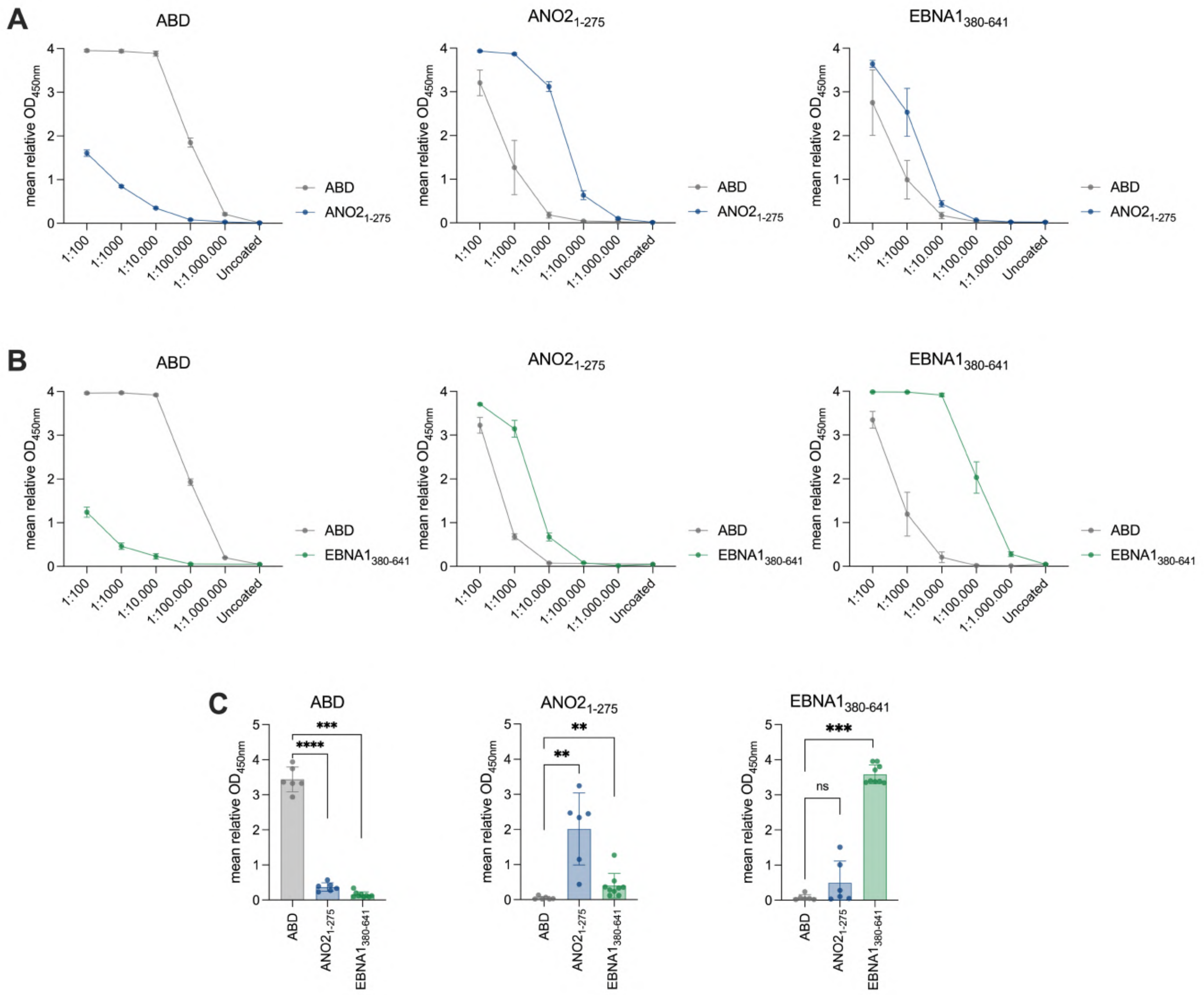


**Figure S11. Optimisation of mouse IgG ELISAs.**

Optimisation of serum dilutions for in-house ELISAs. Plates were coated with recombinant ABD, $ANO2_{1-275}$ or $EBNA1_{380-641}$ proteins and sera added in serial dilution. Sera collected at day 50 was used for ELISA optimisations.

**A.** Serially diluted sera from ABD (n=3) and $ANO2_{1-275}$ pre-immunised (n=3) mice at day 50 after EAE induction were added to plates coated with recombinant proteins: ABD (left), $ANO2_{1-275}$ (middle) and $EBNA1_{380-641}$ (right).

**B.** Serially diluted sera from ABD (n=3) and $EBNA1_{380-641}$ pre-immunised (n=3) mice at day 50 after EAE induction were added to plates coated with recombinant proteins: ABD (left), $ANO2_{1-275}$ (middle) and $EBNA1_{380-641}$ (right).

**C.** Sera from ABD (n=6), $ANO2_{1-275}$ (n=6) and $EBNA1_{380-641}$ pre-immunised (n=9) mice were optimally diluted (1:50,000) and added to ABD (left), $ANO2_{1-275}$ (middle) and $EBNA1_{380-641}$ (right)-coated plates.

Optical density (OD). *$P < 0.05$; **$P < 0.01$; ***$P < 0.001$.

#### Figure S12

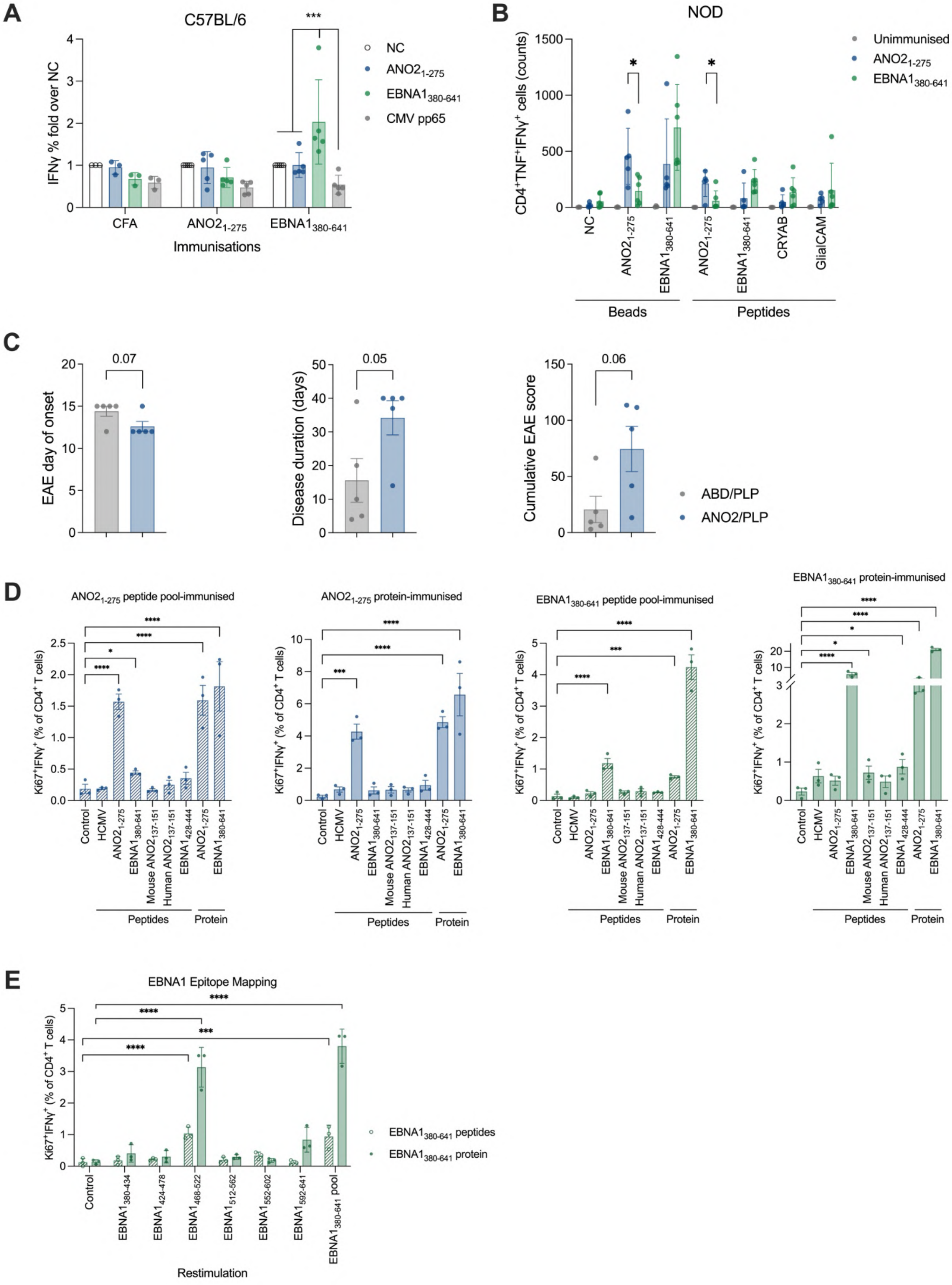
A
C57BL/6
IFNγ % fold over NC
NC
ANO2 1-275
EBNA1 380-641
CMV pp65
CFA
Immunisations
***
B
NOD
CD4+TNF+IFNγ+ cells (counts)
Unimmunised
NC
CRYAB
GlialCAM
Beads
Peptides
C
EAE day of onset
Disease duration (days)
Cumulative EAE score
0.07
0.05
0.06
ABD/PLP
ANO2/PLP
D
ANO2 1-275 peptide pool-immunised
ANO2 1-275 protein-immunised
EBNA1 380-641 peptide pool-immunised
EBNA1 380-641 protein-immunised
Ki67+IFNγ+ (% of CD4+ T cells)
Control
HCMV
Mouse ANO2 137-151
Human ANO2 137-151
EBNA1 428-444
Peptides
Protein
E
EBNA1 Epitope Mapping
EBNA1 380-434
EBNA1 424-478
EBNA1 468-522
EBNA1 512-562
EBNA1 552-602
EBNA1 592-641
EBNA1 380-641 pool
EBNA1 380-641 peptides
EBNA1 380-641 protein
Restimulation

**Figure S12. Priming of ANO2 and EBNA1 T cell responses in C57BL/6, NOD and SJL/J mouse strains.**

**A.** Draining lymph node cells from adjuvant only (CFA, n=3), $ANO2_{1-275}$ (n=5) or $EBNA1_{380-641}$-immunised (n=5) C57BL/6 mice were rechallenged with bead bound antigen from MS-associated autoantigens and the responding T cells analysed by flow cytometry and intracellular cytokine staining (ICS). Data presented as the % fold change $IFN\gamma^{+}$ cells over the negative control (NC) stimulation. Mann-Whitney test.

**B.** Draining lymph node cells from unimmunised (n=3), $ANO2_{1-275}$ (n=5) or $EBNA1_{380-641}$-immunised (n=6) non-obese diabetic (NOD) mice were rechallenged with bead-bound antigen or peptides from MS-associated antigens and responding T cells analysed by flow cytometry and ICS. Mann-Whitney test.

**C.** Data from Figure 2D. Clinical phenotype of mice sacrificed at day 51 post-immunisation represented by day of EAE onset (left), EAE disease duration (centre), and cumulative EAE score (right) in ABD/PLP (n=5) and $ANO2_{1-275}$/PLP-immunised mice (n=5). Data was non-normally distributed and was analysed using the Mann-Whitney test.

**D.** Extended data from Figure 2K. Splenocytes from SJL/J mice which were immunised with $ANO2_{1-275}$ peptide pool (n=3), $ANO2_{1-275}$ protein (n=3), $EBNA1_{380-641}$ peptide pool (n=3) and $EBNA1_{380-641}$ protein (n=3) were rechallenged *in vitro* with each of the antigens used for original immunisations to test for antigen-specific priming of T cell responses. Responses were determined by ICS and flow cytometry and indicated by Ki67 and IFNγ expression. One way ANOVA with Dunnet's multiple comparisons test.

Data from HCMV $IE1_{162-173}$ peptide (n=3), human $ANO2_{137-151}$ peptide (n=3) and $EBNA1_{428-444}$ peptide (n=3)-immunised animals in Figure 2K.

**E.** Splenocytes from SJL/J mice which were immunised with $EBNA1_{380-641}$ peptide pool (n=3) or $EBNA1_{380-641}$ protein (n=3) were rechallenged *in vitro* with overlapping peptides covering different regions of EBNA1 (15-mers overlapping by 11aa, 10 or 11 peptides per pool). One way ANOVA with the Holm-Sidak multiple comparisons test.

Albumin binding domain (ABD), Anoctamin-2 (ANO2), complete Freund's adjuvant (CFA), human cytomegalovirus (HCMV), immediate early 1 (IE1), intracellular cytokine staining (ICS). **P* < 0.05; ***P* < 0.01; ****P* < 0.001, *****P* < 0.0001.

### Figure S13

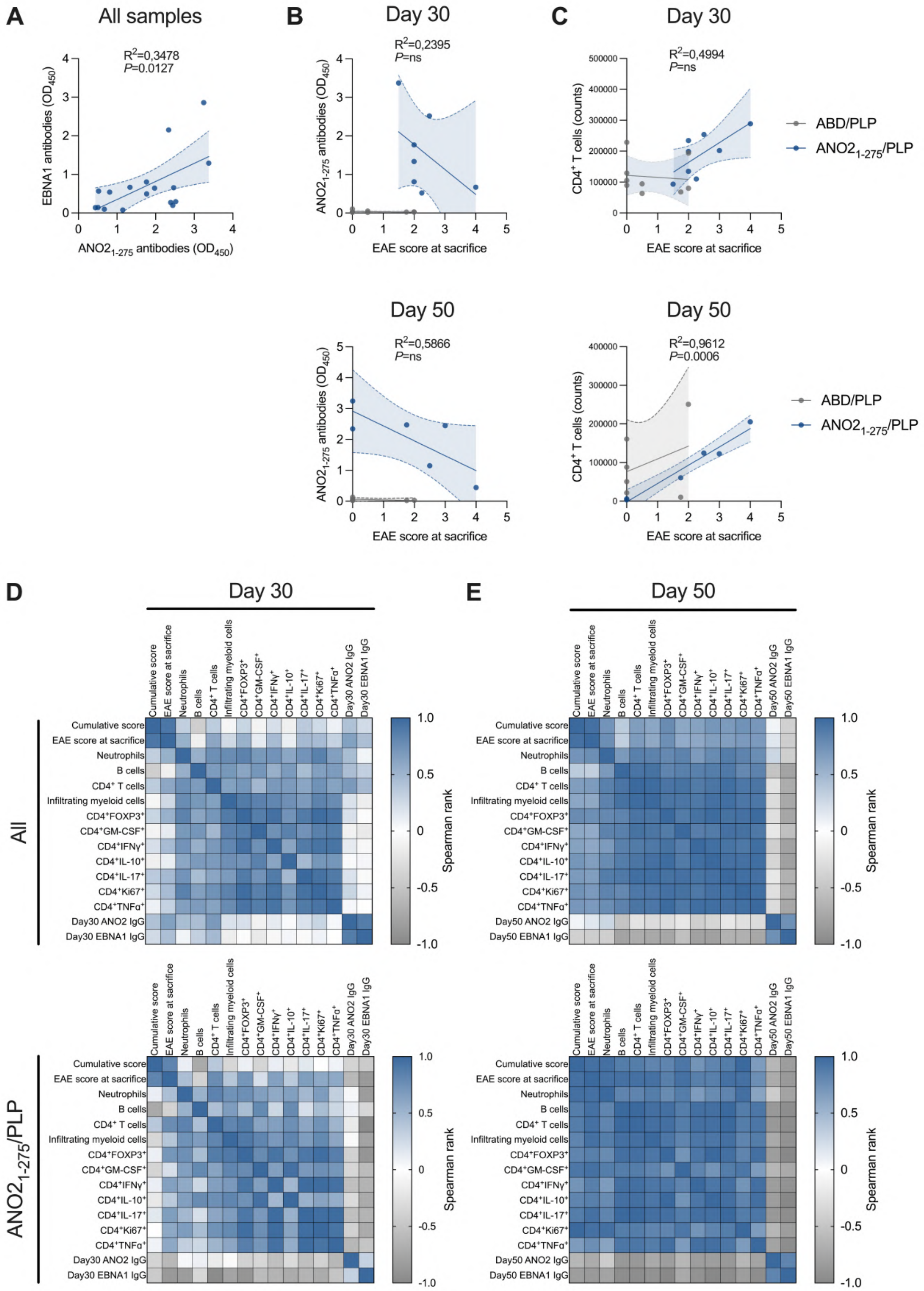
A
All samples
R2=0,3478
P=0.0127
EBNA1 antibodies (OD450)
ANO21-275 antibodies (OD450)
B
Day 30
R2=0,2395
P=ns
ANO21-275 antibodies (OD450)
EAE score at sacrifice
C
Day 30
R2=0,4994
P=ns
CD4+ T cells (counts)
EAE score at sacrifice
ABD/PLP
ANO21-275/PLP
Day 50
R2=0,5866
P=ns
ANO21-275 antibodies (OD450)
EAE score at sacrifice
Day 50
R2=0,9612
P=0.0006
CD4+ T cells (counts)
EAE score at sacrifice
ABD/PLP
ANO21-275/PLP
D
Day 30
E
Day 50
All
ANO21-275/PLP
Cumulative score
EAE score at sacrifice
Neutrophils
B cells
CD4+ T cells
Infiltrating myeloid cells
CD4+FOXP3+
CD4+GM-CSF+
CD4+IFNγ+
CD4+IL-10+
CD4+IL-17+
CD4+Ki67+
CD4+TNFα+
Day30 ANO2 IgG
Day30 EBNA1 IgG
Day50 ANO2 IgG
Day50 EBNA1 IgG
Spearman rank
1.0
0.5
0
-0.5
-1.0

**Figure S13. Correlation of CD4$^+$ T cell and antibody responses with EAE clinical scores.**

**A.** Correlation of anti-ANO2$_{1\text{-}275}$ and anti-EBNA1$_{380\text{-}641}$ IgG in ANO2$_{1\text{-}275}$/PLP-immunised mice from all timepoints. Two-tailed non-parametric Spearman correlation test.

**B.** Correlation of antibody responses to ANO2$_{1\text{-}275}$ at day 30 (top) and day 50 (bottom) with EAE score at sacrifice in ABD/PLP-immunised (grey) and ANO2$_{1\text{-}275}$/PLP-immunised mice (blue).

**C.** Correlation of CD4$^+$ T cells at day 30 (top) and day 50 (bottom) with EAE score at sacrifice in ABD/PLP-immunised (grey) and ANO2$_{1\text{-}275}$/PLP-immunised mice (blue). Lines represent the best-fit non-linear regression slope, and dashed lines represents the 95% CI. *P*-values were calculated using two-tailed non-parametric Spearman correlation tests and indicated where significant. Antibodies measured by ELISA (OD$_{450}$).

**D.** Multivariable analysis of all animals at day 30 (top left), all animals at day 50 (top right), ANO2$_{1\text{-}275}$/PLP-immunised mice only at day 30 (bottom left) and ANO2$_{1\text{-}275}$/PLP-immunised mice only at day 50 (bottom right).

*P*-values were calculated using two-tailed non-parametric Spearman correlation tests. Optical density (OD), confidence interval (CI). $^*P < 0.05$; $^{**}P < 0.01$; $^{***}P < 0.001$.

### Figure S14

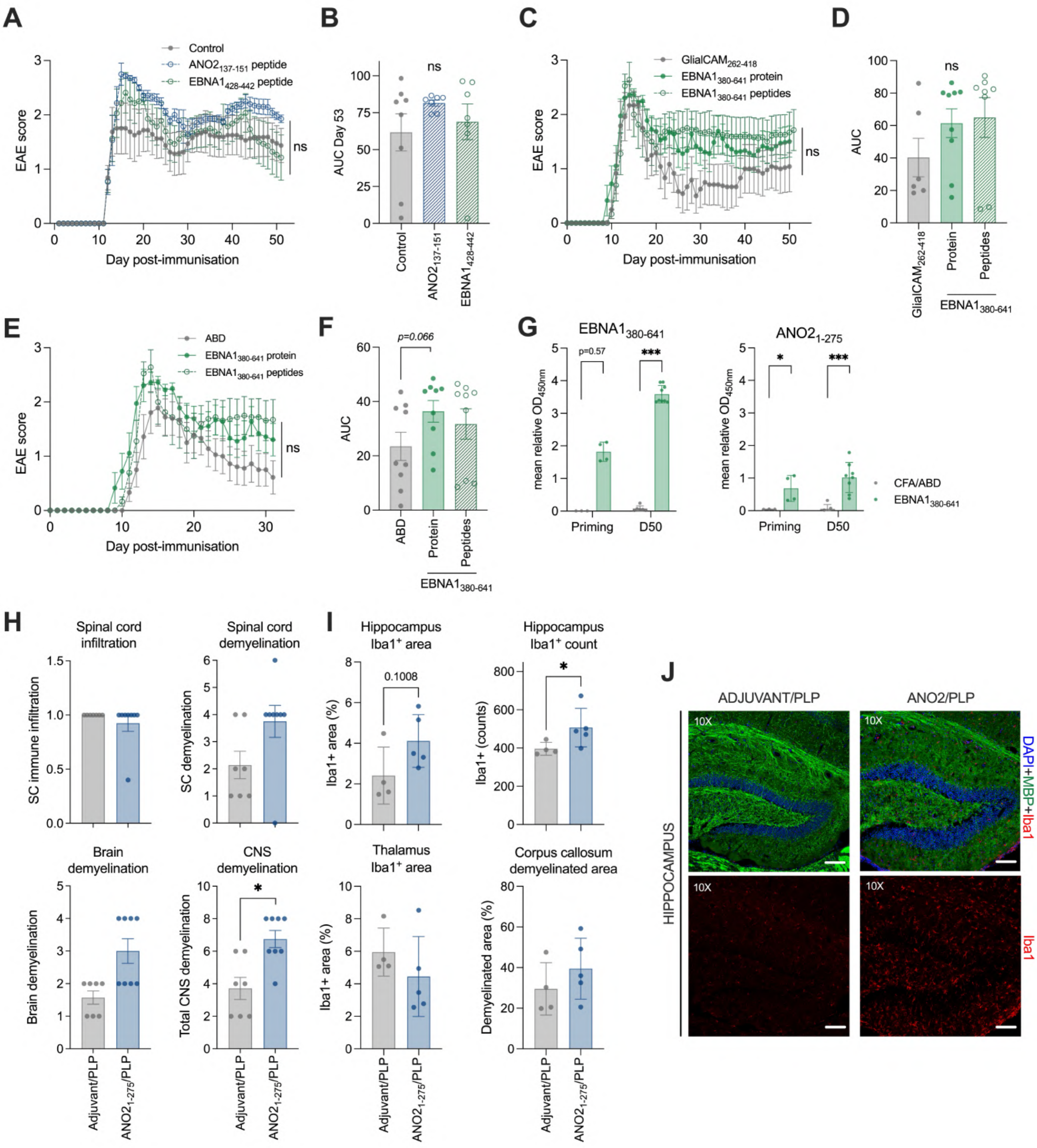

A
Control
ANO2 137-151 peptide
EBNA1 428-442 peptide
EAE score
Day post-immunisation
ns
B
AUC Day 53
ns
Control
ANO2 137-151
EBNA1 428-442
C
GlialCAM 262-418
EBNA1 380-641 protein
EBNA1 380-641 peptides
EAE score
Day post-immunisation
ns
D
AUC
ns
GlialCAM 262-418
Protein
Peptides
EBNA1 380-641
E
ABD
EBNA1 380-641 protein
EBNA1 380-641 peptides
EAE score
Day post-immunisation
ns
F
AUC
p=0.066
ABD
Protein
Peptides
EBNA1 380-641
G
EBNA1 380-641
ANO2 1-275
mean relative OD 450nm
p=0.57
Priming
D50
CFA/ABD
EBNA1 380-641
H
Spinal cord infiltration
SC immune infiltration
Spinal cord demyelination
SC demyelination
Brain demyelination
Brain demyelination
CNS demyelination
Total CNS demyelination
Adjuvant/PLP
ANO2 1-275/PLP
I
Hippocampus Iba1+ area
Iba1+ area (%)
0.1008
Hippocampus Iba1+ count
Iba1+ (counts)
Thalamus Iba1+ area
Iba1+ area (%)
Corpus callosum demyelinated area
Demyelinated area (%)
Adjuvant/PLP
ANO2 1-275/PLP
J
ADJUVANT/PLP
ANO2/PLP
HIPPOCAMPUS
10X
DAPI+MBP+Iba1
Iba1

**Figure S14 (continued)**

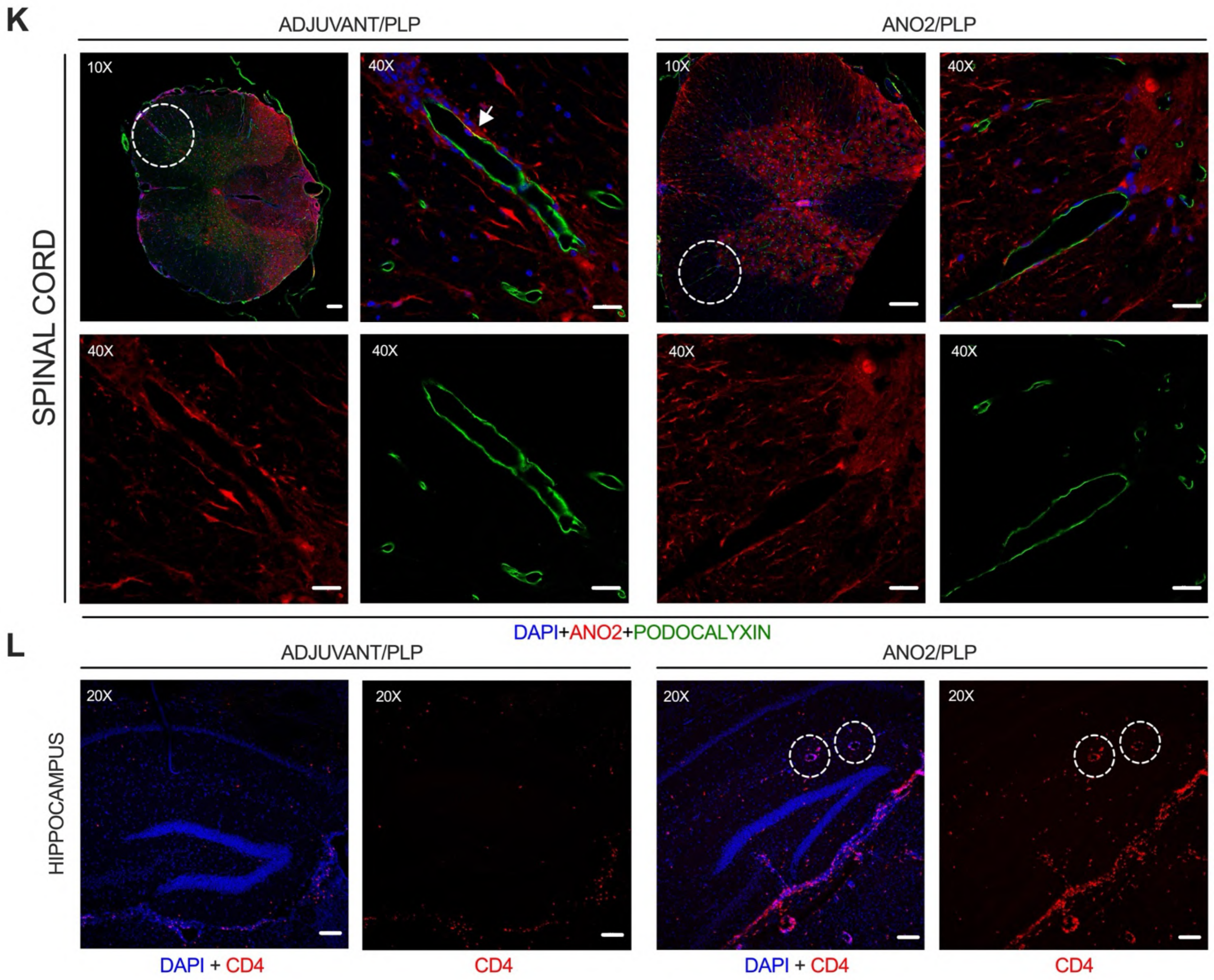


**Figure S14. EAE disease course of EBNA1, ANO2 and GlialCAM pre-immunised mice and immunofluorescent staining of aged ANO2 pre-immunised mice**

SJL/J mice were pre-immunised with ABD, $EBNA1_{380\text{-}641}$ or $GlialCAM_{262\text{-}418}$ proteins, an $EBNA1_{380\text{-}641}$ overlapping peptide pool, $EBNA1_{428\text{-}442}$ or $ANO2_{137\text{-}151}$ peptides plus CpG1826 and incomplete Freund's adjuvant (IFA). Three weeks later, mice were immunised with PLP plus complete Freund's adjuvant (CFA) and their disease course followed for up to 55 days (schematic in Figure 2B). Data only includes animals that got sick.

**A.** EAE scores of control (n=8), $ANO2_{137\text{-}151}$ peptide (n=8) and $EBNA1_{428\text{-}442}$ (n=8) peptide pre-immunised mice over 55 days.

**B.** Area under curve (AUC) of EAE scores at day 53 post EAE induction of control (n=8), $ANO2_{137\text{-}151}$ peptide (n=8) and $EBNA1_{428\text{-}442}$ (n=8) peptide pre-immunised mice. One way ANOVA statistical test.

**C.** EAE scores of $GlialCAM_{262\text{-}418}$ (n=6), $EBNA1_{380\text{-}641}$ protein (n=9) and $EBNA1_{380\text{-}641}$ (n=9) peptide pre-immunised mice over 55 days.

**D.** Area under curve (AUC) at day 52 post EAE induction. One way ANOVA.

**E.** EAE scores of ABD (n=9), $EBNA1_{380-641}$ protein (n=9) and $EBNA1_{380-641}$ (n=9) peptide pre-immunised mice over 55 days. One way ANOVA.

**F.** AUC at day 31 post-immunisation. One way ANOVA.

**G.** Antibody reactivity to $EBNA1_{380-641}$ (left) and $ANO2_{1-275}$ (right) in sera from ABD and $EBNA1_{380-641}$ protein pre-immunised mice at priming and at day 50 post-EAE induction by ELISA. Mann-Whitney test.

**H.** Spinal cord (SC) immune infiltration (top left), SC demyelination (top right), brain demyelination (bottom left) and total central nervous system (CNS) demyelination (bottom right) of mice immunised with adjuvant/PLP (n=8) or $ANO2_{1-275}$/PLP (n=9) as measured by immunofluorescence. Statistical test is the Mann-Whitney test.

**I.** Quantification of $Iba1^{+}$-infiltrating macrophages/microglia in the hippocampus (top left, top right), thalamus (bottom left) and demyelinated area of the corpus callosum (bottom right). Data presented as the $Iba1^{+}$ cell count or the % $Iba1^{+}$ area, Mann-Whitney test.

**J.** Representative immunofluorescent staining of adjuvant/PLP and $ANO2_{1-275}$/PLP-immunised mouse brains stained for DAPI (blue), MBP (green) and Iba1 (red), representative areas the hippocampus (10X). Merge: all channels overlayed.

**K.** Immunofluorescent staining of spinal cords from adjuvant/PLP and $ANO2_{1-275}$/PLP immunised mice for DAPI (blue), ANO2 (red) and podocalyxin (green), all channels overlayed. In the adjuvant/PLP group, ANO2 was expressed partially in endothelial cells in vessels (overlapped with podocalyxin), but more on the vascular basal membrane. In the ANO2/PLP group, vascular ANO2 expression seemed weaker and relatively absent on endothelial cells.

**L.** Immunofluorescent staining of brains from adjuvant/PLP and $ANO2_{1-275}$/PLP immunised mice for DAPI (blue) and CD4 (red). $CD4^{+}$ immune infiltration observed into the hippocampus and was localised around vessels.

For the immunofluorescence staining images, linear adjustments to brightness, contrast and saturation were made for visibility purposes. The changes were universally applied and consistent between the two different groups. The white lines indicate distance, 100µm for 10x and 20x images, and 20µm for 40x images.

Area under curve (AUC). *$P < 0.05$; **$P < 0.01$; ***$P < 0.001$.

**Figure S15**

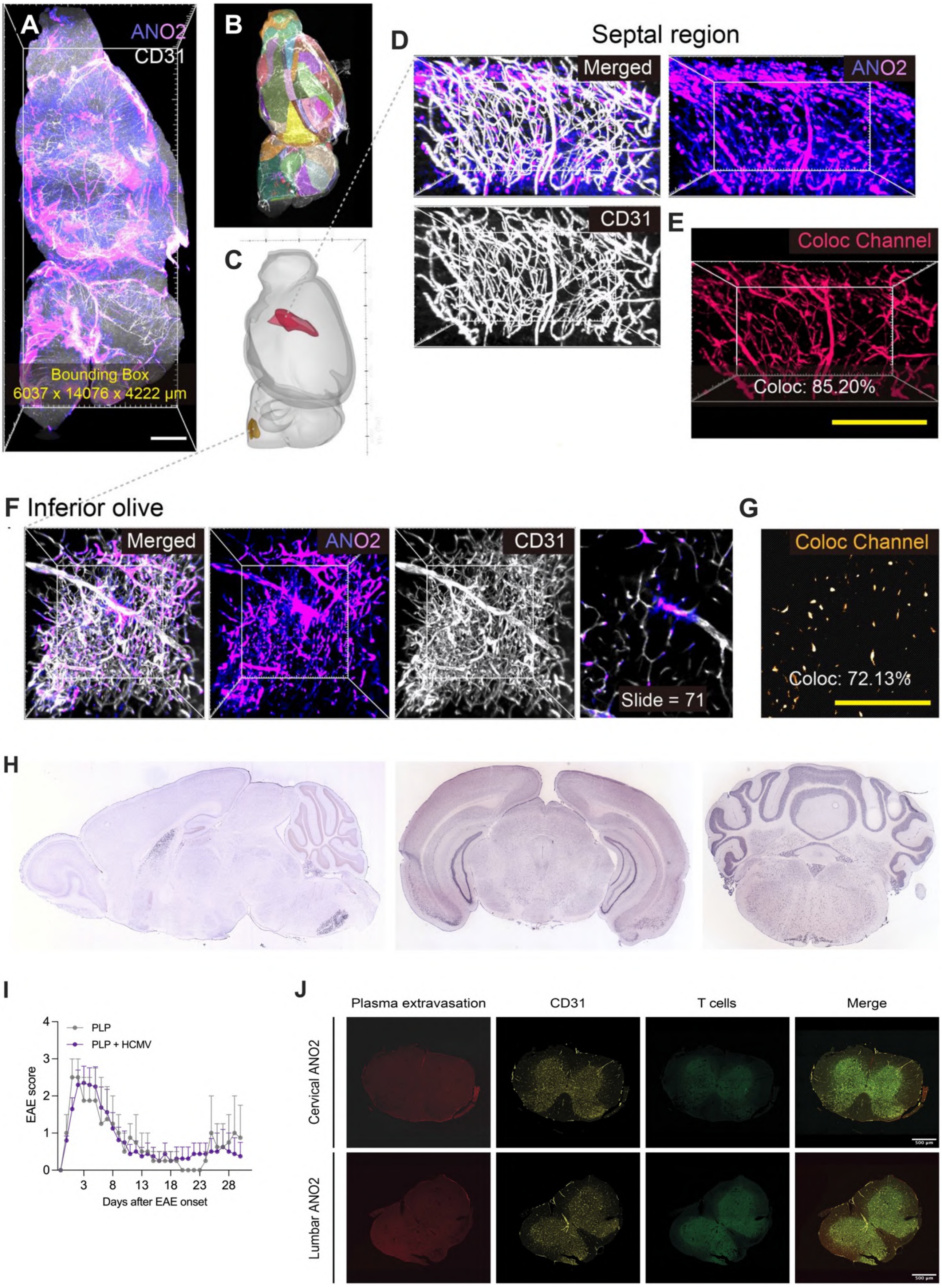
A
ANO2
CD31
Bounding Box
6037 x 14076 x 4222 μm
B
C
D
Septal region
Merged
ANO2
CD31
E
Coloc Channel
Coloc: 85.20%
F Inferior olive
Merged
ANO2
CD31
Slide = 71
G
Coloc Channel
Coloc: 72.13%
H
I
PLP
PLP + HCMV
EAE score
Days after EAE onset
J
Plasma extravasation
CD31
T cells
Merge
Cervical ANO2
Lumbar ANO2

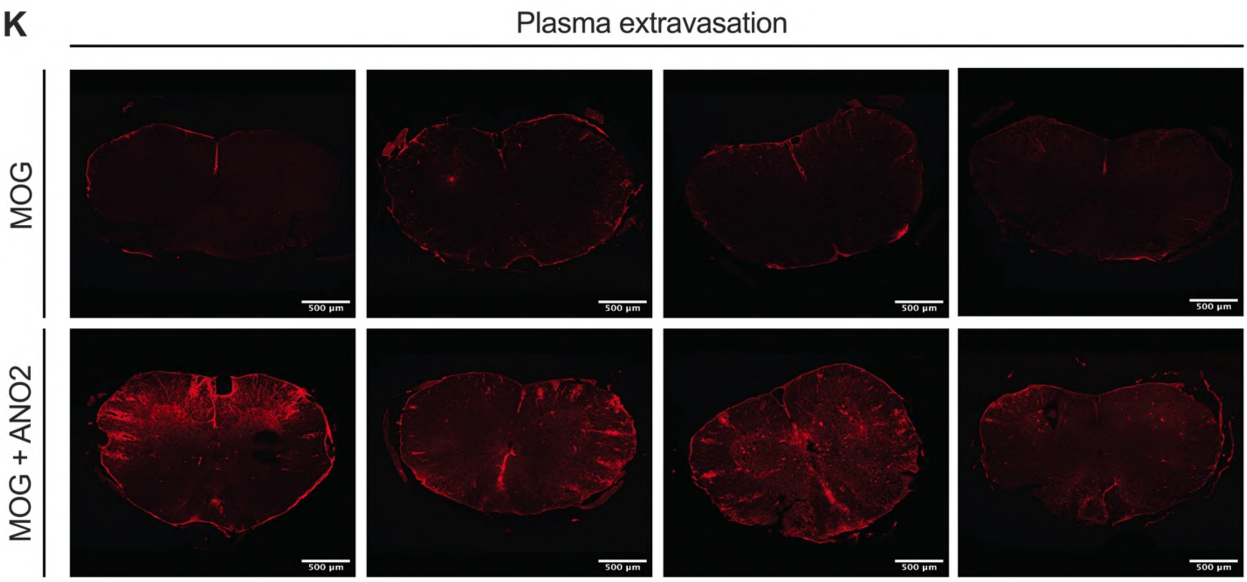
K
Plasma extravasation
MOG
MOG + ANO2
500 µm

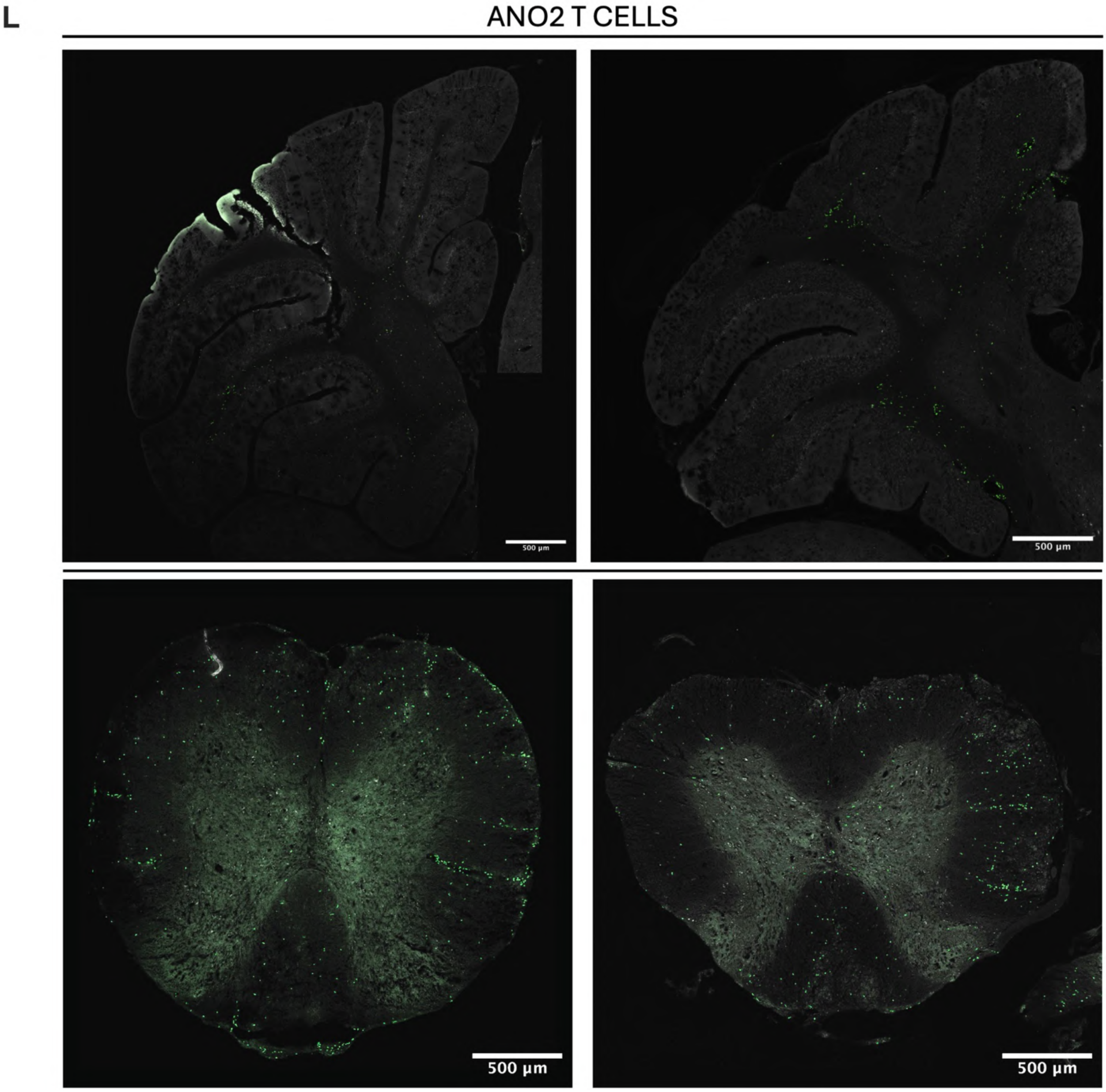
L
ANO2 T CELLS
500 µm

**Figure S15. Whole-mount visualisation and regional co-expression analysis of ANO2 and CD31 in the septum and inferiour olivary nucleus.**

**A.** Volumatic rendering of the representative half brain of an adult SJL/J mouse (6 months old). ANO2 signal is displayed using an intensity-based gradient from violet to pink (ULTRA colormap), and the vasculature marker CD31 is shown in white. Bounding Box: 6037 x 14076 x 4222 µm. Scale bar = 1000 µm, white.

**B.** The whole-brain dataset registered to the Allen Mouse Brain Atlas at 25 µm resolution.

**C.** Based on ANO2 and CD31 co-expression patterns, representative regions were selected for detailed analysis: the septal region (highlighted in crimson) and the inferior olive was selected for detailed analysis (highlighted in orange).

**D–G.** Representative co-expression images and colocalisation analysis of ANO2 and CD31 in the selected regions: d-e. isocortex and f-g. inferior olive. Scale bar = 250 µm, yellow.

**H.** Allen brain atlas RNA expression of ANO2 in mouse brains demonstrating staining in the inferior olivary nucleus, septal region and cerebellum. Sagittal view (left) and two coronal view (middle and right).

**I.** Immunofluorescent imaging of lumbar and cervical spinal cords of mice which received adoptive transfer of ANO2-specific T cells only (GFP+). Plasma extravasation (red), CD31 staining (classical endothelial marker, yellow) and T cells (GFP).

**J.** EAE scores of SJL/J mice adoptively transferred with PLP-specific T cells only or PLP+HCMV-specific T cells as described in Figure 4A.

**K.** Immunofluorescent imaging of plasma extravasation in the cervical spinal cords of mice adoptively transferred with MOG only or MOG+ANO2-specific T cells.

**L.** Immunofluorescent imaging of GFP+ ANO2-specific T cells located in the cerebellum (top) and cervical spinal cord (bottom) of mice which had received adoptive transfer of MOG+ANO2-specific T cells. For this experiment, GFP+ ANO2-specific T cells were generated from $GFP^{+}$ SJL/J x C57BL/6 F1 congenic mice, and were co-transferred with GFP-negative MOG-specific T cells generated from $GFP^{-}$ SJL/J x C57BL/6 F1 congenic.

**Figure S16**

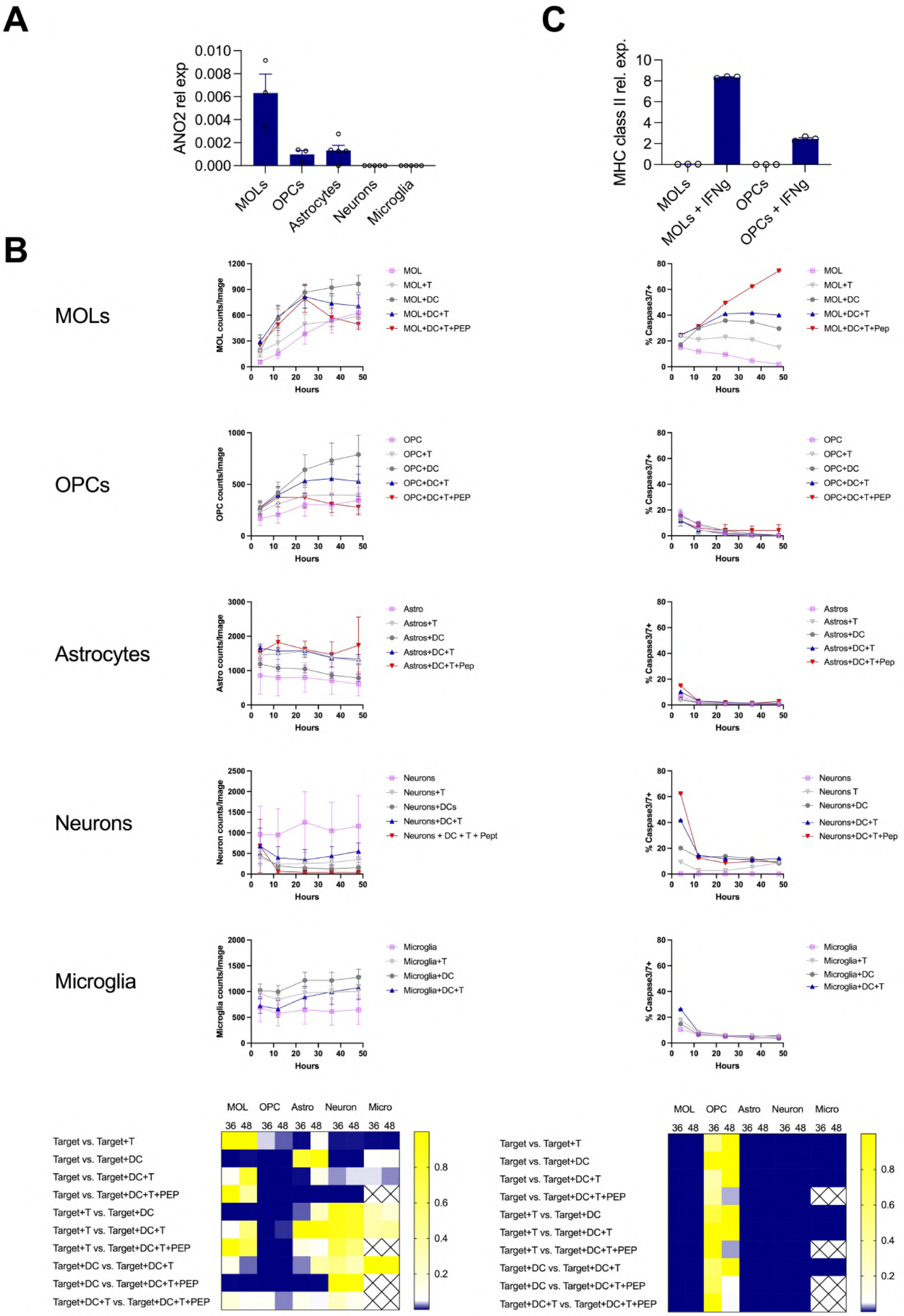
A
ANO2 rel exp
0.010
0.008
0.006
0.004
0.002
0.000
MOLs
OPCs
Astrocytes
Neurons
Microglia
C
MHC class II rel. exp.
10
8
6
4
2
0
MOLs
MOLs + IFNg
OPCs
OPCs + IFNg
B
MOLs
MOL counts/Image
Hours
MOL
MOL+T
MOL+DC
MOL+DC+T
MOL+DC+T+PEP
% Caspase3/7+
MOL+DC+T+Pep
OPCs
OPC counts/Image
OPC
OPC+T
OPC+DC
OPC+DC+T
OPC+DC+T+PEP
Astrocytes
Astro counts/Image
Astro
Astros+T
Astros+DC
Astros+DC+T
Astros+DC+T+Pep
Astros
Neurons
Neuron counts/Image
Neurons+T
Neurons+DCs
Neurons+DC+T
Neurons + DC + T + Pept
Neurons T
Neurons+DC
Neurons+DC+T+Pep
Microglia
Microglia counts/Image
Microglia+T
Microglia+DC
Microglia+DC+T
MOL OPC Astro Neuron Micro
36 48 36 48 36 48 36 48 36 48
Target vs. Target+T
Target vs. Target+DC
Target vs. Target+DC+T
Target vs. Target+DC+T+PEP
Target+T vs. Target+DC
Target+T vs. Target+DC+T
Target+T vs. Target+DC+T+PEP
Target+DC vs. Target+DC+T
Target+DC vs. Target+DC+T+PEP
Target+DC+T vs. Target+DC+T+PEP
0.8
0.6
0.4
0.2

**Figure S16. ANO2-specific T cells induce caspase3/7 activation in mature oligodendrocytes (MOLs) and oligodendrocyte precursor cells (OPCs) *in vitro***

MOLs, OPCs, astrocytes, neurons and microglia were sorted from either post-natal or adult SJL/J x C57BL/6-GFP animals, to allow for *in vitro* tracking, and were cultured with SJL/J-derived ANO2-specific resting polyclonal CD4+ T cells, bone marrow-derived LPS-activated dendritic cells, or a combination of both. A separate cell of wells with target cells, T cells and DCs were incubated with exogenous $ANO2_{1\text{-}275}$ peptide pool as a positive control for all conditions with the exception of microglia.

**A.** ANO2 expression in *in vitro* expanded primary glial cells (n = 3-5) as assessed by qPCR.

**B.** Incucyte tracking of cultures over a 48h period (n=5-6). Total numbers of cells/well (left panel) as well as percent Caspase3/7-positive cells out of the total (right panel) for every given timepoint. One-way ANOVA with Tukey's multiple comparison statistical test for all combinations within one glial cell type and timepoint. Timepoints: 36h and 48h for cell numbers; 4h and 48h for Caspase3/7 to account for early as well as late effects.

**C.** MHC class II expression in *in vitro* expanded primary MOLs and OPCs after 24h incubation with IFNγ (n = 3) as assessed by qPCR.

*$P$ < 0.05.

### Figure S17

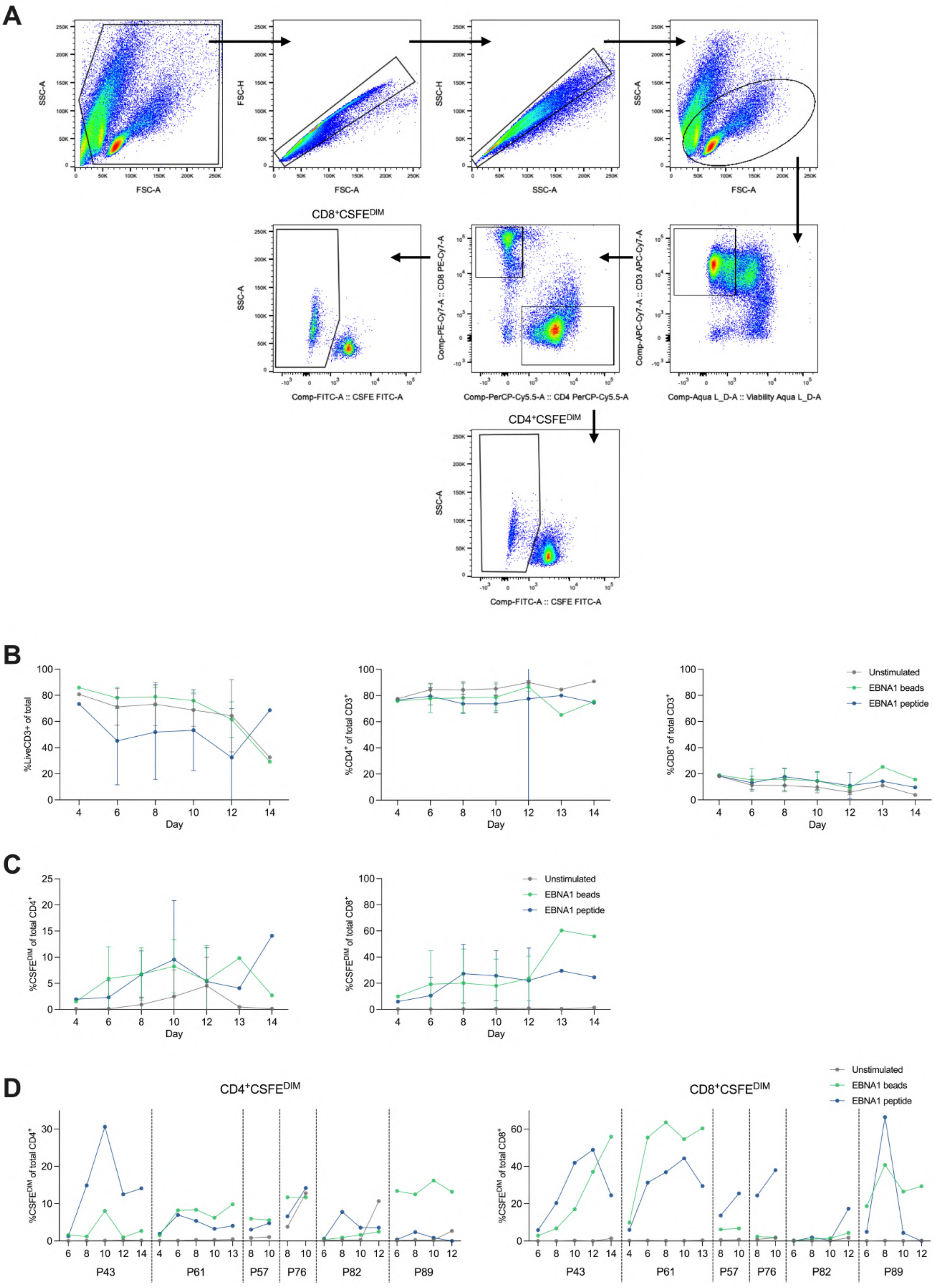
A
FSC-A
SSC-A
FSC-H
SSC-H
CD8+CSFEDIM
CD4+CSFEDIM
Comp-FITC-A :: CSFE FITC-A
Comp-PerCP-Cy5.5-A :: CD4 PerCP-Cy5.5-A
Comp-PE-Cy7-A :: CD8 PE-Cy7-A
Comp-Aqua L_D-A :: Viability Aqua L_D-A
Comp-APC-Cy7-A :: CD3 APC-Cy7-A
B
%LiveCD3+ of total
%CD4+ of total CD3+
%CD8+ of total CD3+
Day
Unstimulated
EBNA1 beads
EBNA1 peptide
C
%CSFEDIM of total CD4+
%CSFEDIM of total CD8+
D
CD4+CSFEDIM
CD8+CSFEDIM
P43
P61
P57
P76
P82
P89

**Figure S17. Optimisation of T cell expansion following antigen bead or peptide stimulation using EBNA1.**

Unstimulated, EBNA1 bead-stimulated or EBNA1 peptide-stimulated $CD45RA^-$ PBMC from MS-Nat donors were cultured for 4-14 days and surface stained to isolate the proliferating T cell population.

**A.** Gating strategy to isolate the $CD4^+CSFE^{DIM}$ and $CD8^+CSFE^{DIM}$ populations.

**B.** Percentage of $CD3^+$, $CD4^+$ and $CD8^+$ T cells in culture following stimulation.

**C.** Percentage of $CSFE^{DIM}$ $CD4^+$ and $CD8^+$ cells following stimulation.

**D.** Percentage of $CSFE^{DIM}CD4^+$ (left) and $CSFE^{DIM}CD8^+$ (right) T cells following stimulation for individual donors.

#### Figure S18

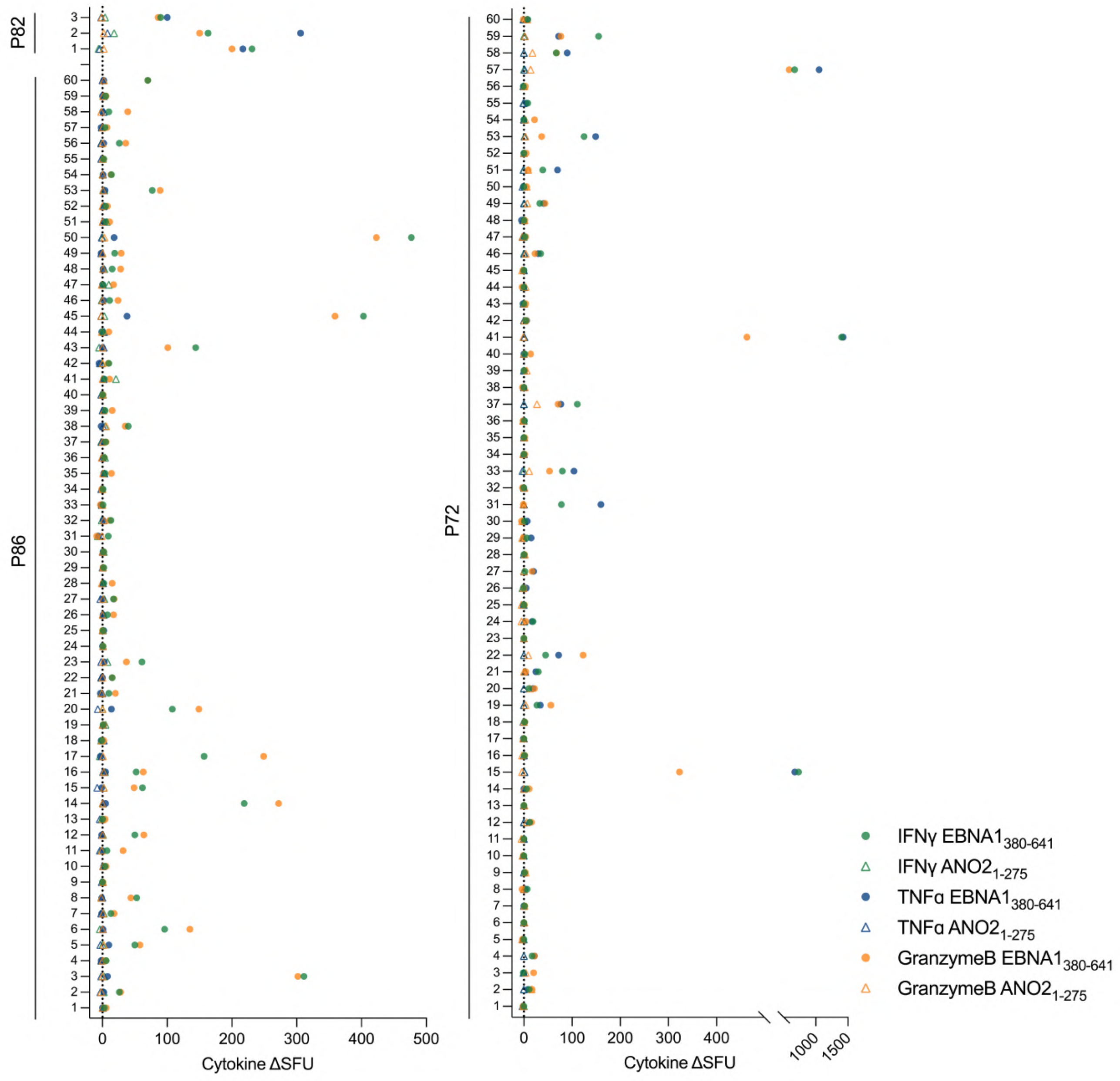


**Figure S18. EBNA1$_{380\text{-}641}$ bulk T cell lines screening for reactivity to EBNA1$_{380\text{-}641}$ and ANO2$_{1\text{-}275}$ by FluoroSpot.**

Proliferating LiveCD3$^{+}$CD4$^{+}$CSFE$^{DIM}$ T cells after 7-day stimulation with EBNA1$_{380\text{-}641}$ beads from three representative donors (P72, P82 and P86) were sorted at 100 cells/well and expanded for a further 2 weeks with PHA, IL-2 and irradiated allogeneic feeder PBMC. After 2 weeks, IL-2 was withdrawn and bulk T cell lines were screened for reactivity to ANO2$_{1\text{-}275}$ and EBNA1$_{380\text{-}641}$ beads by IFNγ/TNFα/GranzymeB FluoroSpot. T cell bulk lines showed a range of specificities and cytokine production although many responded to EBNA1$_{380\text{-}641}$.

#### Figure S19

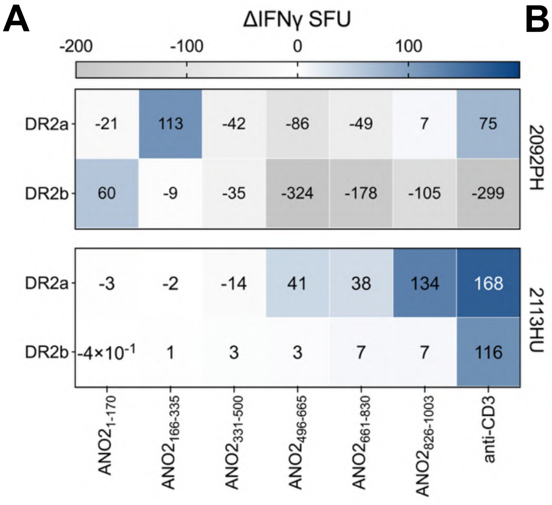


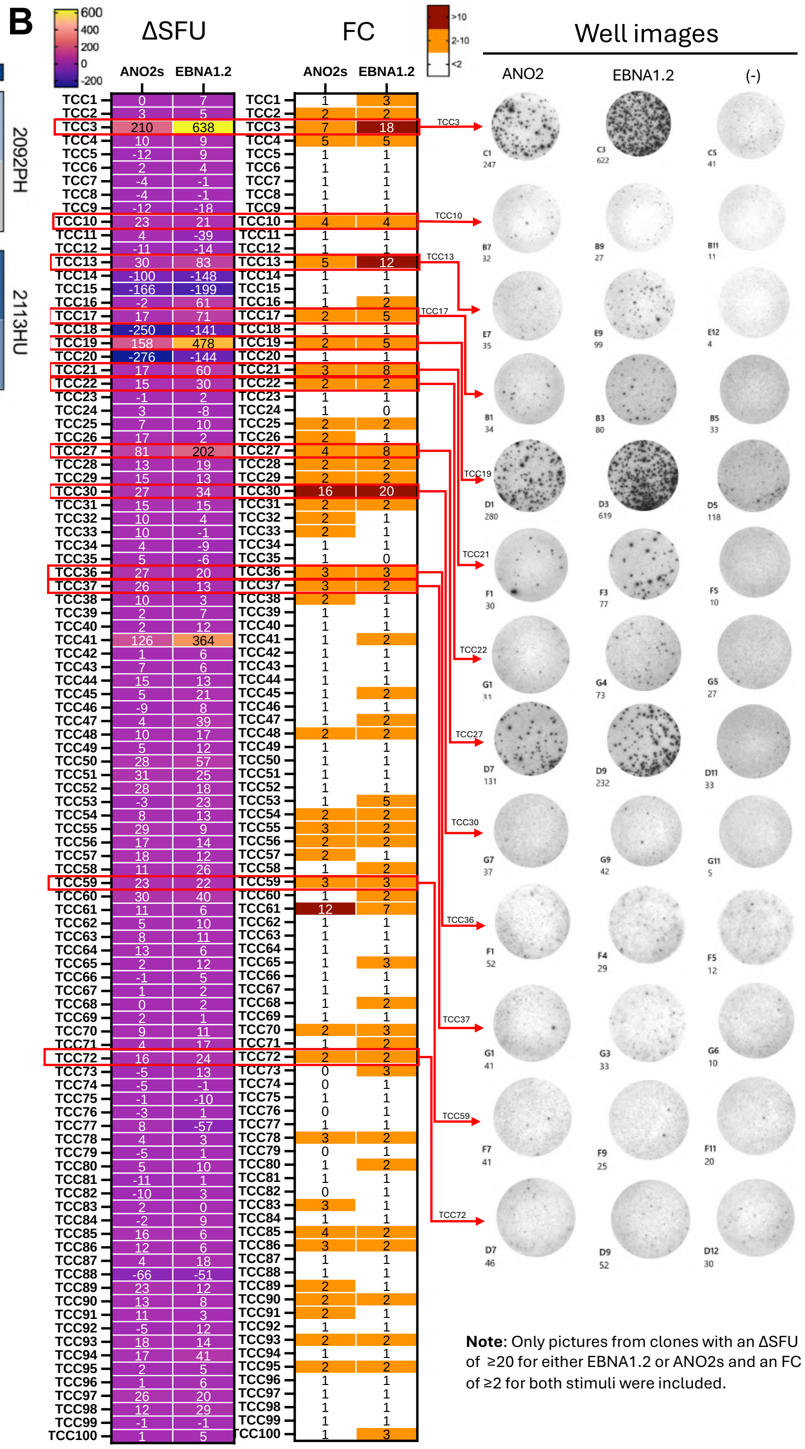


**Figure S19. EBNA1$_{380\text{-}641}$-stimulated T cell bulk and clones show reactivity to EBNA1$_{380\text{-}641}$ and ANO2$_{79\text{-}168}$ by FluoroSpot.**

**A.** T cell bulk (TCB) lines from two untreated HLA-DRB1*15:01$^{+}$ pwMS (top panel: donor 2092PH and bottom panel: donor 2113HU) were generated from CFSE$^{DIM}$ (proliferating, responding) and CFSE$^{HI}$ (non-proliferating, non-responding) sorted after EBNA1$_{380\text{-}641}$ stimulation were screened for ANO2 reactivity by IFNγ ELISpot using overlapping 20-mer peptide pools covering the entire sequence of ANO2 (aa1-1003). Anti-CD3 beads were used as a positive control and BLS cells expressing DR2a and DR2b were used as antigen presenting cells. Data is presented as background subtracted ΔIFNγ SFU using the BLS DR2a and BLS DR2b no peptide control conditions.

**B.** T cell clones (TCC) generated from an *HLA-DRB1*15:01$^{+}$* MS donor after stimulating with EBNA1$_{380\text{-}641}$ and sorting the proliferating LiveCD3$^{+}$CD4$^{+}$CSFE$^{DIM}$ compartment were screened for reactivity to ANO2$_{79\text{-}168}$ and EBNA1$_{380\text{-}641}$ beads by IFNγ ELISpot. TCC were incubated with autologous PBMC and stimulated with anti-CD3 and antigen coupled beads: negative control, ANO2$_{79\text{-}641}$ and EBNA1$_{380\text{-}641}$. TCC showed a range of reactivities, with some isolated clones responding to both EBNA1$_{380\text{-}641}$ and ANO2$_{79\text{-}168}$. Data presented as the ΔIFNγ SFU by subtracting spots in negative control (NC) wells (left) and also the fold change from the negative control condition (right).

ANO2 short aa79-168 (ANO2s), spot forming units (SFU), fold change (FC), negative control (-).

#### Figure S20

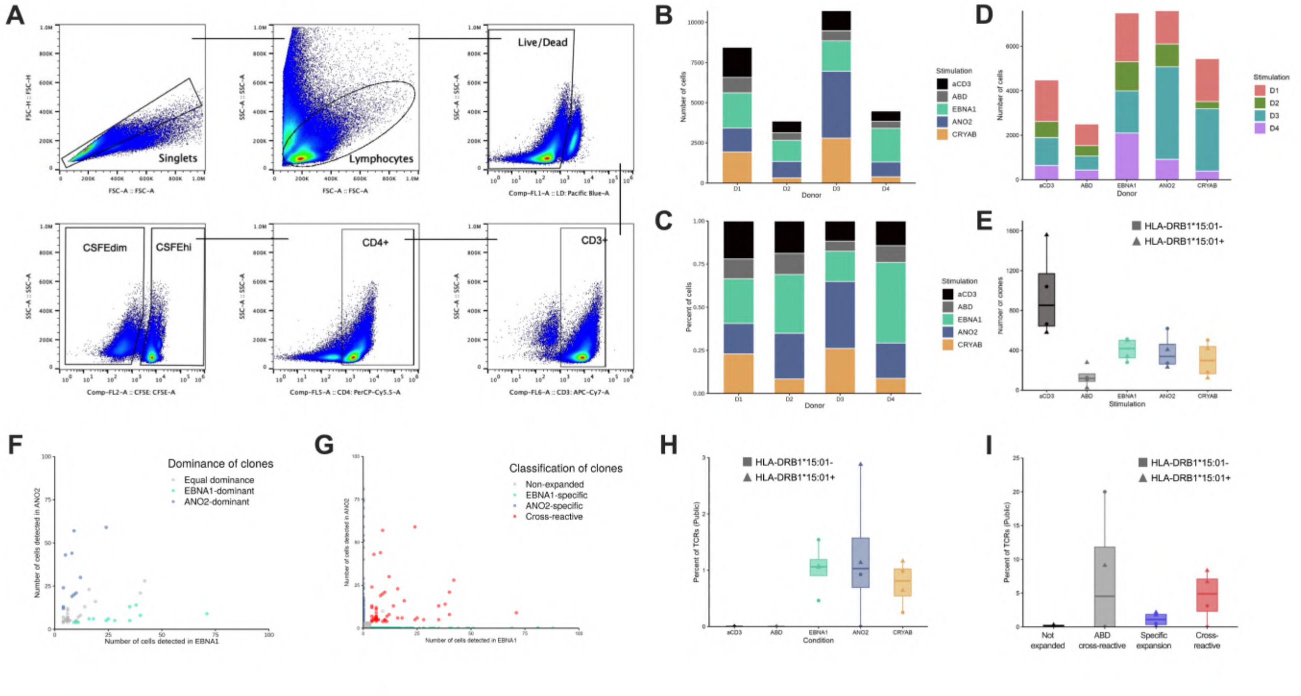


**Figure S20. Analysis of responding TCR repertoires in single cell sequencing data.**

**A.** Gating strategy for identification and sorting of responding LiveCD3$^+$CD4$^+$CSFE$^{DIM}$ T cells after 8-day stimulation with antiCD3, ABD, EBNA1$_{380-641}$, ANO2$_{1-275}$ or CRYAB$_{FL}$ for single cell sequencing.

**B.** Number of cells from each donor (D1-4) and condition (anti-CD3, ABD, EBNA1$_{380-641}$, ANO2$_{1-275}$ and CRYAB$_{FL}$) in single cell sequencing after quality control.

**C.** Percent of cells in each stimulation from each donor (D1-4).

**D.** Total number of cells in each stimulation (separated by donor) in single cell sequencing after quality control.

**E.** Total number of expanded TCR clones per stimulation across donors.

**F.** Comparison of TCR copy numbers of expanded cells found in both EBNA1 and ANO2 expansions from all four donors. Green dots represent TCRs which have a higher copy number in the EBNA1 stimulation, blue dots are TCRs which have a higher copy number in ANO2 and grey dots are TCRs which have an equal copy number in both stimulations.

**G.** Number of copy numbers of TCRs which are only found in ANO2, only found in EBNA1, are found in both EBNA1 and ANO2 stimulations (cross-reactive) and are non-expanded across all four donors.

**H.** Percent of TCRs per condition that are public (shared between multiple donors).

**I.** Percent of non-expanded, ABD cross-reactive, specifically expanded or cross-reactive TCRs which are public (shared between donors).

Albumin binding domain (ABD), T cell receptor (TCR). Squares indicate *HLA-DRB1*15:01* non-carriers, and triangles represent *HLA-DRB1*15:01* carriers for the whole figure.

### Figure S21

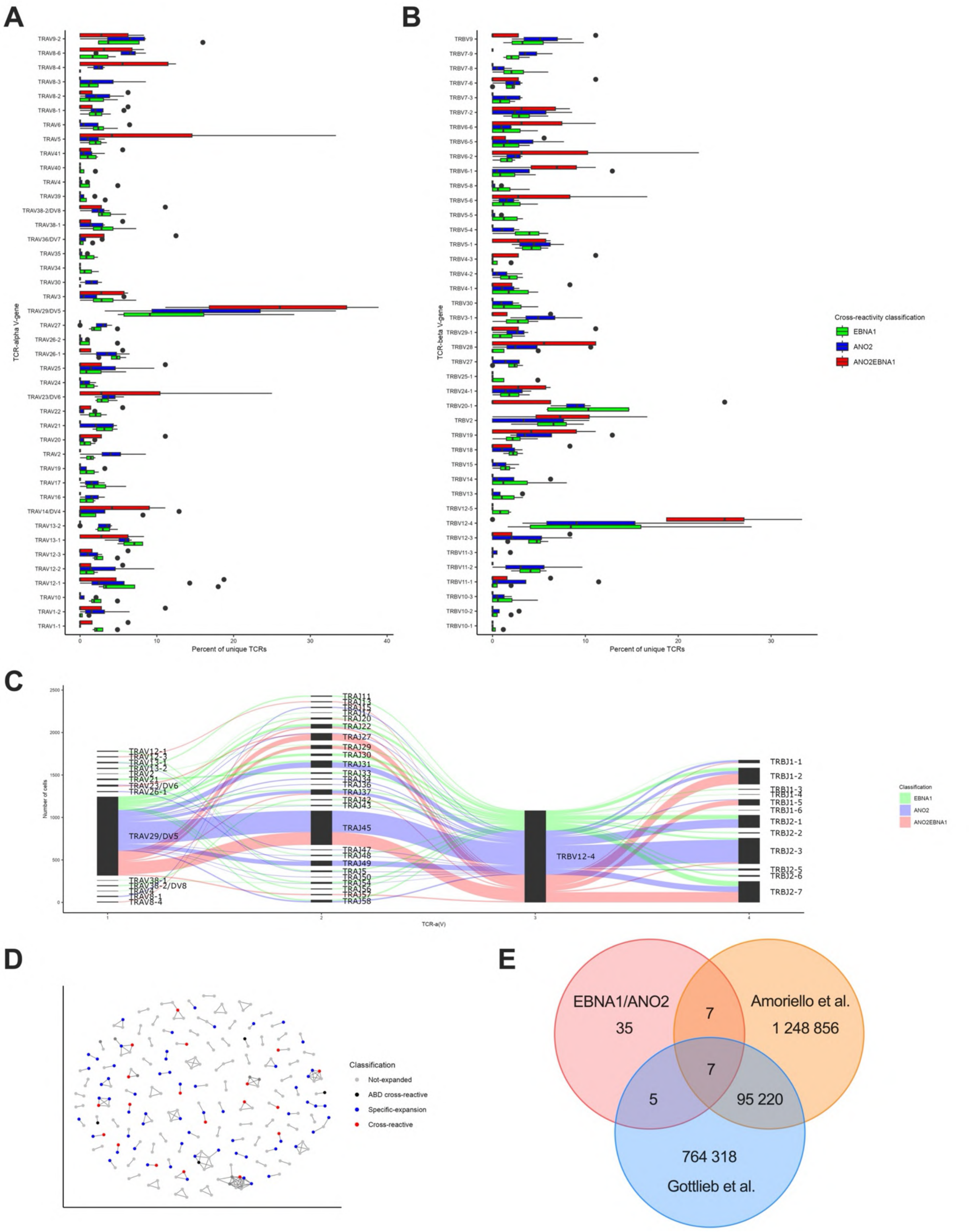


**Figure S21. Characteristics of TCRs from antigen-specific stimulations**

**A.** TCRα gene usage of expanded TCRs which are ANO2-specific (blue), EBNA1-specific (green) and EBNA1/ANO2 cross-reactive TCRs (red).

**B.** TCRβ gene usage of expanded TCRs which are ANO2-specific (blue), EBNA1-specific (green) and EBNA1/ANO2 cross-reactive TCRs (red).

**C.** Sankey plot showing TCR gene usage and linkage of expanded ANO2-specific (blue), EBNA1-specific (green) and EBNA1/ANO2 cross-reactive TCRs (red) which express the **TRBV12-4** gene across all four donors, demonstrating the diversity of TCR gene usage within this population.

**D.** Network plot of clusters of TCRs based on amino acid similarity. Only clusters with greater than 2 nodes were included for visualisation purposes. Cross-reactive clones do not show clear clustering, suggesting diversity of TCR repertoire.

**E.** Overlap of expanded EBNA1/ANO2 cross-reactive TCRVβ CDR3 sequences with previous published data sets from the blood and CSF of MS patients and controls. In total, 19 expanded EBNA1/ANO2 cross-reactive TCRs were found to overlap with previously published datasets: 7 TCRs with Amoriello *et al.*, 5 with Gottlieb *et al.*, and 7 TCRs with both datasets.

T cell receptor (TCR).

### Figure S22

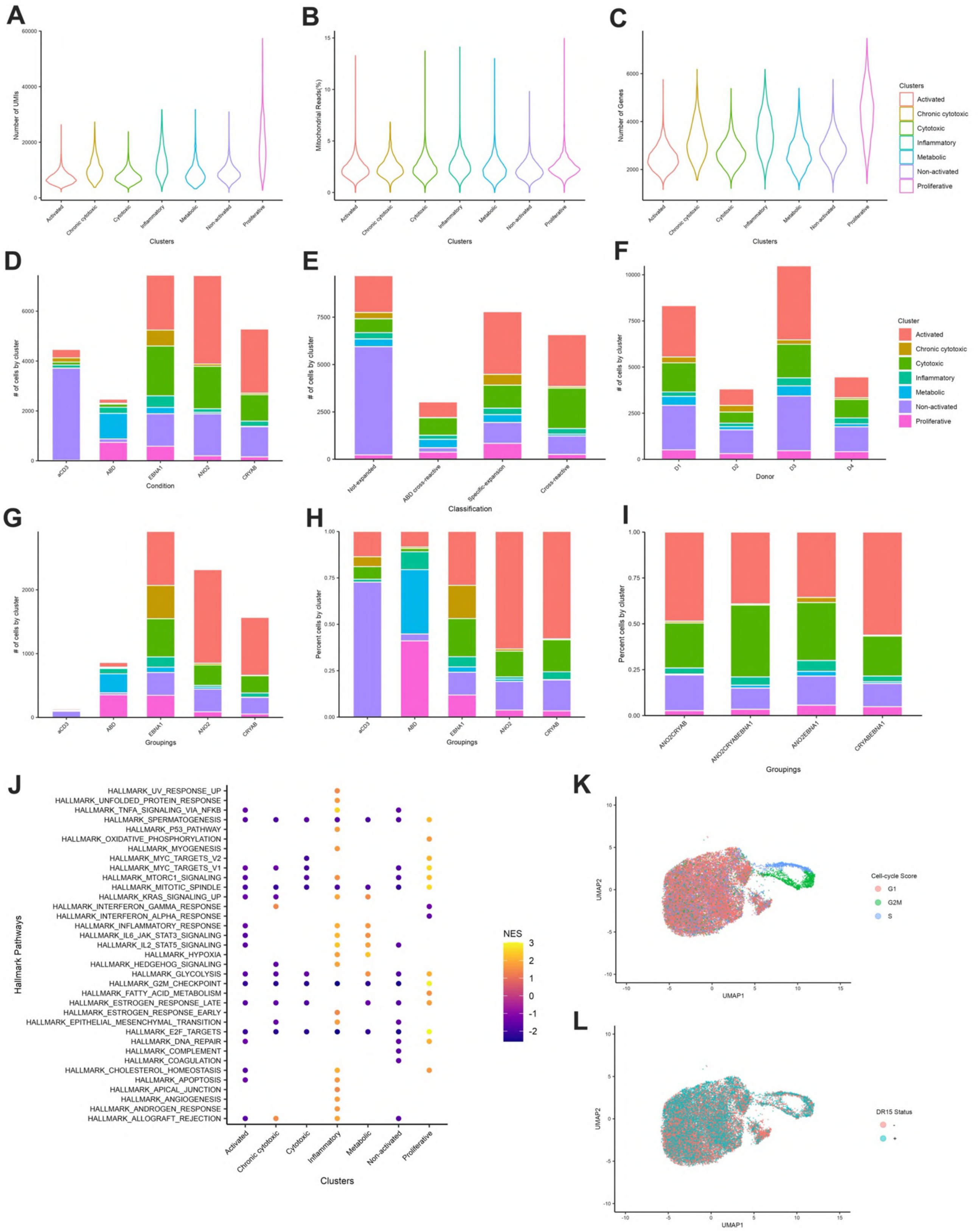


**Figure S22. Quality control and further analysis of single cell transcriptomes of antigen-specific T cells**

Data from Figure 7.

**A.** Total number of unique molecular identifier (UMIs) reads per cluster.

**B.** Frequency of mitochondrial reads per cluster (percent of total cells).

**C.** Total number of genes detected per cluster during single cell transcriptome analysis.

**D.** Total number of cells in each original stimulation condition (anti-CD3, ABD, EBNA1, ANO2 or CRYAB) overlayed with cluster identity to visualise the contribution of each cluster to conditions.

**E.** Total number of cells in each classification (non-expanded, ABD cross-reactive, specifically expanded or cross-reactive) overlayed with cluster identity.

**F.** Total number of cells from each donor (donors 1-4) overlayed with cluster identity.

**G.** Total number of specifically expanded cells (only found in one condition, **not** shared with any other stimulations) from each stimulation overlayed with cluster identity.

**H.** Frequency of specifically expanded cells (only found in one condition, **not** shared with any other stimulations) from each stimulation overlayed with cluster identity. Data presented as percent of total cells in a given stimulation, and is the same data as in Figure S22G.

**I.** Frequency of cells in each cluster overlayed onto cross-reactivity classifications: ANO2CRYAB, ANO2CRYABEBNA1, **ANO2EBNA1** or CRYABEBNA1. Data presented as percent of total cells in each cross-reactive classification.

**J.** Hallmark pathway analysis of clusters.

**K**. UMAP dimensionality reduction of T cells with cell cycle stage highlighted.

**L.** UMAP dimensionality reduction of T cells with HLA-DRB1*15:01 carrier and non-carrier status highlighted.
